# Functional connectome harmonics and dynamic connectivity maps of the preadolescent brain

**DOI:** 10.64898/2026.03.02.708791

**Authors:** Isabella L.C. Mariani Wigley, Aurora Berto, Ilkka Suuronen, Ashmeet Jolly, Ru Li, Harri Merisaari, Elmo P. Pulli, Aylin Rosberg, Hilyatushalihah K. Audah, Aaron Barron, Silja Luotonen, Massimiliano Pastore, Mattia Veronese, Hasse Karlsson, Riikka Korja, Linnea Karlsson, Joana Ribeiro Barbosa Cabral, Morten L. Kringelbach, Selen Atasoy, Jakob Seidlitz, Richard AI Betlehem, Jetro J. Tuulari

**Affiliations:** FinnBrain Birth Cohort Study, Turku Brain and Mind Center, Department of Clinical Medicine, University of Turku, Turku, Finland; Centre for Population Health Research, Turku University Hospital and University of Turku, Turku, Finland; Department of Psychiatry, University of Turku and Turku University Hospital, Turku, Finland; Department of Information Engineering, University of Padua, Italy; Department Psychology and Speech-Language Pathology, University of Turku, Turku, Finland; Department of Teacher Education, University of Turku, Turku, Finland; Department of Psychology and Behavioral Sciences, Aarhus University, Aarhus, Denmark; Turku Brain and Mind Center, University of Turku, Turku, Finland; Centre of Excellence in Learning Dynamics and Intervention Research (InterLearn), University of Jyväskylä and University of Turku, Finland; Department of Pediatric Neurology, Turku University Hospital, Turku, Finland; Department of Developmental and Social Psychology, University of Padua, Padua, Italy; Department of Neuroimaging; King’s College London, UK; Department of Public Health, University of Turku and Turku University Hospital; Department of Child Psychiatry, Turku University Hospital and University of Turku; Institute for Systems and Robotics (ISR-Lisboa), Instituto Superior Técnico, University of Lisbon, 1049-001 Lisboa, Portugal; Department of Bioengineering, Instituto Superior Técnico, University of Lisbon, 1049-001 Lisboa, Portugal; Centre for Eudaimonia and Human Flourishing, Linacre College, University of Oxford, Oxford, UK; Centre for Music in the Brain, Department of Clinical Medicine, Aarhus University, Aarhus, Denmark; Department of Psychiatry, University of Oxford, Oxford, UK; Lifespan Brain Institute, The Children’s Hospital of Philadelphia and Penn Medicine, Philadelphia, PA, 19104 USA; Department of Psychiatry, University of Pennsylvania, Philadelphia, PA, 19104 USA; Department of Child and Adolescent Psychiatry and Behavioral Science, The Children’s Hospital of Philadelphia, Philadelphia, PA, 19104 USA; Institute for Translational Medicine and Therapeutics, University of Pennsylvania, Philadelphia, PA, 19104 USA; Department of Psychology, University of Cambridge, Cambridge, United Kingdom; Neurocenter, Turku University Hospital, Turku, Finland

**Keywords:** Preadolescence, Brain, Resting-state fMRI, ABCD

## Abstract

The maturation of large-scale functional brain networks during preadolescence underpins critical cognitive and behavioral development. However, the spatial and temporal organization of these networks in this age group remains incompletely characterized. Here we applied Functional Connectome Harmonics (FCH) and Leading Eigenvector Dynamics Analysis (LEiDA) to resting-state fMRI data from over 11,000 children aged 9–10 years in the ABCD Study. FCH revealed hierarchical spatial gradients spanning cortical and subcortical regions, while LEiDA identified recurrent dynamic brain states aligned with canonical intrinsic connectivity networks. Linking these spatial and temporal components, we established a low-dimensional harmonic scaffold constraining brain dynamics during this developmental window. These findings provide a large-scale spatiotemporal reference framework of preadolescent functional brain organization, offering a foundation for characterizing neurodevelopmental benchmarks and early neural markers relevant to adolescent mental health.

## Introduction

Spontaneous brain activity measured with resting-state functional MRI exhibits low-frequency fluctuations that organize at the macroscale, giving rise to coherent patterns of co-activation across distributed brain areas known as resting-state networks (Sporns et al., 2021). During preadolescence, these networks undergo substantial maturation through both structural refinement and functional reorganization (Sun et al., 2025; Vijayakumar et al., 2021). These developmental processes reshape large-scale functional interactions, underscoring the need to characterize how brain network organization evolves during this critical period.

Preadolescence represents a sensitive neurodevelopmental window in which rapid neurobiological maturation coincides with the emergence of increasingly complex cognitive, social, and emotional demands (Andersen, 2003). During this period, variability in biological maturation (e.g., pubertal stage and body composition) and demographic factors such as sex and ethnicity may contribute to heterogeneity in large-scale brain organization (Dahl, 2004). As large-scale circuits undergo refinement through synaptic reorganization and white-matter maturation, preadolescents typically navigate expanding peer relationships, greater autonomy, and heightened socioemotional expectations (Andrews et al., 2021). The neurobehavioral phenotype emerging from the interplay of these neural and psychosocial processes has lasting consequences. For instance, mental health status at ages 9–10 strongly predicts adolescent psychopathology, and subclinical symptoms at this stage can forecast the onset of mental disorders only a few years later (Visser et al., 2000; Hou et al., 2025). Yet, the large-scale spatial and dynamic organization of functional networks during this highly relevant developmental window remains poorly understood. Elucidating how functional network architecture supports adaptation to these developmental demands is therefore crucial for identifying early mechanisms that shape adolescent mental health trajectories. However, it remains unclear how spatial frameworks - such as those capturing gradients of functional connectivity - and temporal frameworks - such as those describing dynamic network states-relate to one another in preadolescence, and whether they capture developmentally relevant features of brain organization during this period.

Computational approaches in neuroscience provide mathematical frameworks capable of capturing complementary aspects of large-scale brain organization. Among these, Functional Connectome Harmonics (FCH) decompose functional connectivity into frequency-ordered harmonic modes that characterize hierarchical spatial organization across cortical scales (Atasoy et al., 2016; Atasoy et al., 2018; Glomb et al., 2021), whereas Leading Eigenvector Dynamics Analysis (LEiDA) identifies dominant patterns of time-resolved network coupling that reveal recurring dynamic brain states (Cabral et al., 2017). These methods offer powerful tools for quantifying how spatial and temporal features of functional architecture emerge and interact during preadolescence. Extending this framework, FCH provides a compact representation of the brain’s spatial architecture by linking fine-grained topography with macroscopic gradients and canonical functional networks (Atasoy et al., 2016; Glomb et al., 2021). We have recently expanded the range of brain measures derived from FCHs in neonates; here, we extend this framework for the first time to characterize preadolescent brain organization.

By decomposing functional connectivity into natural harmonic modes, analogous to standing waves in physical systems, FCH captures intrinsic spatial patterns of synchronous activity that are coded in the functional connectivity of the human brain (Atasoy et al., 2016; Glomb et al., 2021). In this framework, each harmonic component is characterized by its energy, the frequency-weighted contribution of that harmonic to cortical dynamics, and its power, which reflects the instantaneous strength of activation of the harmonic at a given time point.

Together, these measures quantify the contribution of each spatial mode to functional architecture. Complementing this spatial perspective, LEiDA identifies moment-to-moment fluctuations in network coupling, detecting transient configurations in which brain regions become synchronized or desynchronized and quantifying their temporal occurrence to reveal the repertoire of dynamic brain states. From these states, LEiDA further quantifies their probability of occurrence (i.e., the proportion of time spent in each state), indexing its dominance, the lifetime (i.e., the mean duration a state persists once entered), and the transitions between states, which capture the flexibility of network dynamics. These coupling modes correspond closely to intrinsic connectivity networks and capture inter-individual variability relevant to cognitive, affective, and clinical phenotypes (Vohryzek et al., 2020; Lord et al., 2019; Figueroa et al., 2019; Mariani Wigley et al., 2026).

Here, we applied FCH and LEiDA to resting-state fMRI data from the multisite Adolescent Brain Cognitive Development (ABCD) baseline cohort, comprising more than 11,000 children aged 9–10 years and collected across 21 imaging sites using standardized acquisition protocols. We validated FCH patterns against established cortical gradients (e.g., Margulies et al., 2016) and confirmed that LEiDA-derived states align with canonical resting-state networks (Yeo et al., 2011). We then tested whether harmonic modes derived from FCH could predict LEiDA states across participants, thereby linking the spatial patterns captured by FCH with the temporal dynamics identified by LEiDA. Critically, we further examined whether connectomic metrics derived from both FCH (power and energy) and LEiDA dynamic state properties capture meaningful demographic and developmental variability.

This approach allowed us to assess the sensitivity of spatial and temporal connectomic frameworks to inter-individual differences during preadolescence. Together, these analyses provide the first large-scale characterization of functional connectome harmonics and dynamic coupling modes in preadolescence, establishing a reference framework for studying maturational processes and their relevance for emerging behavioral and mental health trajectories.

## Results

### Participants

Figure 1 outlines the steps used to derive the analytical sample from the baseline ABCD cohort (N = 11,868). For participants with multiple resting-state fMRI acquisitions, only the first scan was included in the primary preprocessing and subsequent analyses. After preprocessing and quality control, 8,536 participants met imaging-motion criteria (average framewise displacement, FD < 0.4), of which 6,624, matching the right number of ROIs and timepoints, were retained for connectomics analyses. Following exclusion of cases with missing data on key demographic (i.e., sex, ethnicity) and developmental variables (i.e., interview age, PDS, TMI), the final analytical sample (N = 4,453) was used for all statistical analyses unless otherwise specified. To assess potential selection bias, these individuals were compared with a subset of excluded subjects with available data on the variables of interest (N = 7,058), with respect to demographic variables (sex and ethnicity), developmental variables (interview age, pubertal development scale (PDS), and triponderal mass index (TMI)), and site of data-acquisition. PDS was derived from parent- and youth-reported puberty measures (Petersen et al., 1988) while TMI was calculated as weight (kg)/height (m)³ (Li et al., 2025). Included and excluded participants were broadly comparable across demographic variables (Table 1). No differences were observed for age or TMI. Distributions of these variables are shown in Supplementary Material Section 1.1.

**Figure 1.**
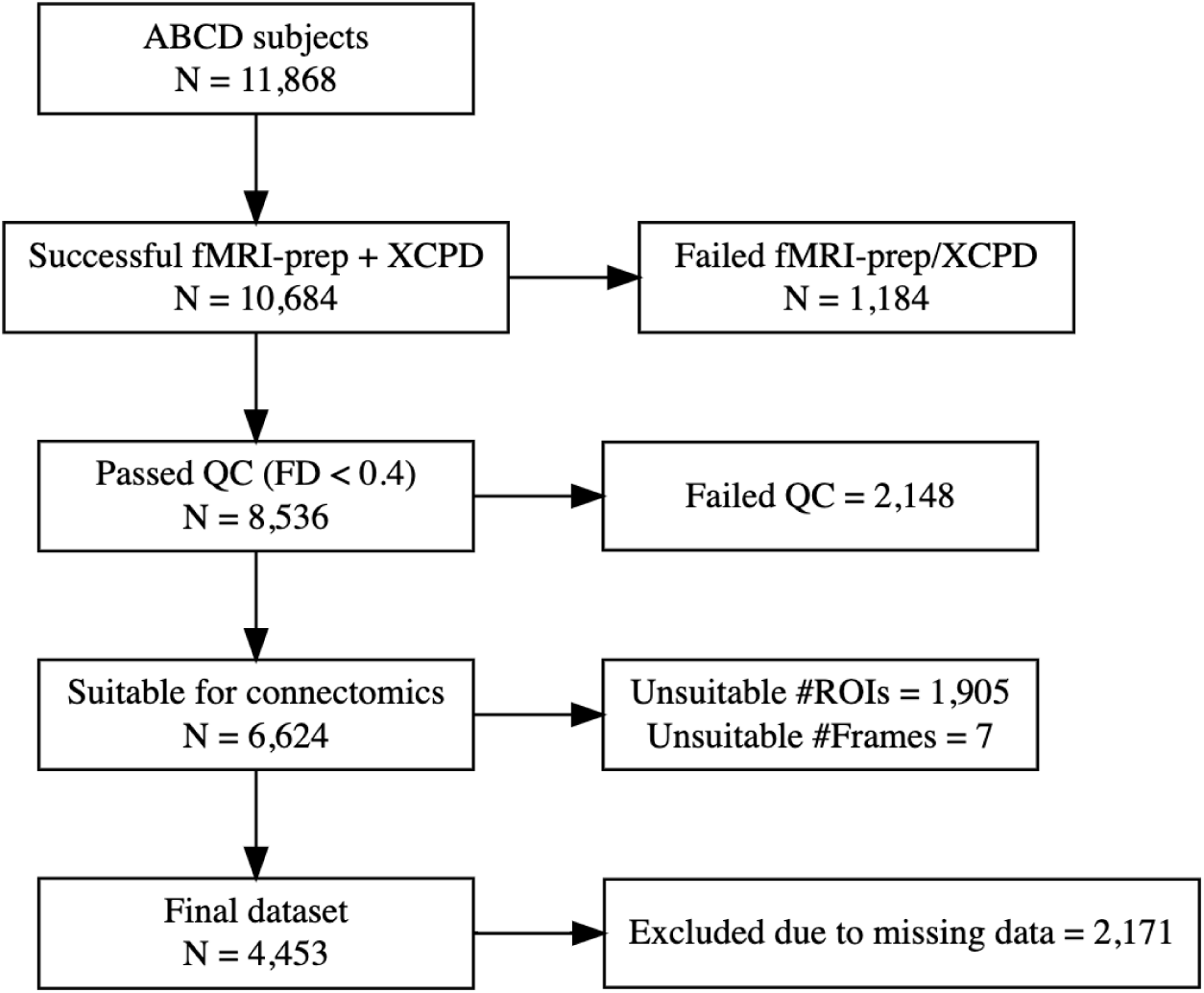
Flowchart of participant inclusion and exclusion criteria. Stepwise selection of the sample from the ABCD baseline cohort (N = 11,868). After fMRI preprocessing with fMRIPrep and postprocessing with XCP-D, data from 10,684 participants were retained for analysis. Of these, 8,536 met imaging quality-control criteria (framewise displacement or FD < 0.4 mm). A total of 6,624 participants were suitable for connectomics analyses after excluding cases with missing values for some ROIs (< 114 ROIs) or less number of usable frames (or timepoints, < 380). The final analytical dataset comprised 4,453 participants after removing individuals with missing data on key demographic or developmental variables. All exclusion counts at each stage are indicated in the figure.

### Low-frequency functional organization in preadolescence revealed by connectome harmonics

Functional connectivity between regions was computed as the correlation of their time series, producing a group-level connectivity matrix parcellated with the 4S atlas, which includes 100 cortical (Schaefer-100 atlas) and 14 subcortical regions (CIT168, Diedrichsen, HCP atlas).

This matrix was then converted into an adjacency matrix for Laplacian eigen decomposition (Figure 2A). To assess robustness across parameter choices, we computed harmonics using nearest-neighbor values (i.e., number of neighboring ROIs considered when computing the adjacency matrix) of 5, 10, 15 and 20; here we reported the nn = 10 solution (with nn number of nearest neighbors used to estimate adjacency matrix), and results for other values are provided in Supplementary Materials (Section 2.1.1). The first eigenvector (eigenvalue = 0), which is spatially constant, was excluded. Figure 2 illustrates the first six harmonics (ψ₁–ψ₆), which collectively capture large-scale spatial gradients spanning both cortical and subcortical areas. We identified the following brain gradients: ψ1 visual-sensorimotor harmonic, ψ2 sensorimotor-multimodal harmonic, ψ3 parieto-temporal harmonic, ψ4 default-mode network (DMN) fronto-parietal harmonic, ψ5 dorsal-attention network (DAN) fronto-parietal harmonic, ψ6 limbic-temporal harmonic (Figure 2B).

**Figure 2.**
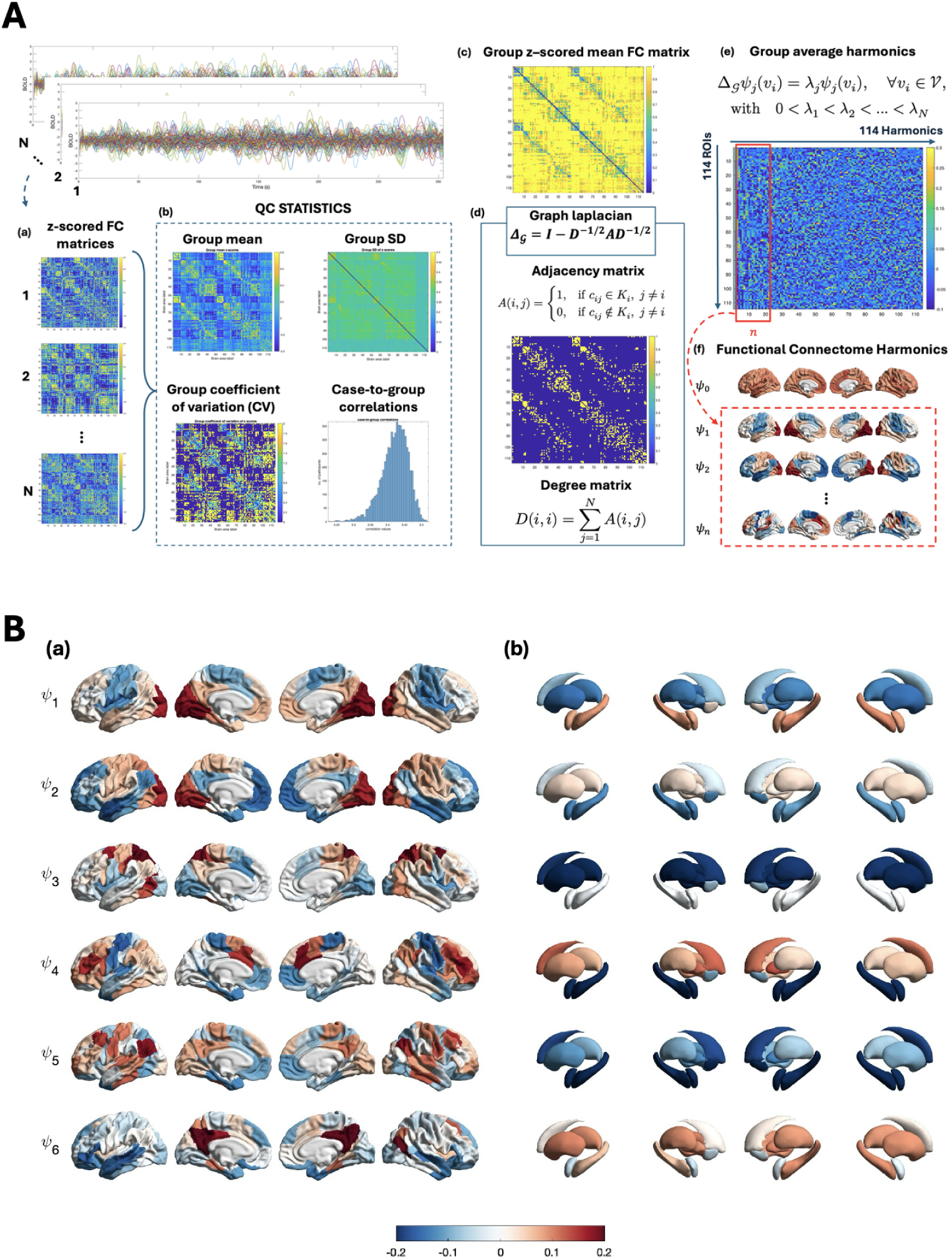
Functional connectome harmonics derived from resting-state fMRI. **A**, Overview of the pipeline used to derive functional connectome harmonics from resting-state fMRI data from 6,624 preadolescents (a-f). **a**, Individual functional connectivity (FC) matrices are computed from BOLD time series and z-scored to standardize across participants. **b**, Group-level mean and standard deviation (SD) of z-scored FC values are calculated, together with the coefficient of variation (CV) and case-to-group correlations, to assess inter-individual consistency. **c**, Z-scored time series from all participants are concatenated and pairwise correlations between brain regions are computed to obtain a group-level FC matrix. **d**, Adjacency and degree matrices are derived from the group-level FC matrix to construct the graph Laplacian, providing a discrete approximation of the Laplacian operator. **e**, Eigen-decomposition of the graph Laplacian yields harmonic modes ordered by increasing spatial frequency, corresponding to principal axes of functional organization; the first *n* harmonics are associated with the lowest eigenvalues. **f**, The resulting eigenmodes are projected onto cortical surfaces, revealing spatially distributed harmonic patterns across the brain. The first harmonic, which exhibits a spatially uniform pattern, is excluded from subsequent analyses. **B**, The first six functional connectome harmonics computed from a group-averaged functional connectivity matrix and displayed on cortical surfaces (**a**) and subcortical structures (**b**). ψ1; visual-sensorimotor harmonic, ψ2; sensorimotor-multimodal harmonic, ψ3; parieto-temporal harmonic, ψ4; default-mode network (DMN) fronto-parietal harmonic, ψ5; dorsal-attention network (DAN) fronto-parietal harmonic, ψ6; limbic-temporal harmonic.

### Preadolescent FCHs converge on canonical adult functional gradients

The first six harmonics were validated using seven reference gradients from NeuroMap (Markello et al. 2022). For each pair of harmonics and reference network, Pearson’s correlations and associated *p*-values were computed and summarized in Figure 3. To ensure comparability with previous studies, reference gradients were selected based on commonly used templates in the literature. These included the first four functional gradients from Margulies et al. (2016), derived using diffusion embedding of the cortical connectivity matrix, the Allometric Scaling gradient from Reardon et al. (2018), which provides a reference for morphometric scaling patterns and the first principal component (PC) of the gene- and cognition-related gradients from the Abagen Toolbox (Markello et al. 2021) and NeuroSynth database (Yarkoni et al. 2011), which summarize the spatial distribution of gene expression and functional activation patterns related to cognitive domains, linking molecular and functional organizational axes. Correlation analyses indicated that the FCHs map onto the major large-scale gradients that characterize adult brain organization. For instance, ψ1 visual-sensorimotor harmonic showed a strong positive correlation with functional gradient #2 (*r* = 0.80) while ψ2 sensorimotor-multimodal sensory harmonic resulted negatively correlated with functional gradient #1 (*r* = −0.80). ψ3 parieto-temporal harmonic was negatively correlated with functional gradient #4 (*r* = −0.65). While the ψ4 DMN fronto-parietal harmonic, showed strong positive correlations with functional gradients #3 (*r* = 0.56), the ψ5 parietal default-mode harmonic, characterised by the parietal components, was positively correlated with the first PC of gene expression (*r* = 0.33). Finally, ψ6 limbic harmonic was positively correlated with Allometric Scaling performed with the NIH database (r = 0.28).

**Figure 3.**
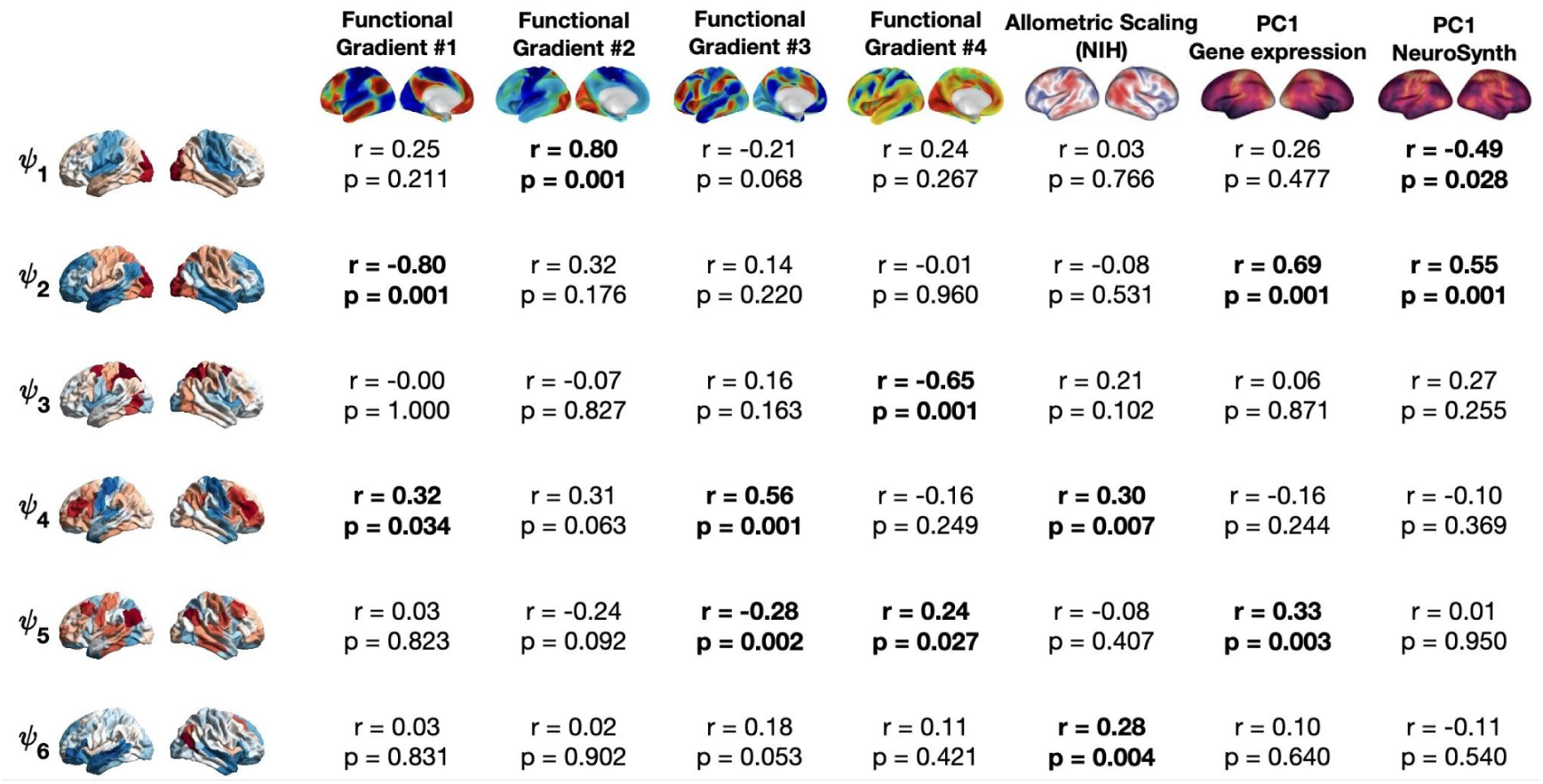
Association between functional connectome harmonics and large-scale brain gradients. Functional connectome harmonics derived from preadolescent resting-state fMRI were compared with reference cortical gradients derived from adult populations. Pearson’s correlation coefficients (*r*) and associated *p*-values (*p*) are reported (N = 6624). NIH; National Institutes of Health, PC1; first principal component. ψ1; visual-sensorimotor harmonic, ψ2; sensorimotor-multimodal harmonic, ψ3; anterior-posterior harmonic, ψ4; DMN fronto-parieta harmonic, ψ5; DAN fronto-parietal harmonic, ψ6 limbic-temporal harmonic.

### LEiDA reveals dynamic network organization in the preadolescent brain

With LEiDA, we identified sets of BOLD phase-locking patterns, each represented as a vector of *N* = 114 elements, capturing the projection of a brain region’s BOLD phase onto the leading eigenvector of all BOLD phases, and providing a compact representation of large-scale synchronization patterns (Figure 4A). To characterize these states, we compared their cortical components with the seven intrinsic functional networks (IFNs) defined by Yeo et al. (2011), restricting the analysis to the 100 cortical Schaefer parcels because the reference networks are cortical. For interpretability, negative centroid values were set to zero, retaining only regions whose BOLD phases aligned with the leading eigenvector. For each clustering solution (K = 2–20), we computed correlations between each centroid and the IFN masks; states exceeding r > 0.4 (p < 10⁻⁴) were assigned to the corresponding network, whereas subthreshold states remained unassigned (Figure 4B). Across clustering solutions, most centroids showed robust correspondence with canonical IFNs, with correlations up to r = 0.846. The DMN and the Visual (VIS) ones appeared consistently and often dominated the low-K solutions, indicating that they represent stable, recurrent modes of resting-state synchronization. Other networks, including DAN, Ventral Attention (VAN), Somatomotor (SMN) and Frontoparietal (FPN), were present even at low K but became progressively more distinct at higher resolutions, suggesting that they reflect finer-grained or less prevalent synchronization patterns. Notably, unlike several prior studies, we did not observe a ‘global brain state,’ which may be attributable to the rigorous preprocessing choices that attenuated large-scale fluctuations (Cabral et al., 2017; Vohryzek et al., 2020). A subset of centroids did not match any IFN and remained unassigned, indicating synchronization patterns not captured by canonical cortical network boundaries. Together, these results provide a systematic functional characterization of LEiDA states and demonstrate that the dominant patterns of preadolescent resting-state dynamics align closely with established large-scale brain networks.

**Figure 4.**
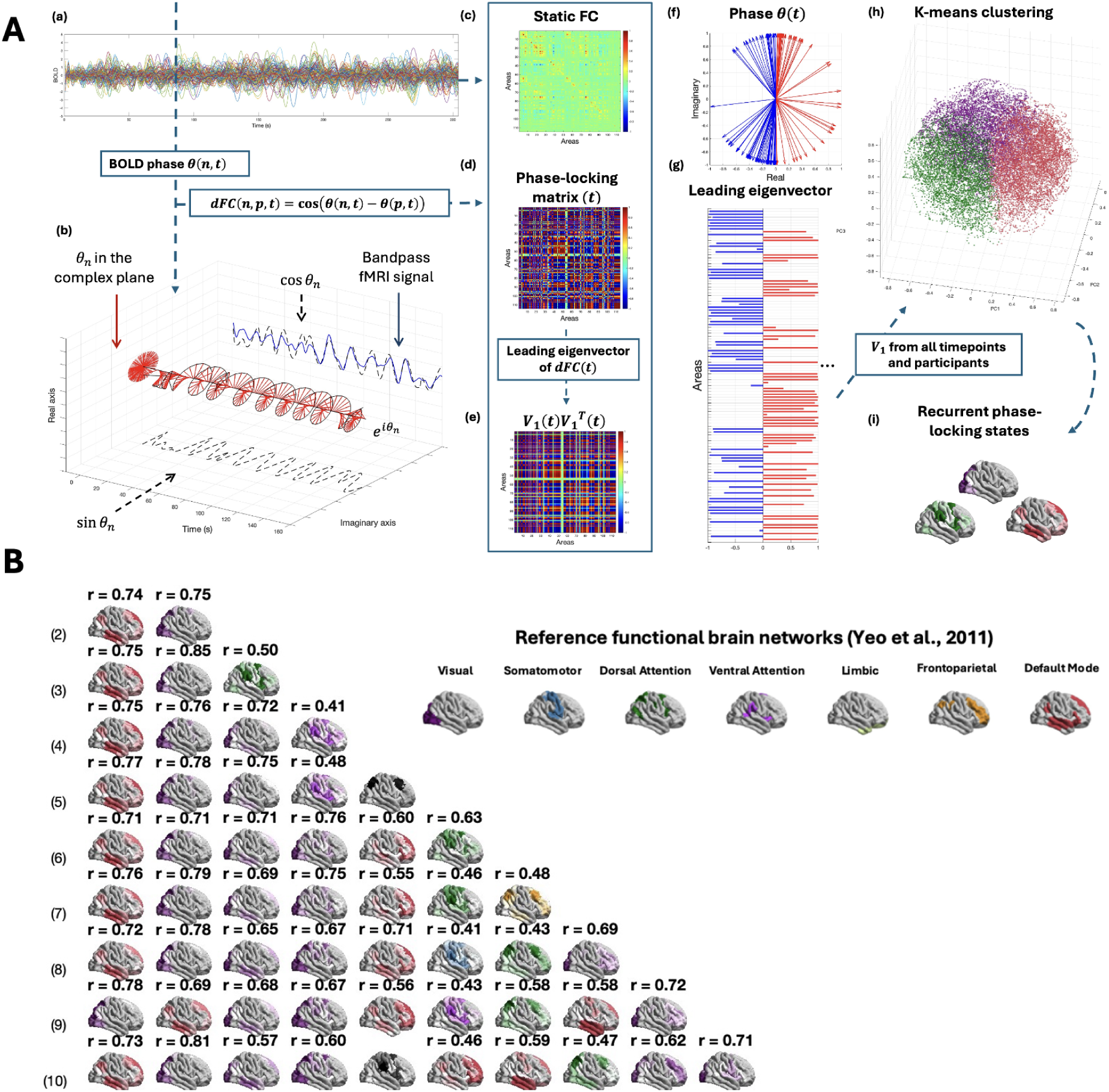
Identification of recurrent whole-brain phase-locking states using LEiDA. **A**, Overview of Leading Eigenvector Dynamics Analysis (LEiDA) applied to resting-state fMRI data from 6,624 preadolescents (a-i). **a**, Resting-state BOLD time series from a representative participant across *N* = 114 brain regions. **b**, BOLD signals are band-pass filtered (0.01–0.1 Hz) and transformed into analytic signals using the Hilbert transform to extract instantaneous phases. **c**, Static functional connectivity (FC) is computed as pairwise correlations between BOLD signals across the entire scan. **d**, Dynamic functional connectivity (dFC) is estimated at each time point using phase coherence, defined as the cosine of phase differences between all pairs of regions, yielding a time-resolved phase-locking (PL) matrix. **e**, The leading eigenvector, *V*₁(*t*), captures the dominant connectivity pattern of dFC(*t*) and allows reconstruction of the principal PL configuration via the outer product. **f**, Instantaneous phases across all regions at a single repetition time are represented in the complex plane; arrow orientation and color indicate the sign of the elements of *V*₁(*t*) (red, positive; blue, negative). **g**, *V*₁(*t*) summarizes the dominant phase-alignment pattern, with regions oscillating in phase or anti-phase relative to the global configuration. **h**, Leading eigenvectors from all time points and participants are concatenated and clustered using *k*-means to identify *k* recurrent PL states; centroids for *k* = 3 are shown. **i**, PL state centroids are projected onto the cortical surface, revealing distinct large-scale phase-locking configurations across the brain. **B**, Correlations between LEiDA-derived PL state centroids and the seven canonical cortical resting-state networks defined by Yeo *et al*. (2011). For each value of *k*, centroids are ordered according to their probability of occurrence, from most to least frequent, and color-coded by the network showing the strongest association. Pearson correlation coefficients (*r*) are reported; centroids with *r* < 0.4 are shown in black. FC, functional connectivity. The figure reports results for K = 2-10; those referred to K=2-20 can be found in the Supplementary Material (Section 2.2.1).

### FCHs delineate the landscape of dynamic phase-locking states

Recent work has demonstrated that brain gradient patterns, such as those derived from FCHs, could form a foundational basis for brain network organization (Rosberg et al., 2026). To test whether FCHs can be used to predict dynamic phase-locking patterns, we used partial least squares regression (PLSR) to model the six LEiDA states (K = 6) using the full set of harmonics, comparing models with ordered versus randomized harmonics. Results showed that one to three PLS components were sufficient to capture the relationship between FCHs and LEiDA states, with most states best fitted by a single component. Across states, models based on ordered harmonics showed the lowest mean-squared errors for states 1 (DMN; MSE = 0.0022), 3 (VIS; MSE = 0.0021), and 5 (DMN; MSE = 0.0016), whereas two- or three-component solutions provided lower errors for states 2 (VIS), 4 (VIS), and 6 (DAN). Model performance for state 1 is illustrated in Fig. 5. In all cases, the variance explained and observed-versus-predicted responses indicated that the original harmonics provided a markedly better fit than the randomized set, demonstrating that the spatial structure of FCHs carries predictive information about dynamic synchronization states. Analyses for states 2--6 are reported in Supplementary Material Section 2.3.

**Figure 5.**
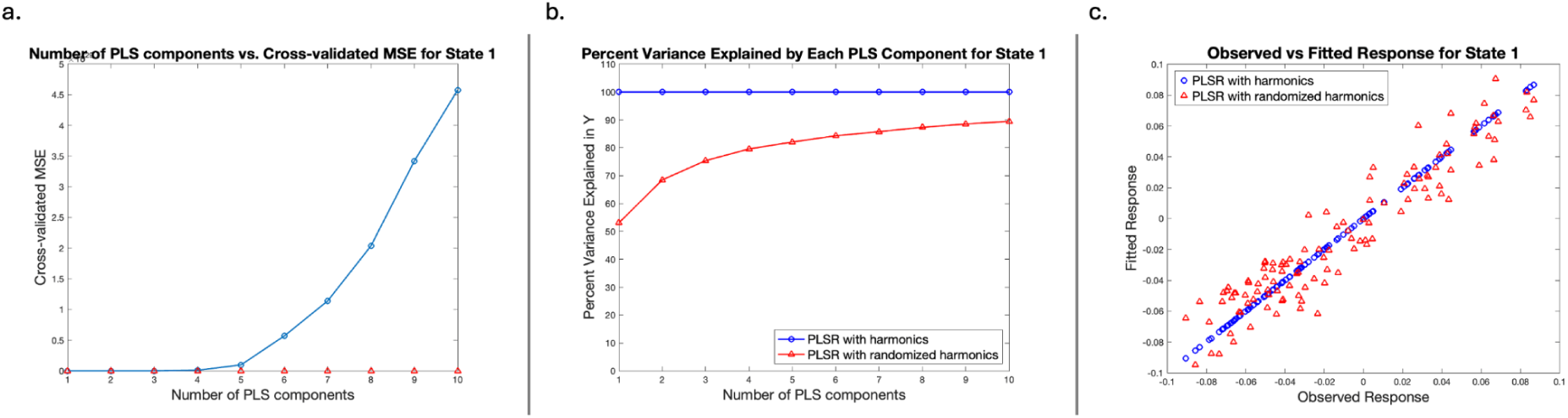
Prediction of Default Mode Network expression from functional connectome harmonics. Partial least squares regression (PLSR) models predicting LEiDA state 1 (default mode network; *K* = 6) from functional connectome harmonic (FCH) components. **a**, Mean squared error (MSE) across PLS components. **b**, Variance explained for ordered and permuted harmonics. **c**, Observed versus predicted values for the selected model (*N* = 6,624).

### Functional harmonic profiles capture subtle demographic and developmental variability

We examined associations between demographic (i.e., sex, ethnicity) and developmental variables (i.e., interview age, PDS, TMI) with the power and energy of the first twenty FCHs (Fig. S12). Power reflects how strongly a given harmonic mode is expressed in the fMRI signal time series, capturing its instantaneous contribution to cortical activity, whereas energy further weights this contribution by the intrinsic spatial frequency of the harmonic, emphasizing modes that contribute more strongly to the dynamic flow of activity across the brain (Supplementary Material, Section 1.3.2 for formal definitions). To mitigate site-related effects inherent to multisite data, ComBat harmonization was applied prior to statistical modeling (Fortin et al. 2018). Results reported below are referred to the harmonized dataset, while corresponding analyses using the original data are summarized in Supplementary Material, Section 2.4, for consistency. The resulting correlations revealed significant positive (blue) and negative (orange) associations exceeding |r| > 0.03 (Fig. 6), which are considered small-to-medium effects (0.03 < r < 0.05) in the context of large population-based neuroimaging studies such as the ABCD study (Owens et al., 2021). ψ₄ energy was positively associated with TMI, ψ₇ energy showed a positive association with interview age (*r* = 0.031), and ψ₁₀ energy was associated with PDS (*r* = 0.031). In addition, ψ₁₄ power and energy were associated with ethnicity (*r* = −0.047 and *r* = 0.042, respectively). ψ₁₆ power was negatively associated with PDS (*r* = −0.030) and TMI (*r* = −0.036), while its energy showed a positive association with TMI (*r* = 0.040). Finally, ψ₁₈ power and energy were again associated with ethnicity (*r* = −0.043 and *r* = 0.037, respectively). Together, these findings indicate that demographic and developmental variables show selective and reproducible associations with specific functional connectome harmonics.

**Figure 6.**
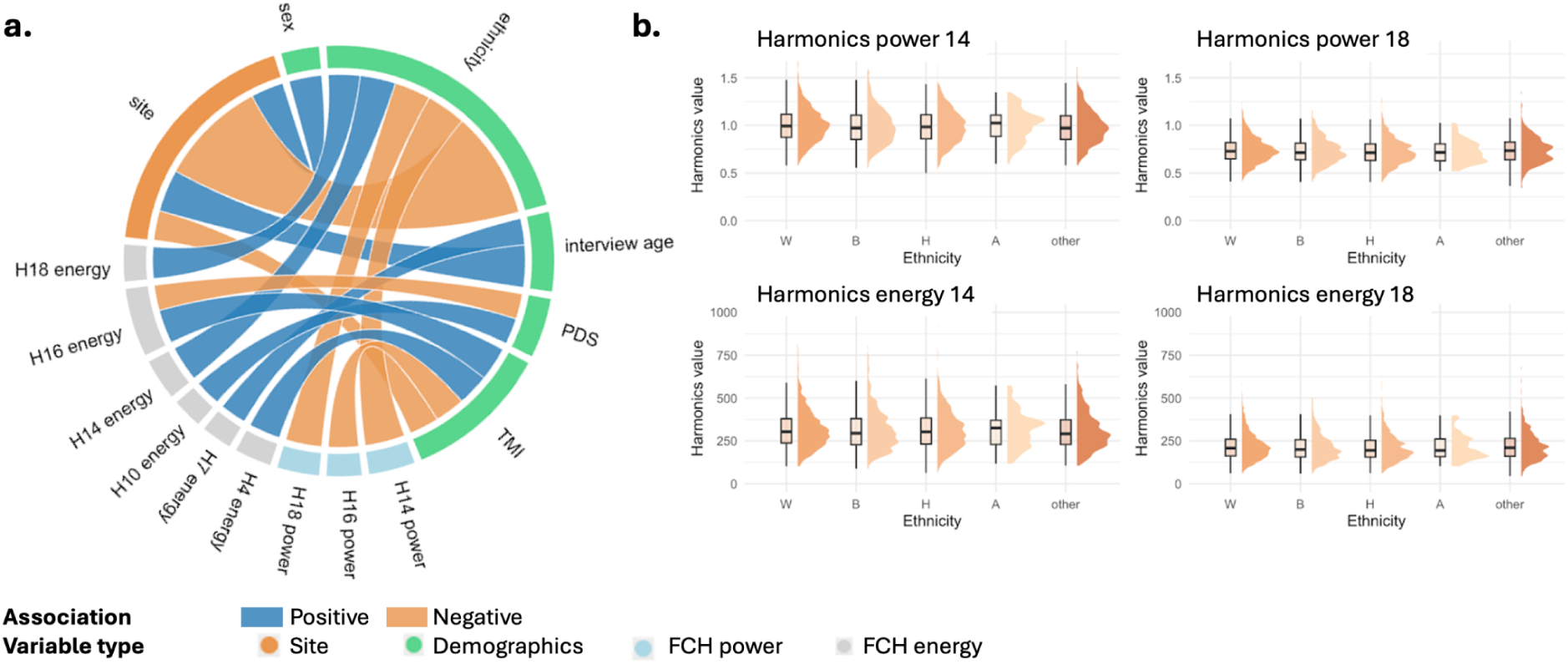
Correlations between FCH metrics and demographic and developmental variables. **a,** Circos plots showing significant partial correlations (|r| > 0.03) between FCH energy (grey) and power (light blue) with key variables. **b,** Half-violin plot showing the distribution of FCH-derived metrics significantly associated with ethnicity. The x-axis represents ethnicity groups: W = White, B = Black, H = Hispanic, A = Asian, other (i.e., any other different ethnicity from those analysed); the y-axis indicates the metric value (N = 4453). PDS; pubertal development scale, TMI; triponderal mass index.

### Phase-locking states capture subtle demographic and developmental variability

We performed the same set of analyses on all LEiDA-derived metrics—fractional occupancy (FO), dwell time (DT), and Markov chain transition probabilities—for every state that could be assigned to one of the Yeo *et al*. (2011) networks across *K* = 2–20. FO reflects the relative proportion of time a participant spends in a given phase-locking (PL) state, providing a measure of the dominance of each recurrent brain configuration, whereas DT quantifies the average duration of uninterrupted visits to a state, capturing the temporal stability of that configuration. Markov chain transition probabilities index the likelihood of transitioning between pairs of states across time, thereby characterizing the structure of state-to-state dynamics. For each metric, partial correlations were computed while regressing out all other demographic covariates. P values were FDR-corrected, and only associations surviving correction (p < 0.05) were considered significant. Effect sizes ranged from |r| ≈ 0.04 to 0.14, with a subset approaching r ≈ 0.18. Overall, LEiDA-derived network states showed significant and largely consistent associations with demographic covariates. For fractional occupancy, the Visual network displayed the strongest and most consistent associations, showing negative relationships with age and positive associations with PDS and sex. The Default Mode network (DMN) was primarily associated with ethnicity and sex, whereas other networks, including the frontoparietal and Dorsal attention networks, showed associations mainly with sex (Fig. 7a). For dwell time, age and PDS were negatively associated with Visual network states, while TMI exhibited significant negative associations across Visual, Frontoparietal, and Dorsal Attention states. Ethnicity was selectively associated with DT within the DMN, and sex showed widespread associations across DMN, Visual, and Dorsal attention networks (Fig. 7b). Markov chain transition probabilities revealed the strongest overall effects. Age negatively influenced transitions involving the DMN and Visual networks, with additional significant transitions emerging in Frontoparietal and Ventral Attention networks. Ethnicity was primarily associated with DMN and Visual transitions, with further associations spanning Dorsal Attention, Ventral Attention, Frontoparietal, and Somatomotor networks. PDS was linked to transitions involving DMN, Visual, Dorsal Attention, and Frontoparietal networks, whereas sex showed broad effects across DMN, Visual, Dorsal Attention, Ventral Attention, Frontoparietal, and Somatomotor transitions. TMI was associated with both negative and positive transitions across DMN, Visual, Dorsal Attention, Frontoparietal, and Somatomotor networks (Fig. 7c).

**Figure 7.**
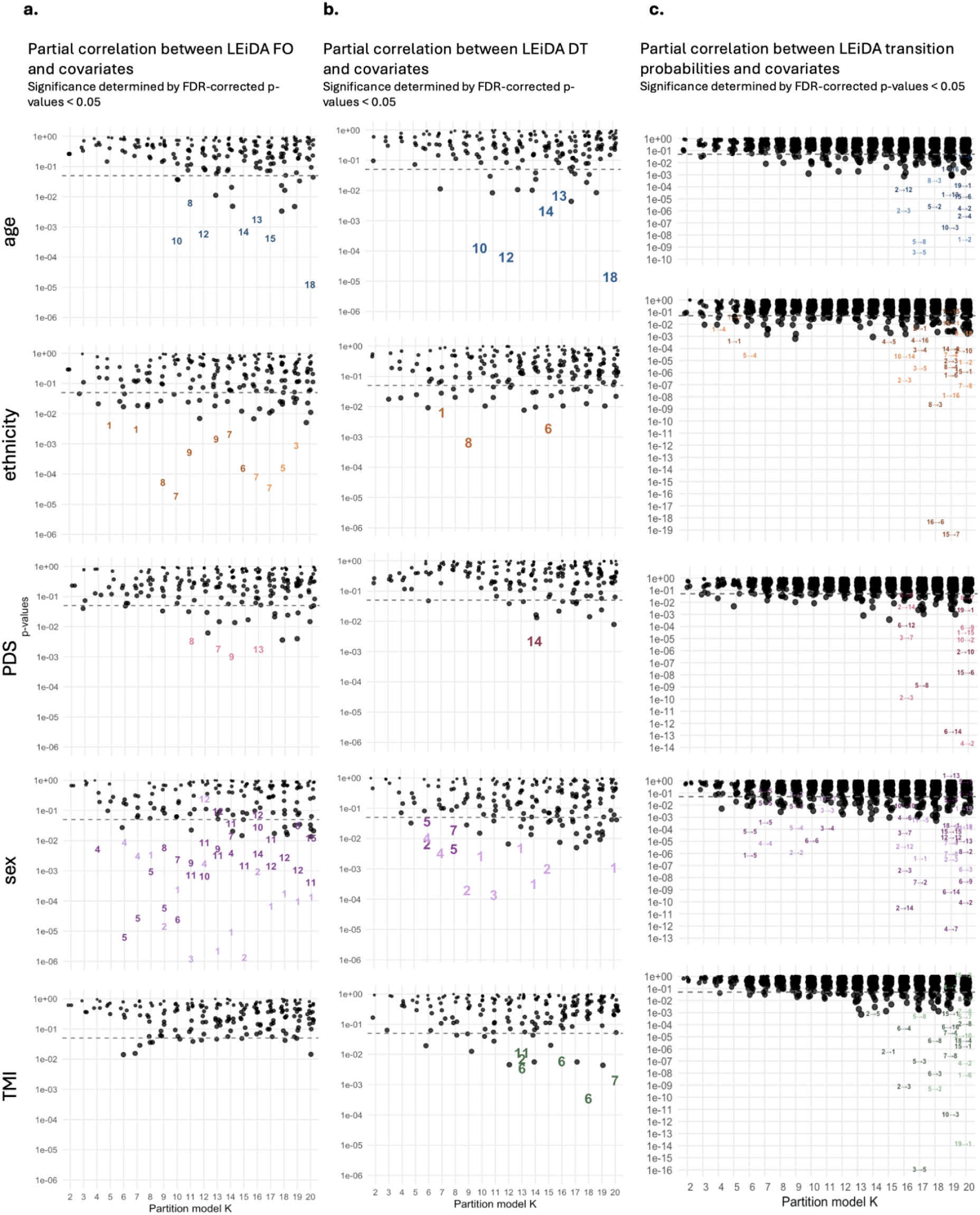
Partial correlations between LEiDA lifetimes, probabilities, transitions and demographics and developmental variables. Panels a–c show results for LEiDA probabilities, lifetimes, and transitions, respectively; each panel contains subplots for sex, ethnicity, interview age, Pubertal Development Scale (PDS), and Triponderal Mass Index (TMI). In each subplot, the x-axis shows cluster number (k = 2–20) across 19 partition models, and the y-axis shows p-values for each PL-state. PL-states with FDR-corrected p > 0.05 are black dots. Numbers indicate the PL-state involved in the association. Positive associations are represented with lighted colors, while negative associations are represented with darker colors.

**Figure 8.**
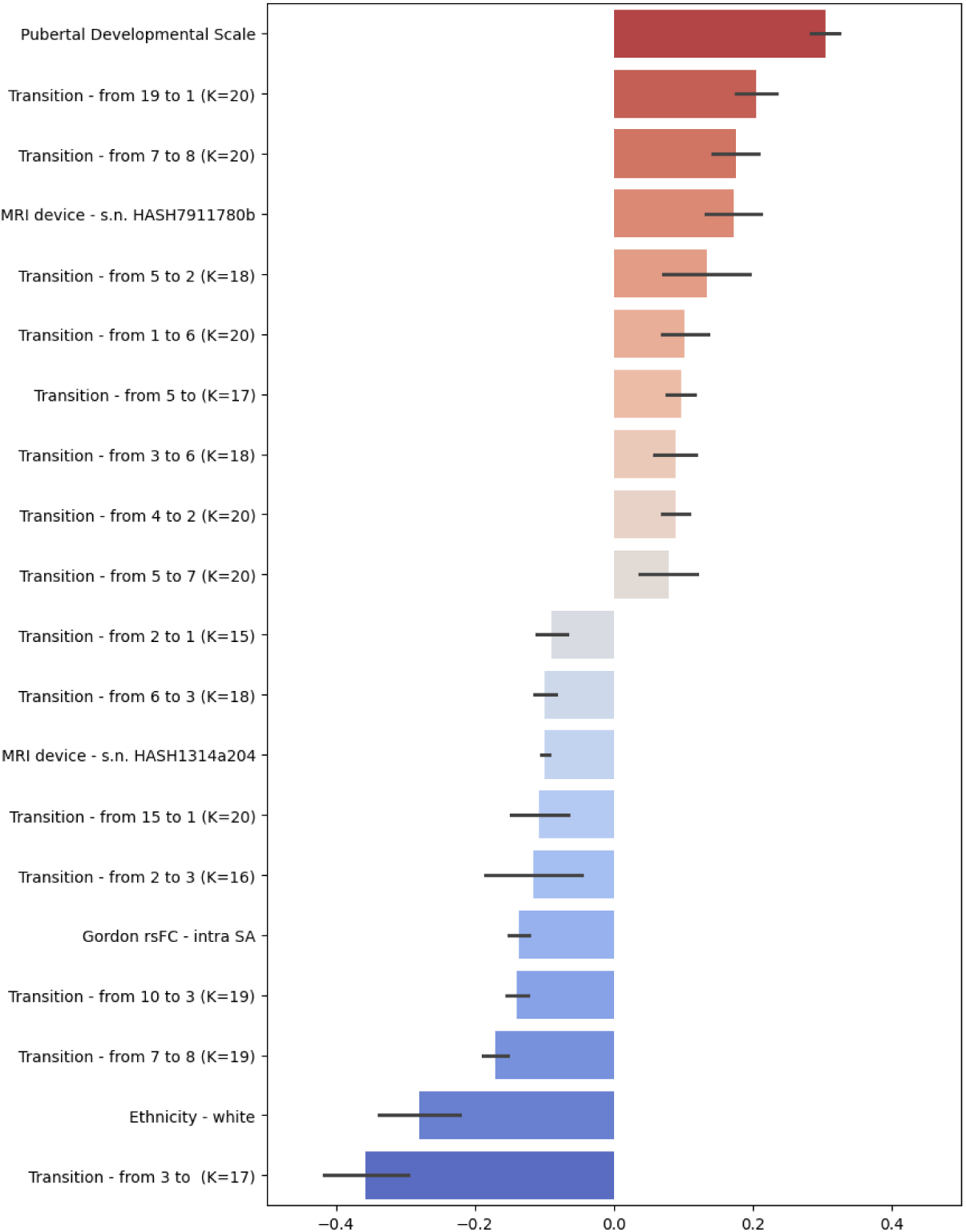
Multivariate predictors of triponderal mass index from combined functional brain features. Ranking plot of feature importance in the best-performing Elastic Net model predicting triponderal mass index (TMI). The model combines demographic covariates with resting-state fMRI connectivity, functional connectome harmonic (FCH) measures and LEiDA-derived dynamic metrics. Bars show the top positive (red) and negative (blue) standardized coefficients, reflecting each feature’s (y-axis) relative contribution to TMI prediction.

### Functional connectivity metrics enhance TMI prediction beyond conventional rs-fMRI features

Previous work has shown that resting-state fMRI features are among the weakest predictors of TMI as compared with structural MRI and DTI measures (Adise et al., 2021; Suuronen et al., 2026 - article in preparation). We therefore tested whether connectome-derived functional metrics—specifically FCHs and LEiDA states—could enhance the predictive value of rs-fMRI. To this end, we used ElasticNet regression to predict TMI from covariates augmented with rs-fMRI features, LEiDA-derived metrics, FCH measures, and their combination. Analyses were performed on both the original and harmonized datasets, with full results for the original dataset reported in the Supplementary Material (Section 2.6). In the harmonized dataset (Table 2), rs-fMRI and FCH features showed comparable performance to prior work using rs-fMRI data (R² = 0.13 training; R² = 0.08 test), whereas LEiDA metrics provided substantially improved predictive accuracy (R² = 0.26 training; R² = 0.15 test). Combining all imaging features yielded the strongest overall performance (R² = 0.29 training; R² = 0.17 test).

**Table 2:** Model performance for predicting triponderal mass index from demographic, developmental and brain-based features. Performance of elastic net regression models trained to predict triponderal mass index (TMI) using different feature sets in the harmonized dataset (N = 4,453). Covariates; demographic covariates only, Rs-fMRI; whole-brain resting-state fMRI connectivity, LEiDA; LEiDA dynamic-state metrics, FCH; FCH-derived metrics, All; all features combined. Metrics reported include the coefficient of determination (R2) and negative mean absolute error (negMAE), averaged across 5 cross-validation folds, with standard deviation (STD) shown for both training and held-out test sets.

| Model | Metric | Training – mean | Training – STD | Test – mean | Test – STD |
| --- | --- | --- | --- | --- | --- |
| Covariates | R2 | 0.1045 | 0.0025 | 0.0865 | 0.0084 |
|  | negMAE | -1.7577 | 0.0119 | -1.7726 | 0.0308 |
| Rs-fMRI | R2 | 0.1693 | 0.0134 | 0.1099 | 0.0135 |
|  | negMAE | -1.6953 | 0.0210 | -1.7491 | 0.0218 |
| LEiDA | R2 | 0.2578 | 0.0155 | 0.1489 | 0.0330 |
|  | negMAE | -1.6179 | 0.0077 | -1.7114 | 0.0166 |
| FCH | R2 | 0.1111 | 0.0026 | 0.0849 | 0.0107 |
|  | negMAE | -1.7517 | 0.0116 | -1.7752 | 0.0287 |
| All | R2 | 0.2890 | 0.0158 | 0.1679 | 0.0343 |
|  | negMAE | -1.5841 | 0.0084 | -1.6945 | 0.0116 |

## Discussion

In this study, we provide the first large-scale characterization of functional connectome harmonics and dynamic coupling states in preadolescence using resting-state fMRI data from the ABCD Study. By integrating Functional Connectome Harmonics (FCH), which capture intrinsic spatial modes of synchronous activity (Atasoy et al., 2016; Atasoy et al., 2018; Glomb et al., 2021), with Leading Eigenvector Dynamics Analysis (LEiDA), which identifies recurrent phase-locking states (Cabral et al., 2017; Vohryzek et al., 2020), we show that large-scale functional organization in 9–10-year-olds is governed by a low-dimensional architecture linking spatial gradients with temporal dynamics.

FCH revealed dominant harmonic modes that closely resembled canonical cortical gradients identified in adults (Margulies et al., 2016; Huntenburg et al., 2018; Markello et al., 2022), demonstrating that the principal axes of large-scale cortical organization are already robustly expressed by late childhood. Rather than undergoing wholescale reconfiguration, these gradients appear to be undergoing refinement, echoing developmental models proposing early stabilization of sensory–transmodal hierarchies (Sydnor et al., 2021; Dong et al., 2021). Low-frequency harmonics spanning visual, sensorimotor, limbic, and default-mode territories support foundational aspects of sensory processing, internal mentation, and emerging transmodal integration (Burt et al., 2018; Reardon et al., 2018; Valk et al., 2020; Pang et al., 2023).

Dynamic analyses revealed a similarly structured repertoire of brain states. LEiDA-derived phase-locking states were dominated by Default Mode and Visual configurations, which were highly recurrent and stable across clustering solutions (Allen et al., 2014; Vidaurre et al., 2017). In contrast, attention and control networks, including Dorsal and Ventral Attention and Frontoparietal systems, appeared more transient and fragmented, consistent with their protracted maturation (Fair et al., 2009; Luna et al., 2010; Sydnor et al., 2021). The convergence of these developmental signatures across spatial and temporal analyses suggests that distinct maturational timetables are already inscribed into the brain’s large-scale dynamics by late childhood.

A central conceptual advance of this work is the demonstration that dynamic phase-locking states are predicted by the FCH’s. Transient synchronization follows the geometry defined by low-frequency eigenmodes, providing empirical support for theoretical accounts positing that large-scale neural activity evolves along functional connectivity eigenbasis (Atasoy et al., 2016; Abdelnour et al., 2018; Bolt et al., 2022; Pang et al., 2023; Runfola et al., 2025). That only one to three PLSR components were required to map LEiDA states onto harmonics indicates that the dynamic repertoire of the preadolescent brain can be described using a compact set of functional components.

Associations with demographic and developmental variables were systematic and serve as a practical validation for our relatively novel derived brain metrics. Higher-order harmonics and dynamic features showed the greatest sensitivity to age, sex, and pubertal status, consistent with evidence that fine-scale functional differentiation continues to mature during adolescence, whereas broad cortical gradients remain comparatively stable (Laumann et al., 2015; Kong et al., 2019; Sydnor et al., 2021; Dong et al., 2021). LEiDA-derived DMN and Visual states showed the most consistent associations to demographic variables. Importantly, effect sizes must be interpreted within the empirical landscape of large-scale cohorts: correlations in ABCD are systematically small (Owens et al., 2021), but even small associations may accumulate meaningfully across contexts (Funder & Ozer, 2019). Within this framework, the magnitude of our effects is consistent with meaningful developmental variability.

Predictive modeling further underscored the relevance of functional dynamics. LEiDA-derived features substantially improved prediction of TMI compared with static connectivity or harmonic-derived measures (Adise et al., 2021), and combining all imaging features yielded the strongest performance. This synergy suggests that static, harmonic, and dynamic measures capture complementary aspects of brain function and that dynamic signatures might carry unique relevance in preadolescence.

Several limitations merit consideration. Cross-sectional data limit inference about developmental trajectories; longitudinal analyses will be essential for tracking how harmonic structure and dynamic repertoires evolve through adolescence. Despite harmonization efforts, residual multisite effects may persist as indicated by site variables in our elastic net regression model. Low-dimensional representations may omit finer temporal or amplitude-related features, and group-level harmonics may obscure meaningful individual variation. Predictive performance of TMI was comparable to that reported for diffusion-based metrics in prior work (Adise et al., 2016; Suuronen et al., 2026 - article in preparation), indicating that LEiDA-derived fMRI measures achieve similar predictive accuracy to DTI. This finding extends developmental prediction beyond structural microarchitecture and demonstrates that functional brain dynamics can provide comparable predictive information. Integrating structural, diffusion, functional, genetic, and environmental factors will be critical for further improving the precision of neurodevelopmental outcome models.

Beyond its conceptual contributions, this study also represents a methodological milestone: it provides the first harmonized derivation of functional connectome harmonics and LEiDA-derived dynamic states in the ABCD cohort, creating a foundational resource for future developmental and longitudinal investigations. These harmonized measures offer a scalable framework for characterizing functional architecture in large pediatric samples and for investigating how individual differences in spatial and temporal brain organization may relate to emerging cognitive, emotional, and mental health trajectories.

Taken together, our findings indicate that the developing brain can be described by a low-dimensional harmonic scaffold that captures aspects of its dynamic repertoire during a critical window of neurodevelopment. By linking spatial gradients to temporal transitions, this work provides a framework for investigating how large-scale neural dynamics may relate to the emergence of cognition and behavior, laying essential groundwork for future studies investigating developmental trajectories and their relevance for adolescent mental health.

## Methods

### Participants

Neuroimaging and demographic data analyzed in this study were obtained from the baseline assessment of the Adolescent Brain Cognitive Development (ABCD) Study, Data Release 5.0 (https://abcdstudy.org/scientists/data-sharing/). The ABCD Study is a large longitudinal cohort that recruited approximately 11,880 children aged 9–11 years across 21 research sites in the United States. Parental written informed consent and child assent were obtained in accordance with local institutional review board approvals.

After standard exclusion criteria were applied—including medication use, relevant health conditions, MRI data quality, and the removal of one participant per twin pair—the final sample comprised 4,453 children aged 9–10 years. Demographic and developmental covariates included sex, ethnicity, age at interview, pubertal development scale scores (PDS), and triponderal mass index (TMI). Additional details on recruitment and data acquisition are available in the ABCD Study protocols (https://abcdstudy.org/scientists/protocols/) and associated publications (Casey et al., 2018; Garavan et al., 2018).

### Brain data preprocessing

Structural and functional MRI data were preprocessed using fMRIPrep v23.1.3 (Esteban et al. 2018, 2019) and XCP-D v0.7.0 (Ciric et al. 2018; Satterthwaite et al. 2013). The pipeline was applied to a total of 11,868 subjects, representing a cumulative computational investment of approximately 100,000 CPU hours (averaging 8 hours for fMRIPrep and 30 minutes for XCP-D per subject). Anatomical preprocessing included intensity non-uniformity correction, skull-stripping, tissue segmentation (CSF, WM, GM), surface reconstruction (FreeSurfer 7.3.2), and spatial normalization to MNI152NLin2009cAsym and MNI152NLin6Asym using ANTs-based nonlinear registration. Functional data underwent slice-timing correction, motion correction (FSL’s mcflirt), fieldmapless distortion correction, and coregistration to the T1w image using bbregister. Confound regressors (e.g., motion parameters, global signals, CompCor components) were calculated, and outputs were resampled to both volumetric and surface spaces. Postprocessing with XCP-D followed the 36-parameter (36P) denoising strategy, including regression of motion parameters, tissue signals, and their derivatives/quadratics. Data were bandpass filtered (0.01–0.08 Hz) and smoothed with a 6 mm FWHM Gaussian kernel. Quality controlled, denoised BOLD time series were extracted using various parcellation atlases (e.g., Schaefer, Glasser, Gordon, Tian), and pairwise functional connectivity was computed using Pearson correlations. To reduce the influence voxel-wise motion artifacts, only participants with a framewise displacement (FD) below 0.4 were retained for further analysis.

Initial acquisitions were sampled using the 4S156 reference atlas, which included 100 cortical Regions of Interest (ROIs) from Schaefer100 atlas (Schaefer et al. 2018) and 56 subcortical ROIs from the COI168 (Pauli et al., 2018), ThalamusHCP (Najdenovska et al. 2018) and SubcorticalHCP (Glasser et al. 2013) atlases. Then, ROIs were relabeled using a different atlas composed of the same 100 cortical ROIs from Schaefer100, along with 14 subcortical means obtained by averaging preselected subcortical regions to match ENIGMA Toolbox requirements (Larivière et al. 2021), used to visualize connectome analysis results on cortical and subcortical surfaces. The 14 subcortical ROIs obtained for the analyses consisted of bilateral accumbens, amygdala, caudate, hippocampus, pallidum, putamen and thalamus. The list with all ROIs included in the analysis can be found in Supplementary Material Section 1.2. Only subjects with a complete set of 114 ROIs and 380 frames (TR = 0.8*s*) were retained for connectome analyses, resulting in a final sample of 6,624 subjects. These steps were performed on CSC’s high-performance computing infrastructure (Puhti), using a Compute Node Shell with 4 CPUs and 16 GB of memory, and required a few hours to complete.

### Functional Connectome Harmonics

Functional Connectome Harmonics (FCHs) were computed from preprocessed rs-fMRI mean time-series, parcelled using the Schaefer100 atlas (Schaefer et al. 2018) along with 14 additional subcortical ROIs derived from the COI168 (Pauli et al., 2018), Thalamus HCP (Najdenovska et al. 2018), and Subcortical HCP (Glasser et al. 2013) atlases. Pairwise correlations over time between brain regions quantified the functional connectivity of the time series across cortical and subcortical areas. The resulting group-level connectivity matrix was used to derive into group-level adjacency and degree matrices, which were then used to model the FCHs via eigendecomposition of the Laplace operator. Therefore, the number of harmonics equals the number of ROIs, ordered accordingly to the respective eigenvalues in increasing order, from the lowest to the highest frequency. By definition, the first functional harmonic (associated to the null eigenvalue) is constant across the cortex and was therefore discarded from all analyses and visualizations. To ensure robustness, FCHs were computed using different numbers of nearest neighbors (*nn* = 5, 10, 15, 20), which defines the number of ROIs considered when constructing the adjacency and degree matrices. See Supplementary Material, Section 2.1.1 for further details. The FCH pipeline was applied to resting-state fMRI time series organized, for each subject, as a 114 × 380 matrix, where 114 corresponds to the number of ROIs and 380 to the number of acquired frames (TR = 0.8 s). The analysis was run on the full sample of subjects (N = 6,624), using Matlab 2022a on a desktop computer.

### Leading Eigenvector Dynamics Analysis

LEiDA was chosen as the processing method for investigating network dynamics because it can detect activation patterns in the brain with high temporal precision, unlike correlation-based measures of functional connectivity (Cabral et al., 2017). Before running LEiDA, we first obtained the average fMRI signals from 100 cortical and 14 subcortical areas defined by the Schaefer atlas (Schaefer et al., 2018). The time courses were bandpass filtered (0.02 – 0.10 Hz), the analytic phase was obtained using the Hilbert transform, and the leading eigenvectors of the phase coherence matrices were calculated at each time point (Cabral et al., 2017). The eigenvectors were then clustered via K-means clustering using cosine distance with 200 replicates. For this exploratory study, the number of clusters (K) was varied between 2 and 20, based on previous studies (Vohryzek et al., 2020). Clustering yields K states of brain activity, each state representing a recurrent pattern of functional connectivity, and whose probability of occurrence can be used in statistical inference. To facilitate interpretation of BOLD PL states and following the methodology described in Vohryzek et al. (2020), we used as a reference the seven resting-state networks (RSNs) from Yeo et al. (2011), namely, Visual Network (VIS), Somatomotor Network (SMN), Dorsal Attention Network (DAN), Salience Ventral Attention Network (VAN), Limbic Network, Frontoparietal Network (FPN) and Default Mode Network (DMN), obtained from correlation-based functional connectivity across 1175 ROIs in 1000 participants. For comparability, analyses were restricted to the first 100 cortical ROIs of the Schaefer100 parcellation (Schaefer et al. 2018), as IFNs are defined only in cortical space (Yeo et al. 2011). Binary masks were defined for the seven non-overlapping networks in Schaefer100 (Schaefer et al. 2018), yielding seven 1 × 100 vectors (see Supplementary Material (Section 1.2) for ROI-IFN correspondence). Pearson’s correlations (with associated *p*-values) were computed between these network vectors and the centroids across all *K*. Negative centroid values were set to zero, restricting the analysis to regions with a positive phase shift from the main orientation. Each centroid was assigned to the IFN with the highest correlation. A threshold of *r* > 0.4 (*p* < 10−4) was applied, below which centroids were considered unassigned to any IFNs and excluded from further analyses. Results were visualized using the ENIGMA Toolbox (Larivière et al. 2021). We thresholded negative phases to zero, and compared centroids with the seven Yeo et al. (2011) networks. Centroids with maximum correlation below 0.4 were considered unassigned. We ran all the modelling in Matlab 2022a. The LEiDA pipeline was applied to resting-state fMRI time series organized, for each subject, as a 114 × 380 matrix, where 114 corresponds to the number of ROIs and 380 to the number of acquired frames (TR = 0.8 s). Computations were performed on CSC’s high-performance computing infrastructure (Puhti), using 40 CPUs with 8 GB of memory each. The analysis was run on the full sample of subjects (N = 6,624) and required approximately 24 hours to complete. All the following calculations, like comparison to Yeo RSNs, have been run on a desktop computer using Matlab 2022a.

### Linking FCH spatial patterns and LEiDA temporal dynamics

To evaluate whether FCHs can predict patterns of dynamic phase-locking states, we used LEiDA centroids for *K* = 6 as the outcome. The FCHs matrix, along with a randomized set of harmonics, was used to predict the LEiDA states using Partial Least Squares Regression (PLSR). PLSR was chosen because it is particularly suitable for high-dimensional and collinear data, as it extracts latent components that maximize the covariance between predictors and outcomes. This makes it well adapted for identifying the most relevant harmonic patterns associated with dynamic brain states. The optimal number of PLSR components was determined via 10-fold cross-validation, selecting the model that minimized the Mean Squared Error (MSE) of prediction. The analysis was conducted separately for the original FCH matrix and its randomized counterpart, on a desktop computer using Matlab 2022a.

### Derived measures

We extracted FCH power and energy for each harmonic, and for LEiDA, we computed the lifetime, probability, and transition probability of each dynamic state across all K. According to Atasoy et al. (2017), to characterize FCHs we estimated power and energy as additional measures (see Supplementary Material (Section 1.3.2)). These metrics provide complementary information on the contribution of each harmonic mode to the overall cortical dynamics. Specifically, the power of a harmonic brain state reflects the instantaneous strength of activation of a given harmonic at a particular time point, quantifying how strongly the spatial pattern of brain activity projects onto that harmonic. The energy, in turn, represents the frequency-weighted contribution of each connectome harmonic to the ongoing cortical dynamics, by combining its instantaneous power with its intrinsic frequency, captured by the associated eigenvalue. Power was obtained by taking the absolute value of the normalized FCH projection at each time point and averaging it across time, thus reflecting the mean amplitude of activation of each harmonic mode. Energy, instead, was computed as the squared product of the harmonic amplitude and its intrinsic spatial frequency (represented by the corresponding eigenvalue), averaged across time and summed over the cortex, capturing the frequency-weighted contribution of each harmonic to cortical dynamics. Both power and energy values were estimated separately for all harmonics (*n* = 113, excluding the first one associated to the null eigenvalue), derived considering *nn* = 10 nearest neighbors, and all participants (*N* = 6, 624). According to Vohryzek et al. (2020), to characterize LEiDA-derived PL states we calculated additional metrics, including Fractional Occupancy (FO), Dwell Time (DT), and Markov chain transition probabilities (see Supplementary Material (Section 1.4.2)). FO reflects the relative amount of time a subject spends in each PL state, providing a measure of how dominant each recurrent brain pattern is. DT quantifies the average duration of uninterrupted periods in a given state, capturing the temporal stability of brain configurations. Markov chain transition probabilities represent the likelihood of moving from one PL state to another, describing the temporal sequence of state transitions. These metrics were calculated using the cluster assignments obtained from k-means applied to the leading eigenvectors of the phase-locking matrices. FO was estimated as the proportion of time points assigned to each state for every subject. DT was computed by identifying consecutive sequences of time points in the same state and averaging their lengths. Transition probabilities were calculated by counting the number of times each state was followed by another for each subject, normalizing by the total number of transitions, and adjusting for the fractional occupancy of each state to account for differences in prevalence. Any undefined values arising from empty states were set to zero. All calculations were performed separately for each subject in the dataset (*N* = 6, 624) and for all tested numbers of clusters (*K* = 2–20).

### Data harmonization

Given the presence of 21 acquisition sites differing by scanner and location, we applied the ComBat harmonization procedure (Johnson, Li, and Rabinovic 2007; Fortin et al. 2018) to control for site effects, before running statistical analyses. Harmonization was performed using ComBat algorithm as implemented in neuroCombat package in Python (https://github.com/Jfortin1/neuroCombat). ‘Site’ was modeled as the batch variable, while preserving variability associated with the covariates. The covariates included in the harmonization were sex, ethnicity, interview age, PDS, and TMI. Harmonization was performed separately for the three groups of fMRI-derived features: (i) rsfMRI Gordon network and subcortical correlations, (ii) FCH-derived power and energy, and (iii) LEiDA-derived FO, DT, and Markov Chain transition probabilities. LEiDA metrics were included only for states previously associated with one of the seven IFNs (Yeo et al. 2011), while FCH metrics were considered for all harmonics. This procedure removed site-related variability while retaining meaningful inter-individual differences, thereby enabling more reliable and generalizable analyses.

### Statistical analysis

The following statistical analyses were performed on the previously constructed datasets, including both the original (see Supplementary Material, Section 2.5) and the harmonized versions. Performing the same analyses on both datasets allowed us to assess potential differences introduced by the harmonization procedure.

### Partial correlation analysis

Following the approach of Alonso (2023), statistical analyses focused on partial correlation analyses between LEiDA- and FCH-derived metrics and demographic and developmental variables (sex, ethnicity, interview age, PDS and TMI). Partial correlations were used to quantify the association between pairs of variables while controlling for the remaining covariates, thereby isolating their unique contributions and reducing potential confounding effects. FCH metrics were examined for the first 20 harmonics, whereas LEiDA metrics were included only for states associated with one of the seven intrinsic functional networks (IFNs; Yeo et al., 2011). For each harmonic, partial correlations were computed between FCH-derived power and energy metrics and each covariate, regressing out the effects of all remaining variables. Partial correlation coefficients with absolute values greater than 0.03 were considered potentially relevant. For LEiDA, associations between demographic variables and FO, DT, and Markov chain transition probabilities were assessed. Analyses were conducted separately for each of the 19 partition models derived from k-means clustering. Resulting p-values were adjusted for multiple comparisons using the false discovery rate (FDR) procedure (Benjamini & Hochberg, 1995; Figueroa et al., 2019), with associations considered significant at p < 0.05 after correction. Partial correlation analyses were performed using the *ppcor* package in R (Kim, 2015) and were applied to both the original and harmonized datasets.

### ElasticNet regression

To ensure consistency with prior work (Adise et al., 2021), an Elastic Net regression framework (Zou & Hastie, 2005) was used to predict TMI from multiple sets of imaging features (i.e., sMRI, DTI, and resting-state fMRI). Five models were defined: 1) a base model with covariates only (sex, ethnicity, PDS and age); 2) rs-fMRI model with Gordon cortical correlations and subcortical correlations, plus covariates; 3) LEiDA model including LEiDA-derived features restricted to states significantly associated with intrinsic functional networks (IFNs; Yeo et al., 2011), plus covariates; 4) FCH model with measures derived from the first 20 functional connectome harmonics (FCHs), plus covariates; and finally the 5) combined model including all imaging-derived features together with covariates. Analyses were performed on both the original datasets and harmonized datasets, the latter adjusted to reduce site-specific variability. Elastic Net regression combines L1 (Lasso) and L2 (Ridge) regularization penalties. Hyperparameters were optimized using 10-fold cross-validation to determine the optimal L1–L2 balance (l1_ratio ∈ [0.01, 0.025, 0.05, 0.1, 0.5, 0.7, 0.9, 1]) and regularization strength (α ∈ [2⁻⁸, 2⁷]). The maximum number of iterations was set to 100 to limit model complexity and mitigate convergence issues associated with high-dimensional resting-state fMRI features. Model performance was evaluated using 5-fold cross-validation, with approximately 80% of the data used for training and 20% for testing in each fold.

Performance metrics included the coefficient of determination (R²) and negative mean absolute error (negMAE), computed separately for training and test sets. All analyses were implemented in Python 3.9 using scikit-learn (v1.2.2) (Pedregosa et al., 2011).

## Data Availability

Anonymized data from the Adolescent Brain Cognitive Development (ABCD) Study are released annually and made available to the research community. Information on accessing ABCD data through the National Data Archive (NDA) is provided on the ABCD Study data-sharing website (https://abcdstudy.org/scientists_data_sharing.html), with instructions for creating an NDA study available at https://nda.nih.gov/training/modules/study.html. The ABCD data repository is dynamic and continues to expand over time.

The data used in this study were obtained from the ABCD Study (add specific dataset URL). Raw data are accessible via the NDA (add specific URL).

## Code Availability

All code used in the present study are openly available via a public GitHub repository (https://github.com/isawig/ABCD_brain-dynamics_analyses). Bash scripts were used for data download, preprocessing, and for the preparation of analysis pipelines and inputs for LEiDA. MATLAB code was used to perform LEiDA and harmonic modeling and analyses. Python scripts were employed to implement elastic net models, and R scripts were used for other statistical analyses.

## Acknowledgments

The authors wish to acknowledge CSC – IT Center for Science, Finland, for generous computational resources. The ABCD Study is supported by the NIH and National Institute on Drug Abuse and additional federal partners under award Numbers U01DA041022, U01DA041025, U01DA041028, U01DA041048, U01DA041089, U01DA041093, U01DA041106, U01DA041117, U01DA041120, U01DA041134, U01DA041148, U01DA041156, U01DA041174, U24DA041123, U24DA041147, U01DA050987, U01DA050988, U01DA050989, U01DA050988, U01DA051018, U01DA051037, U01DA051038, and U01DA051039. A full list of supporters is available at https://abcdstudy.org/about/federal-partners/. A listing of participating sites and a complete listing of the study investigators can be found at https://abcdstudy.org/principal-investigators/. This work was supported by the Sigrid Jusélius Foundation, Emil Aaltonen Foundation, Finnish Medical Foundation, Alfred Kordelin Foundation, Juho Vainio Foundation, Turku University Foundation, Hospital District of Southwest Finland, State Grants for Clinical Research (ERVA), Orion Research Foundation, Signe and Ane Gyllenberg Foundation, Yrjö Jahnsson Foundation, Jalmari and Rauha Ahokas foundation, Päivikki and Sakari Sohlberg foundation, Finnish Cultural foundation, University of Turku Graduate School and Finnish Brain Foundation. JC was supported by LARSyS FCT funding (10.54499/LA/P/0083/2020, 10.54499/UIDP/50009/2020 and 10.54499/UIDB/50009/2020), Portugal.

## Author contributions

IMW - Conceptualization, Methodology, Data curation, Formal analysis, Supervision, Writing - review & editing

AB - Data curation, Formal analysis, Review & editing IS - Data curation, Review & editing

AJ - Designed and implemented the HPC pre-processing pipeline on CSC Puhti, including development and execution of Bash and SLURM scripts for fMRIPrep and XCP-D processing of all participants.

RL - Data curation, Review & editing

EPP; HM; AR; AB; SL; MP; MV; HK; LK; JRBC; MLK; SA; JS; RAIB- Review & editing

JJT - Conceptualization, Resources, Data curation, Funding acquisition, Review & editing

