## Supplementary Material for "Functional connectome harmonics and dynamic connectivity maps of the preadolescent brain"

2025-12-22

### Abstract

The complexity of the human brain arises from its highly organized topography, where functionally specialized yet interconnected regions form dynamic networks that support cognition and behavior. During preadolescence, these networks undergo maturation, offering a critical window for characterizing large-scale functional organization. Using resting-state fMRI data from 11,868 participants aged 9–10 years in the Adolescent Brain Cognitive Development (ABCD) Study, we applied Functional Connectome Harmonics (FCH) and Leading Eigenvector Dynamics Analysis (LEiDA) to quantify spatial gradients and dynamic network states. FCH revealed brain gradient patterns capturing hierarchical organization across cortical and subcortical areas, while LEiDA identified recurrent phase-locked brain states aligned with canonical intrinsic connectivity networks. We cross-validated spatial and temporal components, examined associations with demographic and developmental variables, and harmonized data across acquisition sites. Together, these findings establish the first large-scale spatiotemporal reference framework for preadolescent functional brain organization, providing a foundation for tracking neurodevelopmental trajectories and early neural markers of adolescent health.

### Introduction

This document provides additional information and results that complement those presented in the main manuscript, with the aim of guiding the reader toward a deeper understanding of the methods, analyses, and findings.

The first chapter, **Methods**, offers an expanded description of the participant samples. In particular, it includes a detailed comparison between the final sample used for statistical analyses and the excluded subjects, supported by additional distribution plots illustrating differences between the two datasets. This chapter also presents the complete list of regions of interest (ROIs) composing the atlas used in the study, together with their correspondence to reference intrinsic functional networks (IFNs) as defined by Yeo et al. (2011). Finally, a comprehensive overview of the two main connectome approaches employed—Functional Connectome Harmonics (FCH) and Leading Eigenvector Dynamics Analysis (LEiDA)—is provided, including their theoretical background and detailed descriptions of the computational procedures.

The second chapter, **Results**, reports additional findings from the FCH and LEiDA connectome analyses, as well as from the statistical analyses and elastic net models. These include results obtained using different FCH parameter settings (varying the number of nearest neighbors), a more detailed characterization of the correspondence between LEiDA states and IFNs, and partial least squares regression (PLSR) results for LEiDA states 2 to 6 (K = 6), complementing the results for state 1 reported in the main text. Furthermore, this chapter assesses site-wise effects, providing a clear demonstration of the impact of data harmonization on FCH and LEiDA metrics. Finally, additional plots related to the statistical analyses are presented for the original (non-harmonized) dataset, complementing those shown for the harmonized data in the main manuscript. For the partial correlation analyses, distribution plots, heatmaps, and scatterplots are reported for each significant association in both datasets, offering a more detailed view of the main results.

### 1 Methods

#### 1.1 Participants

**Table 1: Comparison of included and excluded participants on key covariates.** Table summarizing the differences between participants included in the final analyses (N = 4,453) and those excluded (N = 7,058). Only subjects with complete data for all required covariates were considered in the excluded group. Welch's t-tests were used for continuous variables (interview age, pubertal developmental scale (PDS), triponderal mass index (TMI)), and Chi-squared tests for categorical variables (sex, ethnicity, site). Test statistics and p-values are reported, with significance levels below 0.001 indicated as p < 0.001.

| **Variable** | **Value** | **Included (N = 4453)** | **Excluded (N = 7058)** | **t** | **X²** | **p** |
|---|---|---|---|---|---|---|
| | | | *N (%) or M (SD)* | | | |
| Sex | | | | | 5.7383 | 0.0166 |
| | *Males* | 2272 (51.02%) | 3764 (53.33%) | | | |
| | *Females* | 2181 (48.98%) | 3294 (46.67%) | | | |
| Ethnicity | | | | | 106.2 | < 0.001 |
| | *White* | 2561 (57.51%) | 3512 (49.76%) | | | |
| | *Black* | 491 (11.03%) | 1182 (16.75%) | | | |
| | *Hispanic* | 905 (20.32%) | 1409 (19.96%) | | | |
| | *Asian* | 76 (1.71%) | 167 (2.37%) | | | |
| | *Other* | 420 (9.43%) | 788 (11.16%) | | | |
| Site | | | | | 964.48 | < 0.001 |
| | *G010* | 82 (1.84%) | 261 (3.70%) | | | |
| | *G031* | 212 (4.76%) | 508 (7.20%) | | | |
| | *G032* | 254 (5.70%) | 478 (6.77%) | | | |
| | *G075* | 183 (4.11%) | 526 (7.45%) | | | |
| | *G087* | 69 (1.55%) | 311 (4.40%) | | | |
| | *P023* | 42 (0.94%) | 496 (7.03%) | | | |
| | *P043* | 124 (2.78%) | 444 (6.29%) | | | |
| | *P064* | 85 (1.91%) | 295 (4.18%) | | | |
| | *S011* | 313 (7.03%) | 289 (4.09%) | | | |
| | *S012* | 239 (5.37%) | 345 (4.89%) | | | |
| | *S013* | 128 (2.87%) | 289 (4.09%) | | | |
| | *S014* | 634 (14.24%) | 364 (5.16%) | | | |
| | *S020* | 257 (5.77%) | 339 (4.80%) | | | |
| | *S021* | 270 (6.06%) | 411 (5.82%) | | | |
| | *S022* | 250 (5.61%) | 301 (4.26%) | | | |
| | *S042* | 308 (6.92%) | 261 (3.70%) | | | |
| | *S053* | 262 (5.88%) | 319 (4.52%) | | | |
| | *S065* | 219 (4.92%) | 206 (2.92%) | | | |
| | *S076* | 208 (4.67%) | 232 (3.29%) | | | |
| | *S086* | 176 (3.95%) | 190 (2.69%) | | | |
| | *S090* | 138 (3.10%) | 193 (2.73%) | | | |
| Interview age | | 119.08 (7.55) | 118.91 (7.47) | 1.1366 | | 0.2557 |
| PDS | | 1.88 (0.75) | 1.91 (0.76) | -2.0365 | | 0.04173 |
| TMI | | 13.35 (2.61) | 13.33 (2.75) | 0.33377 | | 0.7386 |

To evaluate the representativeness of the final sample relative to the broader ABCD cohort, we compared included (N = 4,453) and excluded participants (N = 7,058; after removal of subjects with missing data in any of the six covariates). Group differences were assessed using Chi-squared tests for categorical variables (sex, ethnicity, site) and Welch's t-tests for continuous variables (age at interview, PDS, TMI).

Distributions of all covariates for included and excluded participants are shown in Figure 1 to Figure 6. Categorical variables are reported as percentages within each group, whereas continuous variables are visualized using density plots.

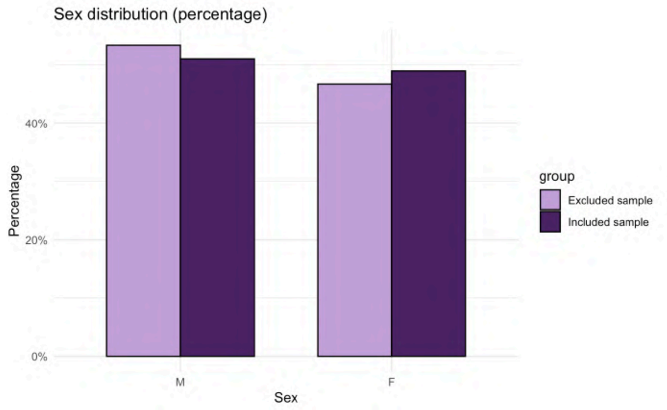

**Figure 1: Distribution of variable sex.** Distribution of sex categories (M = male, F = female) in the two datasets. The y-axis shows the percentage of subjects belonging to each category within each dataset (N = 4,453 included and N = 7,058 excluded subjects).

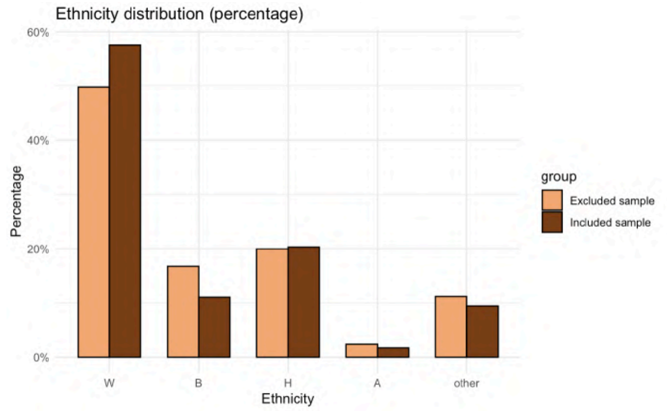

**Figure 2: Distribution of variable ethnicity.** Distribution of ethnicity categories (W = white, B = black, H = hispanic, A = asian, other) in the two datasets. The y-axis shows the percentage of subjects belonging to each category within each dataset (N = 4,453 included and N = 7,058 excluded subjects).

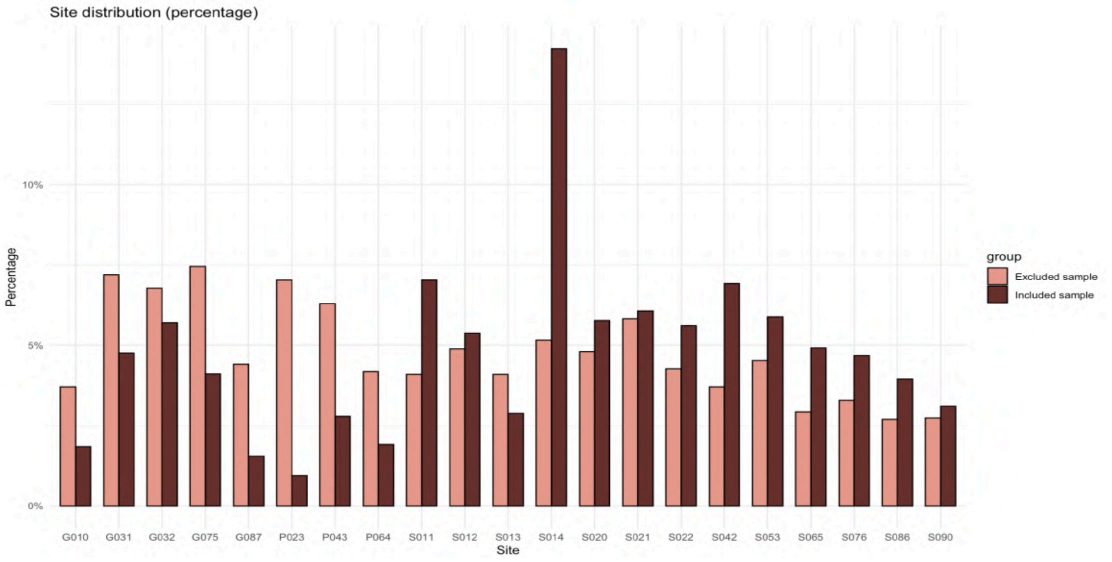

**Figure 3: Distribution of variable site.** Distribution of site categories in the two datasets. The y-axis shows the percentage of subjects belonging to each category within each dataset (N = 4,453 included and N = 7,058 excluded subjects).

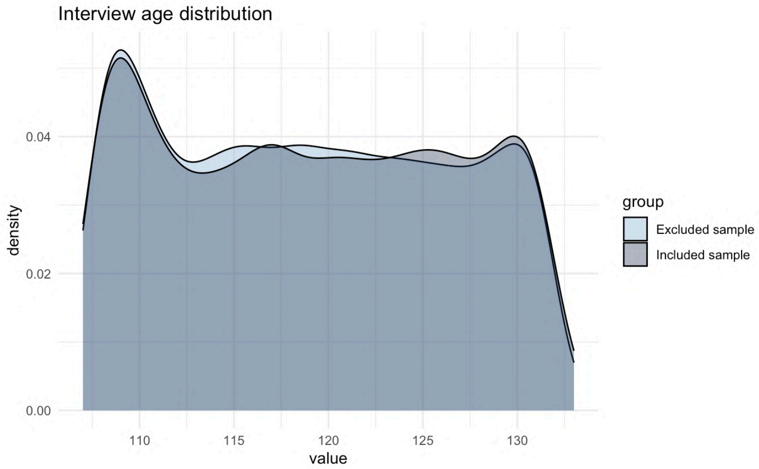

**Figure 4: Distribution of interview age.** Density plot showing the distribution of the variable interview age in the two datasets (N = 4,453 included and N = 7,058 excluded subjects).

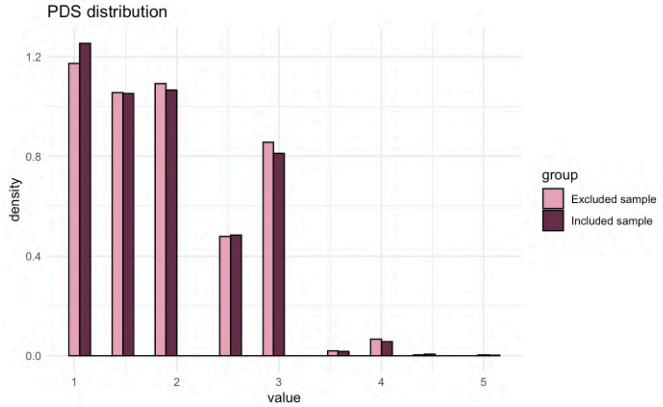

**Figure 5: Distribution of pubertal developmental scale (PDS).** Histogram showing the distribution of the discrete PDS variable in the two datasets (N = 4,453 included and N = 7,058 excluded subjects).

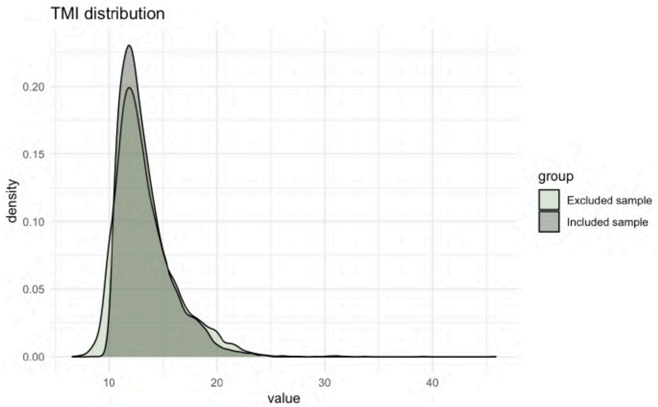

**Figure 6: Distribution of triponderal mass index (TMI).** Density plot showing the distribution of the variable TMI in the two datasets (N = 4,453 included and N = 7,058 excluded subjects).

For TMI comparison, subjects with abnormal TMI values were excluded, defined as those outside the range $Q1 - 1.5 \times \text{IQR} < \text{TMI} < Q3 + 3 \times \text{IQR}$, where $Q1$ and $Q3$ are the first and third quartiles, respectively, and $\text{IQR} = Q3 - Q1$ is the interquartile range. This procedure was performed consistently with previous studies on ABCD data involving TMI analyses (Ru Li et al., 2025).

Overall, although some significant differences were observed between included and excluded subjects, particularly for site, and to a lesser extent for sex, ethnicity, and PDS, the final analytical sample remains broadly representative of the initial ABCD cohort. Continuous variables such as interview age and TMI showed minimal but non-significant differences between groups. The largest discrepancies were observed across sites, likely reflecting site-specific differences in fMRI data quality and subsequent exclusions.

#### 1.2 Regions of Interest (ROIs) and network assignment

Initial acquisitions were sampled using the 4S156 reference atlas, which included 100 cortical Regions of Interest (ROIs) from the Schaefer100 atlas and 56 subcortical ROIs from the COI168, ThalamusHCP, and SubcorticalHCP atlases. Subsequently, ROIs were relabeled using a modified atlas composed of the same 100 cortical ROIs from Schaefer100, along with 14 subcortical ROIs obtained by averaging preselected subcortical regions to meet ENIGMA Toolbox requirements. The 14 subcortical ROIs included bilateral accumbens, amygdala, caudate, hippocampus, pallidum, putamen, and thalamus.

The complete list of 114 ROIs used in the analysis, is provided below. Cortical ROIs, named accordingly to Schaefer100 atlas, are assigned to the seven canonical Yeo networks (Visual, Somatomotor, Dorsal Attention, Ventral Attention, Limbic, Frontoparietal, and Default Mode; Yeo et al.,2011), while subcortical ROIs are labeled according to anatomical structures. Each ROI is represented in both the left (LH) and right (RH) hemisphere.

**Table 2: Cortical and Subcortical ROIs with Yeo Networks.** Summary of 114 ROIs included in connectome analysis with specified Yeo reference RSN.

| ROI Name | Type | Yeo Network |
|---|---|---|
| Vis 1 | Cortical | Visual |
| Vis 2 | Cortical | Visual |
| Vis 3 | Cortical | Visual |
| Vis 4 | Cortical | Visual |
| Vis 5 | Cortical | Visual |
| Vis 6 | Cortical | Visual |
| Vis 7 | Cortical | Visual |
| Vis 8 | Cortical | Visual |
| Vis 9 | Cortical | Visual |
| SomMot 1 | Cortical | Somatomotor |
| SomMot 2 | Cortical | Somatomotor |
| SomMot 3 | Cortical | Somatomotor |
| SomMot 4 | Cortical | Somatomotor |
| SomMot 5 | Cortical | Somatomotor |
| SomMot 6 | Cortical | Somatomotor |
| DorsAttn Post 1 | Cortical | Dorsal Attention |
| DorsAttn Post 2 | Cortical | Dorsal Attention |
| DorsAttn Post 3 | Cortical | Dorsal Attention |
| DorsAttn Post 4 | Cortical | Dorsal Attention |
| DorsAttn Post 5 | Cortical | Dorsal Attention |
| DorsAttn Post 6 | Cortical | Dorsal Attention |
| DorsAttn PrCv 1 | Cortical | Dorsal Attention |
| DorsAttn FEF 1 | Cortical | Dorsal Attention |
| SalVentAttn ParOper 1 | Cortical | Salience Ventral Attention |
| SalVentAttn FrOperIns 1 | Cortical | Salience Ventral Attention |

| ROI Name | Type | Yeo Network |
|---|---|---|
| SalVentAttn FrOperIns 2 | Cortical | Salience Ventral Attention |
| SalVentAttn PFCl 1 | Cortical | Salience Ventral Attention |
| SalVentAttn Med 1 | Cortical | Salience Ventral Attention |
| SalVentAttn Med 2 | Cortical | Salience Ventral Attention |
| SalVentAttn Med 3 | Cortical | Salience Ventral Attention |
| Limbic OFC 1 | Cortical | Limbic |
| Limbic TempPole 1 | Cortical | Limbic |
| Limbic TempPole 2 | Cortical | Limbic |
| Cont Par 1 | Cortical | Frontoparietal |
| Cont PFCl 1 | Cortical | Frontoparietal |
| Cont pCun 1 | Cortical | Frontoparietal |
| Cont Cing 1 | Cortical | Frontoparietal |
| Default Temp 1 | Cortical | Default Mode Network |
| Default Temp 2 | Cortical | Default Mode Network |
| Default Par 1 | Cortical | Default Mode Network |
| Default Par 2 | Cortical | Default Mode Network |
| Default PFC 1 | Cortical | Default Mode Network |
| Default PFC 2 | Cortical | Default Mode Network |
| Default PFC 3 | Cortical | Default Mode Network |
| Default PFC 4 | Cortical | Default Mode Network |
| Default PFC 5 | Cortical | Default Mode Network |
| Default PFC 6 | Cortical | Default Mode Network |
| Default PFC 7 | Cortical | Default Mode Network |
| Default pCunPCC 1 | Cortical | Default Mode Network |
| Default pCunPCC 2 | Cortical | Default Mode Network |
| Accumbens | Subcortical | - |
| Amygdala | Subcortical | - |
| Caudate | Subcortical | - |
| Hippocampus | Subcortical | - |
| Pallidum | Subcortical | - |
| Putamen | Subcortical | - |
| Thalamus | Subcortical | - |

#### 1.3 Functional Connectome Harmonics

##### 1.3.1 Method overview

FCH extend the classical Fourier basis to the human connectome, offering a frequency-specific description of cortical activity under the hypothesis that RSNs follow harmonic wave patterns at distinct frequencies. The following Figure 7 provides an overview of the FCH framework, illustrating the successive steps from preprocessing to the identification of connectome harmonics. Each step will be described in detail in the following sections.

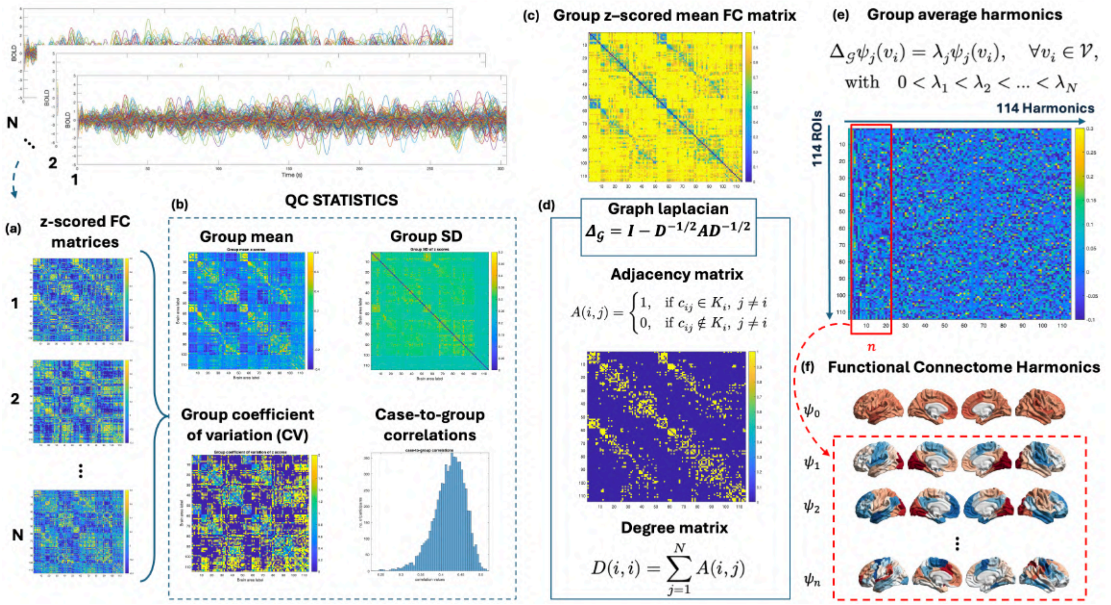

**Figure 7: Overview of the pipeline used to derive functional connectome harmonics from resting-state fMRI data from 6,624 preadolescents. a**, Individual functional connectivity (FC) matrices are computed from BOLD time series and z-scored to standardize across participants. **b**, Group-level mean and standard deviation (SD) of z-scored FC values are calculated, together with the coefficient of variation (CV) and case-to-group correlations, to assess inter-individual consistency. **c**, Z-scored time series from all participants are concatenated and pairwise correlations between brain regions are computed to obtain a group-level FC matrix. **d**, Adjacency and degree matrices are derived from the group-level FC matrix to construct the graph Laplacian, providing a discrete approximation of the Laplacian operator. **e**, Eigen-decomposition of the graph Laplacian yields harmonic modes ordered by increasing spatial frequency, corresponding to principal axes of functional organization; the first n harmonics are associated with the lowest eigenvalues. **f**, The resulting eigenmodes are projected onto cortical surfaces, revealing spatially distributed harmonic patterns across the brain. The first harmonic, which exhibits a spatially uniform pattern, is excluded from subsequent analyses. QC = quality check; FC = functional connectivity.

The human connectome can be represented as a graph $\mathcal{G}$, where vertices sampled from the grey matter surface are represented by nodes $\mathcal{V} = \{v_i \mid i = 1, ..., N\}$, with $N$ the total number of nodes, and the connections between vertices are represented as edges $\mathcal{E} = \{e_{ij} \mid (v_i, v_j) \in \mathcal{V} \times \mathcal{V}\}$, such that the graph is $\mathcal{G} = (\mathcal{V}, \mathcal{E})$. From the definition of the graph structure we can formulate the adjacency matrix $A$.

In this study, the graph was derived from a group-average functional connectivity (FC) matrix.

First, individual FC matrices were averaged across participants, and then, for each node, only the $k$ strongest correlations were retained to construct a symmetric, undirected adjacency matrix. In this way, each node represents a brain region (ROI) from the atlas, and edges reflect the strength of functional coupling between regions. Therefore, the graph structure is represented by the $N \times N$ adjacency matrix $A$, from which it is possible to define also the degree matrix $D$, both defined as in panel d of Figure 7.

From the graph $\mathcal{G} = (\mathcal{V}, \mathcal{E})$, the symmetric graph Laplacian $\Delta_{\mathcal{G}}$ is computed to obtain a discrete analogue of the Laplace operator $\Delta$ applied to the human connectome. Several formulations of the graph Laplacian exist, differing in their normalization and spectral properties. In the present study, we adopted the symmetric normalized Laplacian which captures the connectivity structure of the graph by penalizing connections based on node degree (panel d, Figure 7).

Finally, connectome harmonics $\psi_j$, $j = 1, ..., N$, are obtained by solving the eigenvalue problem shown in panel e of Figure 7, where $\lambda_j$ are the eigenvalues of $\Delta_{\mathcal{G}}$, and $\psi_j$ are the corresponding eigenvectors, representing the connectome harmonics. Derived connectome harmonics have are all orthonormal, i.e. $||\psi_k|| = 1, \forall k$.

##### 1.3.2 Metrics to characterize FCHs

To compare brain activity with the connectome harmonic basis, fMRI data are first mapped onto the cortical surface, yielding the spatiotemporal signal $\mathcal{F}(v,t)$ for all vertices $v \in \mathcal{V}$ and time points $t \in [1, ..., T]$.

At each time point $t_i$, the spatial pattern of cortical activity $\mathcal{F}_{t_i}(v)$ can be expressed as a linear combination of connectome harmonics $\Psi = \{\psi_k\}_{K=1}^{N}$:

$$\mathcal{F}_{t_i} = \alpha_1(t_i)\psi_1 + \alpha_2(t_i)\psi_2 + ... + \alpha_N(t_i)\psi_N = \sum_{K=1}^{N} \alpha_k(t_i)\psi_k \tag{1}$$

where the temporal activity $\alpha_k(t)$ of each connectome harmonic $\psi_k$ was estimated by projection of $\mathcal{F}(v,t)$ onto that particular harmonic. The coefficients $\alpha_k$ are estimated as:

$$\alpha_k(t) = \langle \mathcal{F}_t, \psi_k \rangle \tag{2}$$

Where $k$ denotes the spatial frequency of the harmonic and $t$ the specific moment in time. From the computed harmonics it is possible to derive additional metrics to characterize FCHs results to provide an index of total information flow across the identified spatial pattern.

###### 1.3.2.1 Power of an harmonic brain state

The **power** of an harmonic brain state reflects the strength of its activation at a given time point in the fMRI data.

For each connectome harmonic $\psi_k$, with $k \in [1, ..., N]$, the power at time $t$ is computed as:

$$\mathrm{P}(\psi_k, t) = |\alpha_k(t)| \tag{3}$$

This measure quantifies how strongly the cortical activity at a specific moment projects onto a given harmonic.

The **total power** of a harmonic state is obtained by summing its contributions across all time points. Because power is defined as the dot product between the cortical activity pattern and an orthonormal connectome harmonic, its possible range is constrained: the minimum value is always 0 (no projection onto that harmonic); while the maximum value depends on the magnitude of the cortical activity pattern at that time point.

###### 1.3.2.2 Energy of an harmonic brain state

The **energy** of a connectome harmonic represents its frequency-weighted contribution to cortical dynamics. It combines the instantaneous power of a harmonic with its intrinsic frequency, captured by the eigenvalue $\lambda_k$.

Formally, the energy of a harmonic brain state $\psi_k$ at time $t$ is defined as:

$$\mathrm{E}(\psi_k, t) = |\alpha_k(t)|^2 \lambda_k^2 \tag{4}$$

While the **total energy** of brain activity for any given time point $t$ is given by:

$$\mathrm{E}_{total}(t) = \sum_{K=1}^{N} |\alpha_k(t)|^2 \lambda_k^2 = ||\Delta \mathcal{F}_t(v)||^2 \tag{5}$$

Since the Laplace operator $\Delta$ captures the flow of activity across the network, the total energy provides an index of the overall flow of neural activity across the cortex at that specific time. In other words, energy quantifies how dynamic and spatially distributed the cortical activity is, weighted by the harmonic frequencies.

#### 1.4 Leading Eigenvector Dynamics Analysis

##### 1.4.1 Method overview

Leading Eigenvector Dynamics Analysis (LEiDA) is a data-driven method for studying dynamic functional connectivity (dFC). It captures recurrent brain states by extracting the leading eigenvector of phase coherence matrices at each time point, thus providing a low-dimensional representation of whole-brain dynamics. This approach enables characterization of the temporal evolution of connectivity patterns while being less sensitive to high-frequency noise compared to traditional methods.

The following Figure 8 provides an overview of the LEiDA framework, illustrating the successive steps from preprocessing to the identification of recurrent phase-locking states. Each step will be described in detail in the following sections.

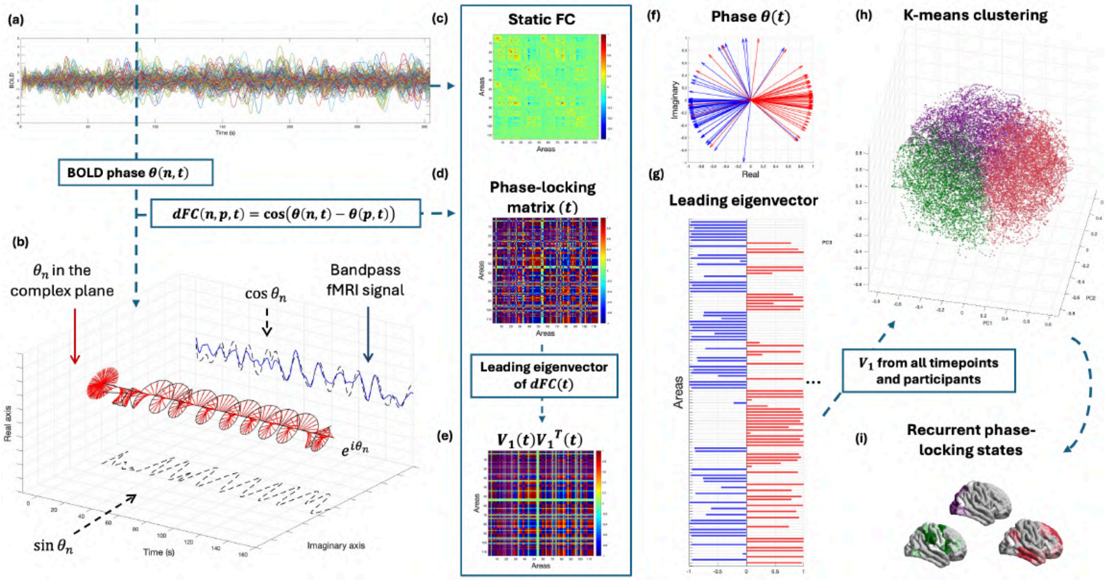

**Figure 8: Overview of Leading Eigenvector Dynamics Analysis (LEiDA) applied to resting-state fMRI data from 6,624 preadolescents.** **a**, Resting-state BOLD time series from a representative participant across N = 114 brain regions. **b**, BOLD signals are band-pass filtered (0.01–0.1 Hz) and transformed into analytic signals using the Hilbert transform to extract instantaneous phases. **c**, Static functional connectivity (FC) is computed as pairwise correlations between BOLD signals across the entire scan. **d**, Dynamic functional connectivity (dFC) is estimated at each time point using phase coherence, defined as the cosine of phase differences between all pairs of regions, yielding a time-resolved phase-locking (PL) matrix. **e**, The leading eigenvector, V (t), captures the dominant connectivity pattern of dFC(t) and allows reconstruction of the principal PL configuration via the outer product. **f**, Instantaneous phases across all regions at a single repetition time are represented in the complex plane; arrow orientation and color indicate the sign of the elements of V (t) (red, positive; blue, negative). **g**, V (t) summarizes the dominant phase-alignment pattern, with regions oscillating in phase or anti-phase relative to the global configuration. **h**, Leading eigenvectors from all time points and participants are concatenated and clustered using k-means to identify k recurrent PL states; centroids for k = 3 are shown. **i**, PL state centroids are projected onto the cortical surface, revealing distinct large-scale phase-locking configurations across the brain. FC = functional connectivity.

BOLD Phase Coherence Connectivity can be used to compute a time-resolved dynamic functional connectivity (dFC) matrix of size $N \times N \times T$, where $N$ is the number of brain regions (ROIs) and $T$ is the number of recording frames. To compute the phase coherence, first the BOLD phase $\theta(n,t)$ of each ROI $n$ is estimated using the Hilbert transform, allowing the regional signal $x(t)$ to be represented as:

$$x(t) = A(t) \cdot \cos(\theta(t)) \tag{6}$$

where $A(t)$ and $\theta(t)$ denote the instantaneous amplitude and phase, respectively. Given the phase of the BOLD signals, the phase coherence between brain areas $n$ and $m$ at time $t$ is defined as in second blue panel close to d one in Figure 8.

Instead of comparing the full dFC matrices across time, LEiDA considers only the **leading eigenvector** $V_1(t)$ (size $N \times 1$) of each dFC matrix, capturing the dominant connectivity pattern at time $t$. Considering only the leading eigenvector of the dFC and neglecting the patterns captured by the remaining eigenvectors is justified by the fact that the leading eigenvector always represent more than half of the variance associated to the overall dFC. The first eigenvector $V_1(t)$ represents the BOLD phase-locking pattern at each time point. Since dFC($t$) is symmetric, its eigenvectors are orthogonal ($V^{-1}(t) = V^T(t)$) and the eigenvalues are real. The dominant connectivity pattern can be reconstructed via the outer product $V_1(t)V_1^T(t)$, reducing the dimensionality from $N(N-1)/2$ to $N$ while retaining most variance (panel e, Figure 8).

To identify recurrent FC patterns, **k-means clustering** (panels h and i in Figure 8) is applied to the leading eigenvectors $V_1(t)$ across time points and subjects. The algorithm iteratively finds $k$ cluster centroids that minimize the distance to the assigned observations. The output is $k$ centroids $V_{C_\alpha}$ (each of size $N \times 1$), representing distinct BOLD phase-locking states $R^\alpha$ with $\alpha = 1, ..., k$, where fMRI time point is assigned to the closest centroid.

##### 1.4.2 Metrics to characterize LEiDA

Following the partition of the trajectory into $k$ PL states over time $t$, we computed additional metrics to characterize their temporal properties: Fractional Occupancy (FO), Dwell Time (DT), and Markov chain transition probabilities. These measures quantify, respectively, the prevalence of each state, the average duration of its visits, and the likelihood of transitioning between states during the fMRI acquisition.

###### 1.4.2.1 Fractional Occupancy

**Fractional Occupancy** is defined as the number of times when the trajectory $\underline{x}(t)$ is assigned to each of the defined clusters $R^\alpha$ divided by the total number of time points $T$, assuming stationarity within each scan, since subjects are constantly in resting-state. It can be formulated as follows:

$$\Pi_\alpha^{(S)} = \frac{1}{T} \sum_{t=1}^{T} \chi[\underline{x}(t) \in R^\alpha] \tag{7}$$

where $\chi$ is the indicator function, i.e. $\chi(A) = 1$ if the event $A$ is true, $\chi(A) = 0$ otherwise and $T$ is the number of timepoints corresponding to each fMRI scan ($S$).

###### 1.4.2.2 Dwell Time

**Dwell Time** is defined as the mean of all the consecutive periods of each state, and is used to describe the average time periods when a given PL state $\alpha$ is being visited in each fMRI scan $S$. It can be formulated as follows:

$$DT_{\alpha}^{(S)} = \frac{1}{p_{\alpha}} \sum_{1}^{p_{\alpha}} C_{p_{\alpha}} \tag{8}$$

where $DT_{\alpha}$ is the Dwell Time of PL state $\alpha$, $p_{\alpha}$ is the number of consecutive periods assigned to PL state $\alpha$ and $C_{p_{\alpha}}$ is the duration of each consecutive period.

###### 1.4.2.3 Markov Chain transition probabilities

**Markov Chain transition probability** matrix $W_{\alpha\beta}^{(S)}$ of each fMRI scan $S$ define the transition matrix from state $\alpha$ to state $\beta$, i.e.:

$$W_{\alpha\beta}^{(S)} = P[\underline{x}(t+1) \in R^{\beta} | \underline{x}(t) \in R^{\alpha}] = \frac{\Pi_{\alpha\beta}^{(S)}}{\Pi_{\alpha}^{(S)}} \tag{9}$$

Where $\Pi_{\alpha\beta}$ defines the probability of being in the PL state $\alpha$ at time $t$ and in the PL state $\beta$ at time $t+1$, i.e.:

$$\Pi_{\alpha\beta}^{(S)} = \frac{1}{T-1} \sum_{t=1}^{T-1} \chi[\underline{x}(t) \in R^{\alpha}, \underline{x}(t+1) \in R^{\beta}] \tag{10}$$

The obtained transition matrix defines an homogeneous Markov Chain, characterizing the transition between BOLD phase locking states, and is estimated separately for each scan $S$. A Markov Chain models a stochastic process in which the probability of moving to a given state depends only on the current one, providing a compact framework to describe temporal dependencies between brain states.

### 2 Results

#### 2.1 Functional Connectome Harmonics

##### 2.1.1 Robustness of FCHs across nearest neighbors

FCHs were estimated using different values of nearest neighbors ($nn = 5, 10, 15, 20$) to assess the robustness of the results across parameters. Here we report results for the other values of different neighbors chosen ($nn = 5, 15, 20$), visualizing the first six harmonics ($\phi_1$, $\phi_2$, ..., $\phi_6$), spanning both cortical and subcortical regions. As before, the first functional harmonic, associated with eigenvalue 0.

Spatial patterns of connectome harmonics are consistent across all nearest-neighbor solutions, showing similar distributions of regional gradients regardless of the chosen $nn$ value. However, polarity inversions can occur between different $nn$ configurations, that is, regions showing positive values in one case may display negative values in another. Note that the polarity of each functional harmonic is arbitrary: if $\phi$ is an eigenvector, $-\phi$ represents the same spatial pattern with inverted signs. For this reason, differences in positive versus negative regions across $nn$ values reflect this arbitrary choice of sign rather than substantive changes in the underlying spatial organization. Therefore, the same functional networks identified for $nn = 10$ also emerge across other $nn$ values, confirming the robustness of the results to this parameter choice.

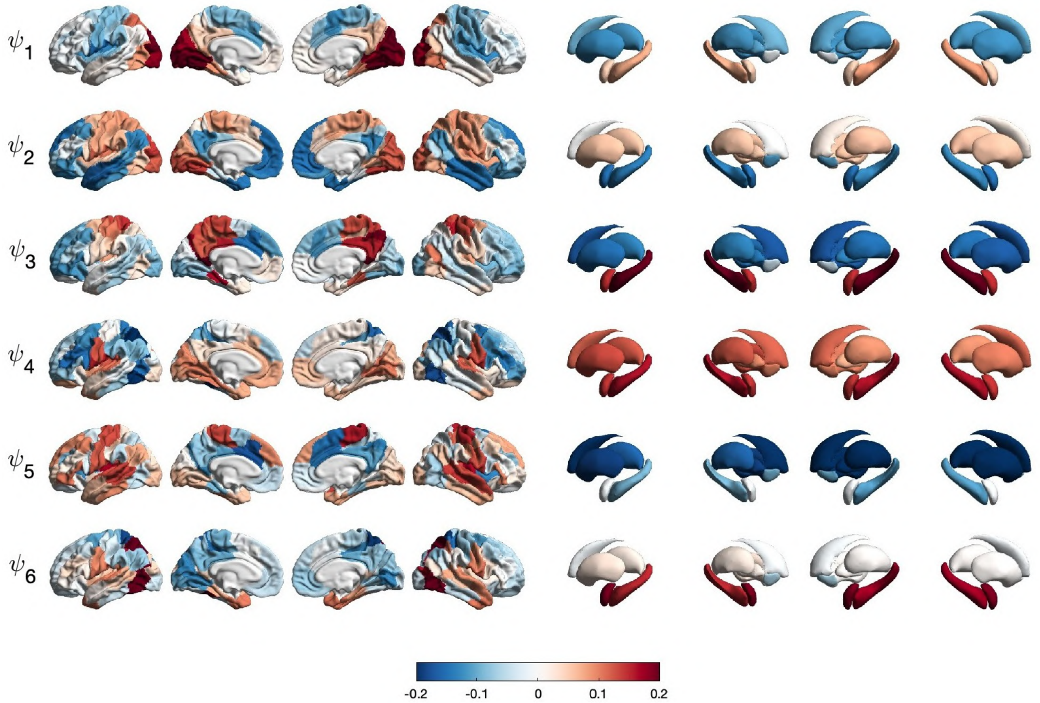

**Figure 9: First six FCHs - nn = 5.** Spatial distribution of the first six harmonics, mapped onto cortical and subcortical surfaces and estimated using nn = 5. The first harmonic, associated to eigenvalue 0, was excluded due to lack of anatomical specificity (N = 6,624).

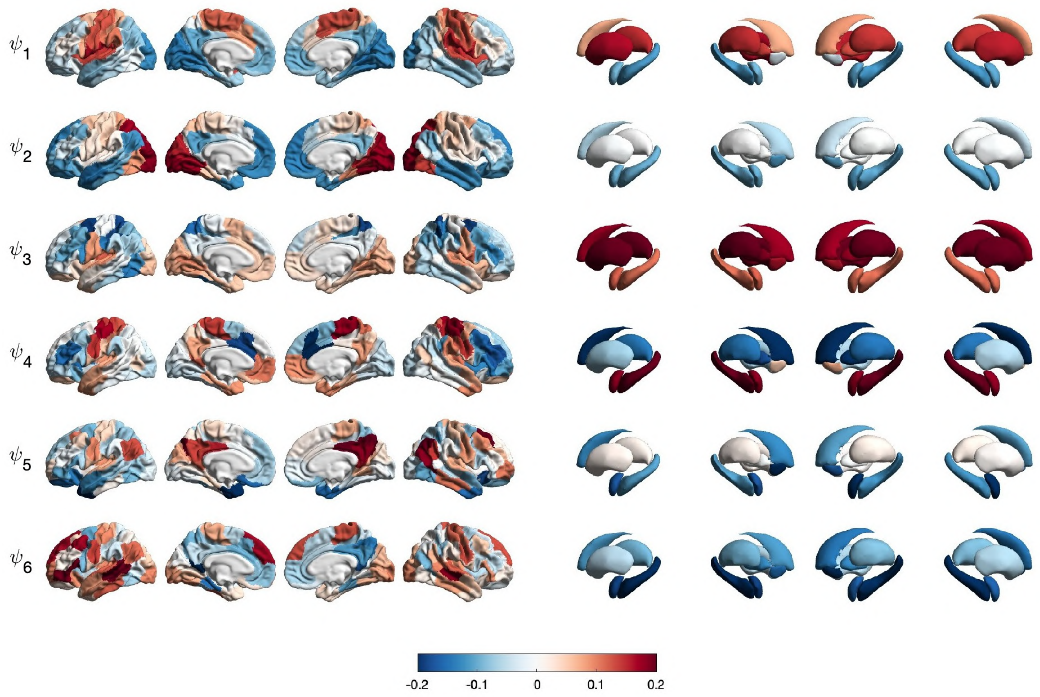

**Figure 10: First six FCHs - nn = 15.** Same procedure as in figure 9 (N = 6,624).

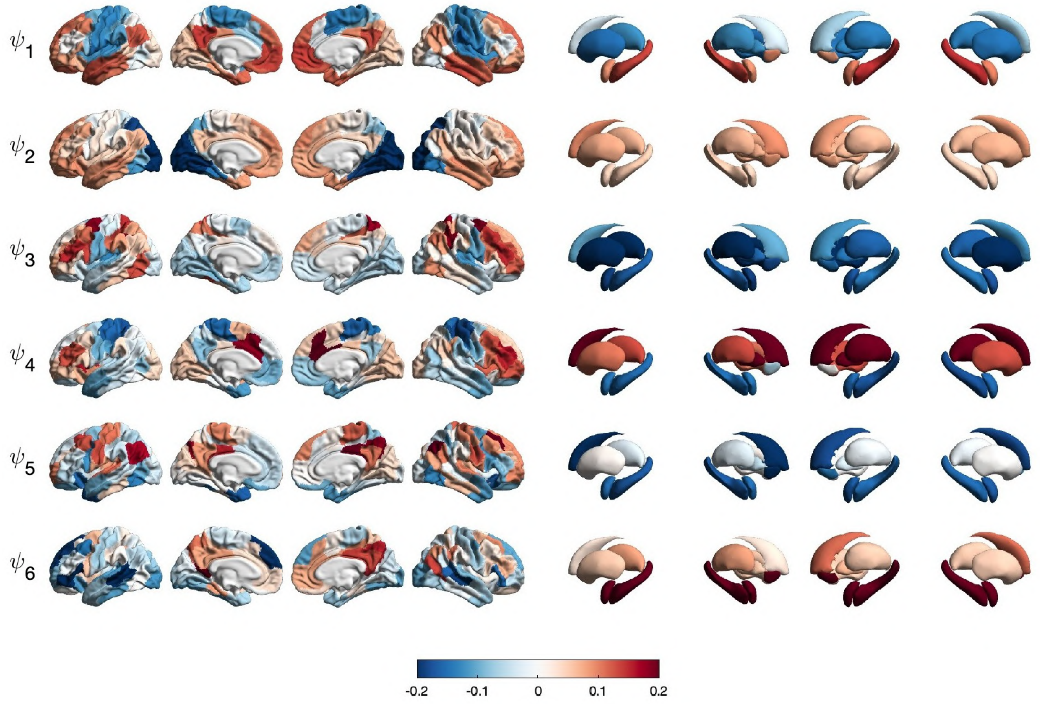

**Figure 11: First six FCHs - nn = 20.** Same procedure as in figure 9 (N = 6,624).

##### 2.1.2 First twenty FCHs spatial patterns

Since statistical analysis is carried out considering the first twenty harmonics, here we visualized them on the cortical surface following the procedure already described.

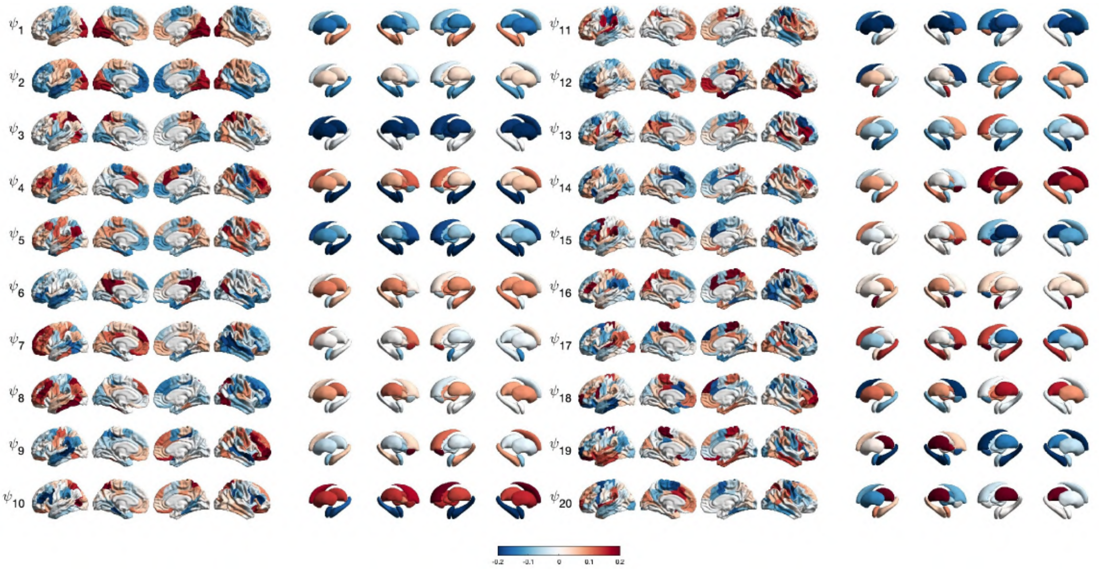

**Figure 12: First twenty Functional Connectome Harmonics (FCH).** Spatial distribution of the first twenty harmonics mapped onto cortical and subcortical surfaces, estimated using nn = 10. The first component was excluded due to lack of anatomical specificity (N = 6,624).

#### 2.2 Leading Eigenvector Dynamics Analysis

##### 2.2.1 Correspondence with IFNs

To functionally characterize LEiDA states, we compared them with the seven canonical intrinsic functional networks (IFNs) previously defined by Yeo et al. (Figure 13).

For comparability, our analyses were restricted to the first 100 cortical ROIs of the Schaefer100 parcellation, as IFNs are defined only in cortical space.

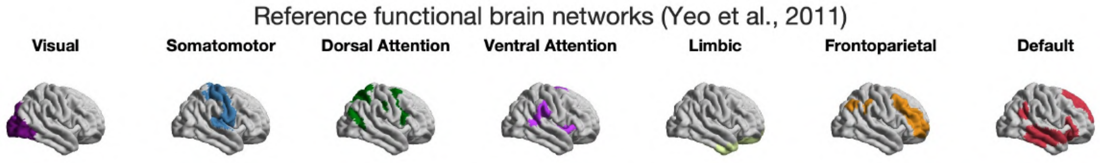

**Figure 13: Reference resting-state networks.** Seven cortical resting-state networks defined by Yeo et al. (2011), namely: Visual, Somatomotor, Dorsal Attention, Ventral Attention, Limbic, Frontoparietal, and Default Mode networks. Each network is represented as a binary mask of the corresponding ROIs from the Schaefer100 parcellation, and color-coded according to its network assignment.

The following Figure 14 expands upon the findings previously presented, adding p-values for all centroids across clustering solutions from $K = 2$ to 20.

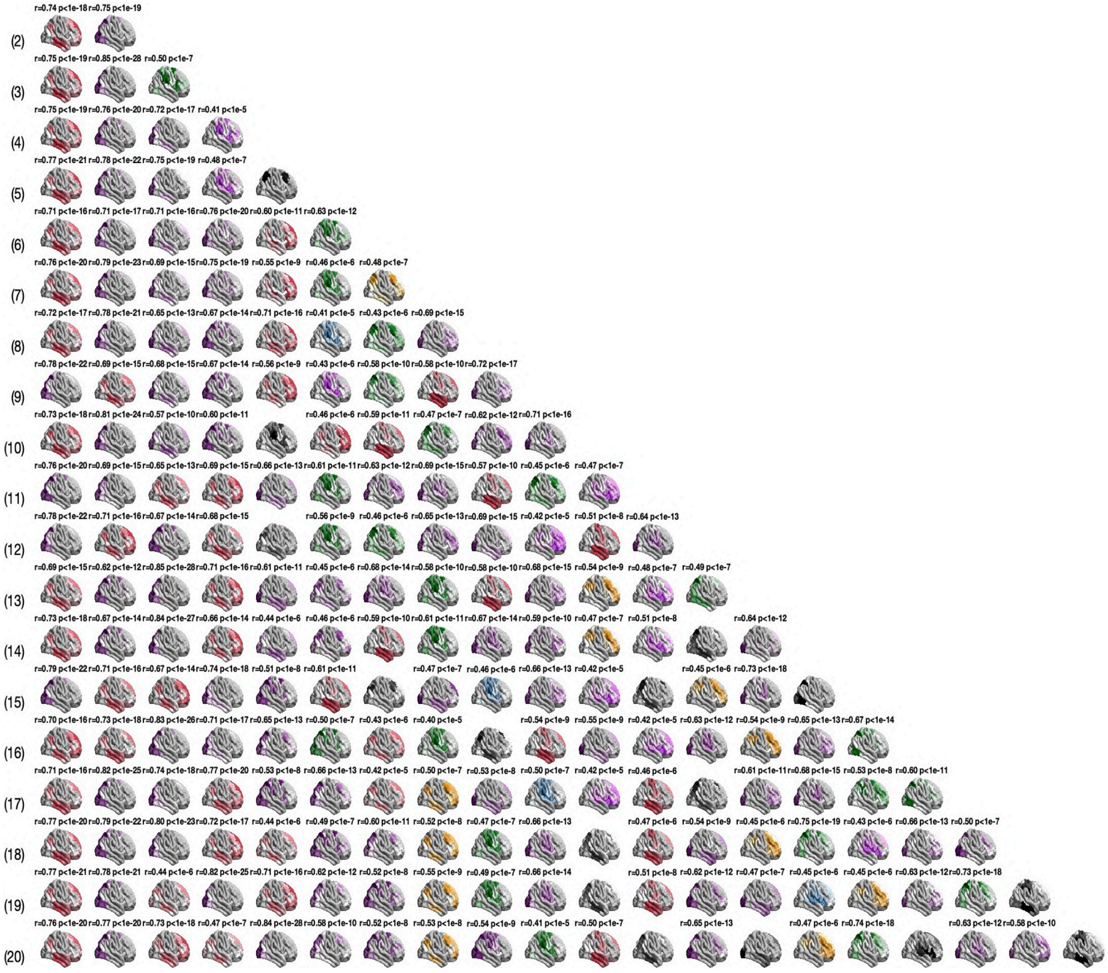

**Figure 14: LEiDA centroids and reference resting-state networks.** Correlations between LEiDA-derived centroids and the seven cortical resting-state networks defined by Yeo et al. (2011). For each K, centroids are ordered according to their FO from the most to the least frequent, and color-coded by the most strongly correlated RSN. Unassigned centroids (r < 0.4) are shown in black. Here both r and p-value are visualized (N = 6,624).

#### 2.3 PLSR analysis

To evaluate whether FCHs can predict patterns of dynamic phase-locking (LEiDA) states, we applied PLSR using the full set of harmonics to model LEiDA states with $K = 6$. Models were tested with both the original ordered harmonics and randomly permuted harmonics, and model performance was assessed using cross-validated MSE.

While the main text presents illustrative results for LEiDA state 1, additional results for states 2 through 6 ($K = 6$) are provided in this Supplementary.

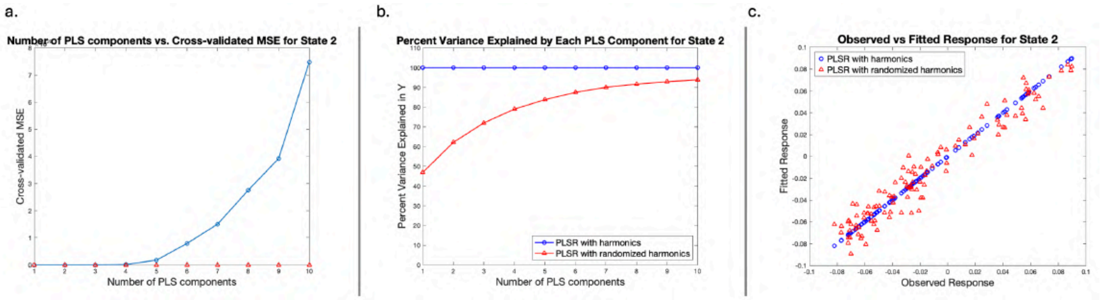

**Figure 15: Prediction of LEiDA states from FCH using partial least square (PLS) regression - state 2.** Partial least squares regression (PLSR) models predicting LEiDA state 2 (Visual network; K = 6) from functional connectome harmonic (FCH) components. a, Mean squared error (MSE) across PLS components. b, Variance explained for ordered and permuted harmonics. c, Observed versus predicted values for the selected model (N = 6,624).

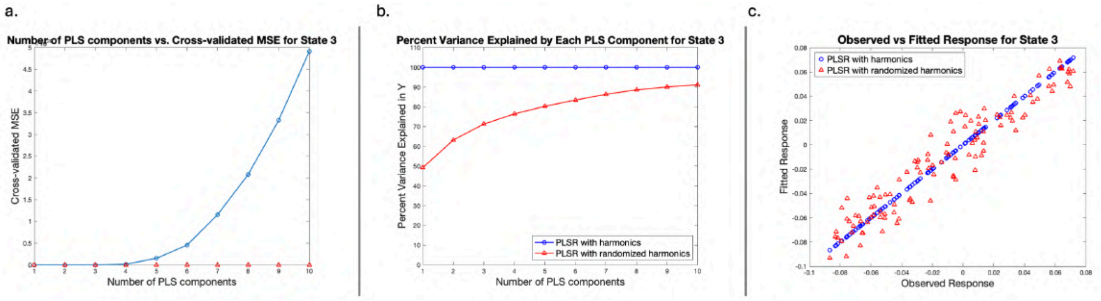

**Figure 16: Prediction of LEiDA states from FCH using PLS regression - state 3.** Partial least squares regression (PLSR) models predicting LEiDA state 3 (Visual network; K = 6) from functional connectome harmonic (FCH) components. a, Mean squared error (MSE) across PLS components. b, Variance explained for ordered and permuted harmonics. c, Observed versus predicted values for the selected model (N = 6,624).

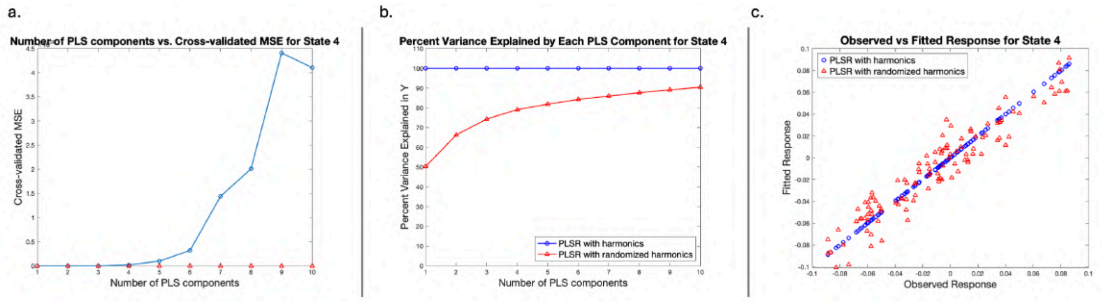

**Figure 17: Prediction of LEiDA states from FCH using PLS regression - state 4.** Partial least squares regression (PLSR) models predicting LEiDA state 4 (Visual network; K = 6) from functional connectome harmonic (FCH) components. a, Mean squared error (MSE) across PLS components. b, Variance explained for ordered and permuted harmonics. c, Observed versus predicted values for the selected model (N = 6,624).

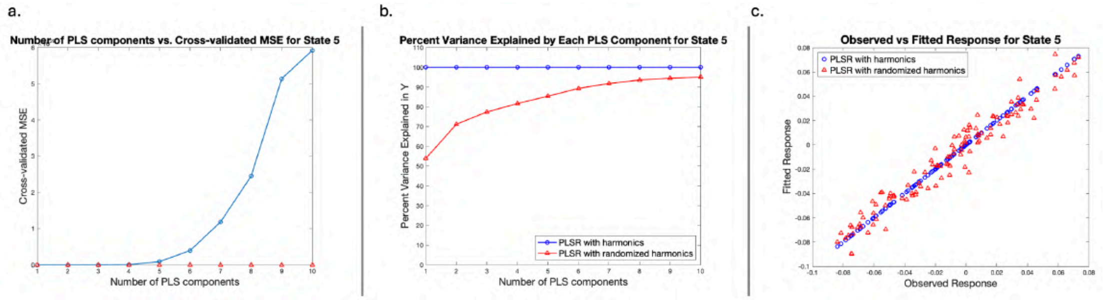

**Figure 18: Prediction of LEiDA states from FCH using PLS regression - state 5.** Partial least squares regression (PLSR) models predicting LEiDA state 5 (Default Mode network; K = 6) from functional connectome harmonic (FCH) components. a, Mean squared error (MSE) across PLS components. b, Variance explained for ordered and permuted harmonics. c, Observed versus predicted values for the selected model (N = 6,624).

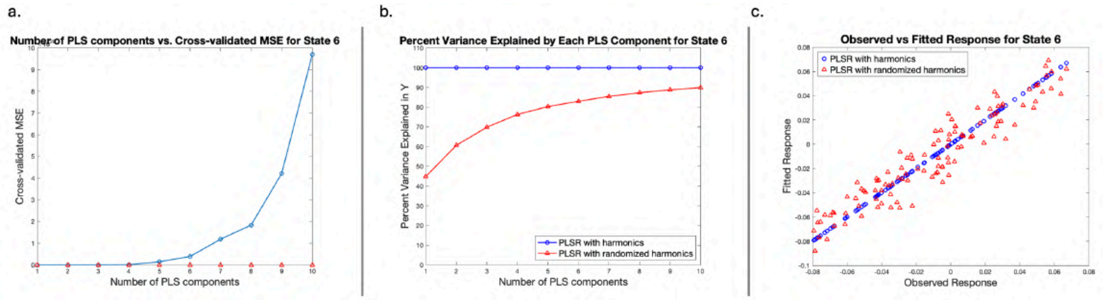

**Figure 19: Prediction of LEiDA states from FCH using PLS regression - state 6.** Partial least squares regression (PLSR) models predicting LEiDA state 6 (Dorsal Attention network; K = 6) from functional connectome harmonic (FCH) components. a, Mean squared error (MSE) across PLS components. b, Variance explained for ordered and permuted harmonics. c, Observed versus predicted values for the selected model (N = 6,624).

#### 2.4 Assessment of site-wise effect

Given the presence of 21 acquisition sites differing by scanner and location, a crucial step in our analysis was dataset harmonization using neuroCombat to control for site effects. This procedure was applied separately to three groups of fMRI-derived features: (i) rs-fMRI Gordon network and subcortical correlations, (ii) FCH-derived power and energy, and (iii) LEiDA-derived fractional occupancy, dwell time, and Markov Chain transition probabilities.

Harmonization effectively removed site-related variability while preserving meaningful inter-individual differences, enabling more reliable and generalizable analyses.

##### 2.4.1 FCH-derived metrics

We first examined the effect of harmonization on the relationship between FCH-derived power and energy and each of the twenty-one sites.

Partial correlations were computed between the first twenty FCH-derived metrics and each site, while regressing out the contributions of the remaining twenty sites, for both the original and harmonized datasets. This approach ensured that the unique contribution of each site was isolated and evaluated.

Before harmonization, weak associations between FCH metrics and site were observed for some harmonics, with correlation coefficients ranging from $r = 0.01$ to $r = 0.05$ in both positive and negative directions (Figure 20).

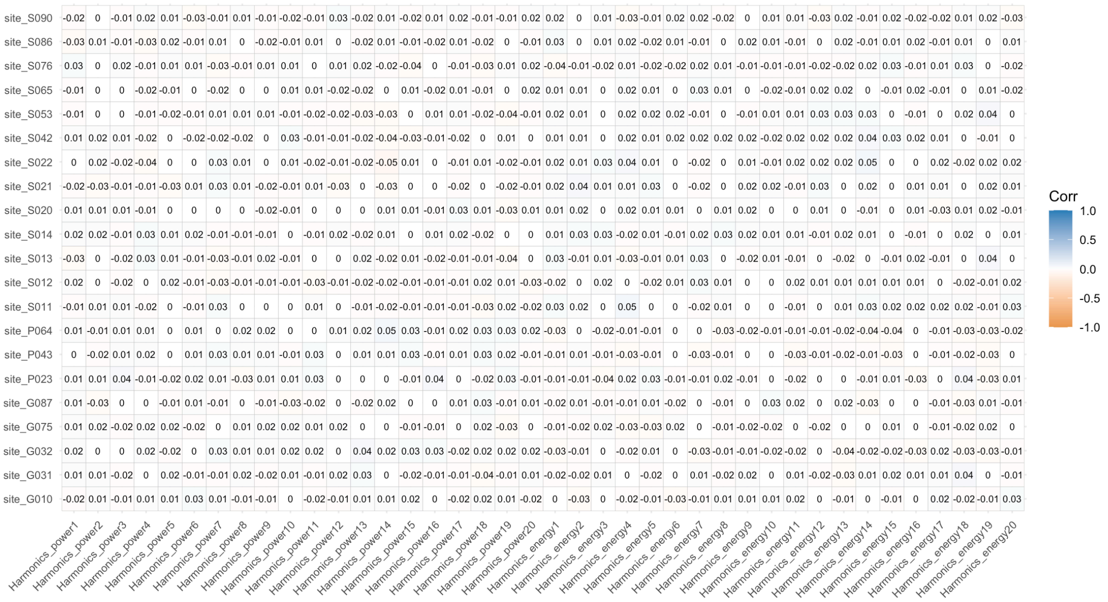

**Figure 20: Correlations between FCH metrics and site before harmonization.** Correlation matrix showing site-wise associations with FCH metrics before the harmonization procedure. FCH-derived metrics are on x-axis while sites on y-axis (N = 4,453).

After harmonization, these correlations were substantially reduced for both energy and power across all 21 sites. As shown in Figure 21, almost all FCH-derived features showed null correlations with site, and none exceeded $|r| = 0.02$, representing a below-average effect in the ABCD dataset.

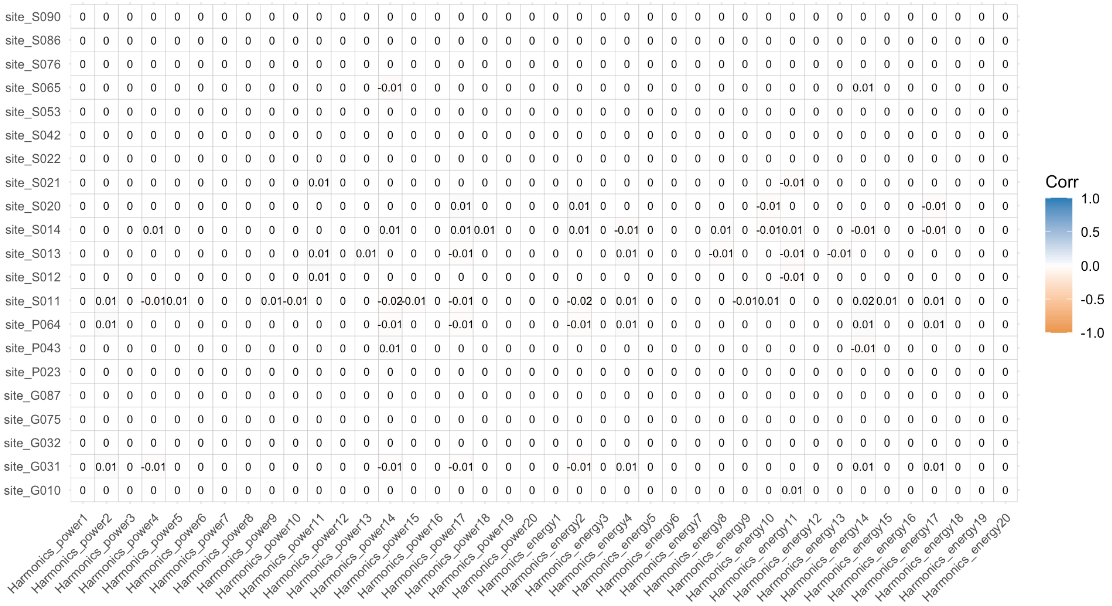

**Figure 21: Correlations between FCH metrics and site after harmonization.** Correlation matrix showing site-wise associations with FCH metrics after the harmonization procedure. FCH-derived metrics are on x-axis while sites on y-axis (N = 4,453).

##### 2.4.2 LEiDA-derived metrics

To assess whether similar patterns existed for LEiDA metrics, the same partial correlation analysis was performed between all 21 sites and both FO and DT, focusing on LEiDA states significantly associated with intrinsic functional networks (IFNs) as defined by Yeo et al. For this exploratory analysis, we set $K = 15$, as this solution provided the highest number of IFNs matched while keeping the dimensionality low.

Figure 22 shows that no significant associations were observed between LEiDA FO or DT and site, even before harmonization, with all correlations ranging between $r = 0$ and $r = 0.05$ in both directions. Whereas, following harmonization, correlations between LEiDA metrics and site were further reduced for both FO and DT across all sites.

As with FCH-derived metrics, almost all LEiDA-derived features showed null correlations, and none exceeded $|r| = 0.01$, confirming that harmonization effectively mitigated site-related effects.

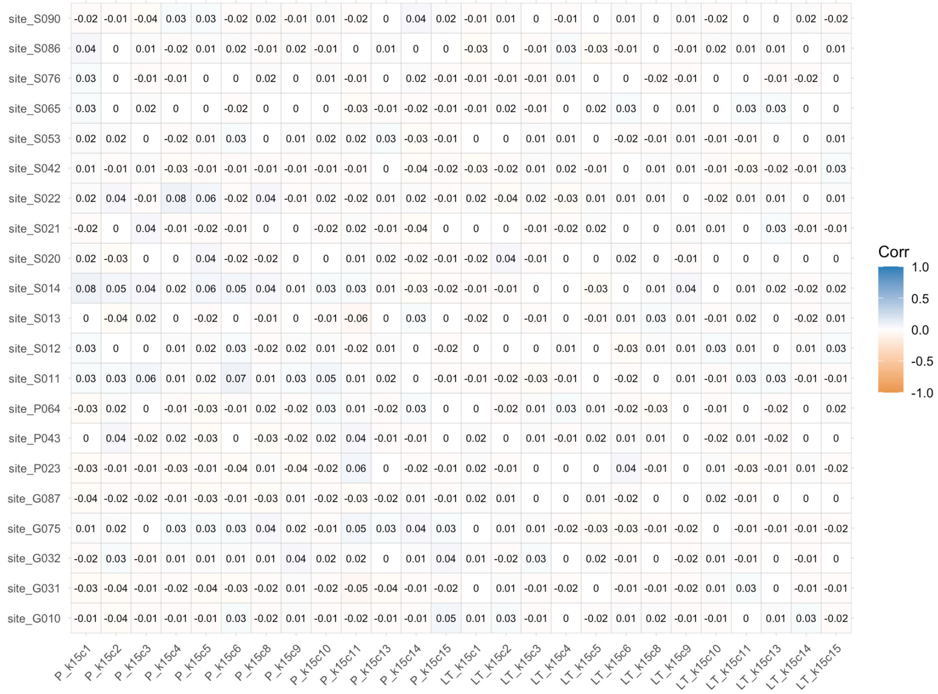

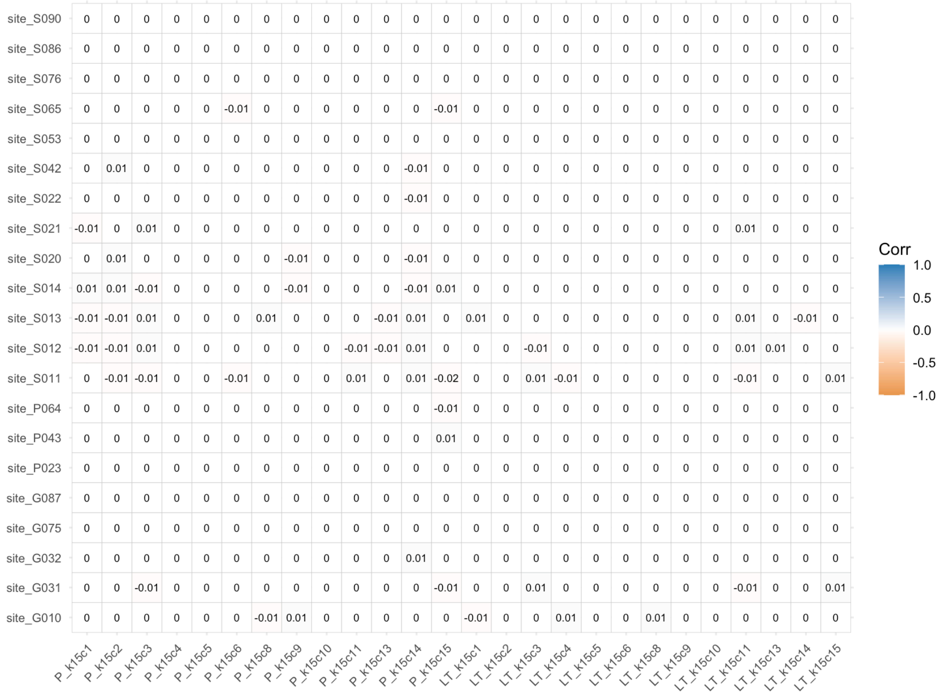

**Figure 22: Correlations between LEiDA FO and DT, and sites for K=15.** Correlation matrix displaying the association between DT of each LEiDA state (y-axis) and the 21 acquisition sites (x-axis) in the ABCD dataset before (upper figure) and after (lower figure) harmonization (N = 4,453). FO; fractional occupancy. DT; dwell time.

#### 2.5 Partial correlation analyses

##### 2.5.1 Partial correlation analysis on FCH metrics

We computed partial correlations between FCH-derived power and energy for the first twenty harmonics, excluding the one associated with the null eigenvalue, and demographic variables (site, sex, ethnicity, interview age, PDS, and TMI). Results are reported for the original FCH dataset (Figure 23).

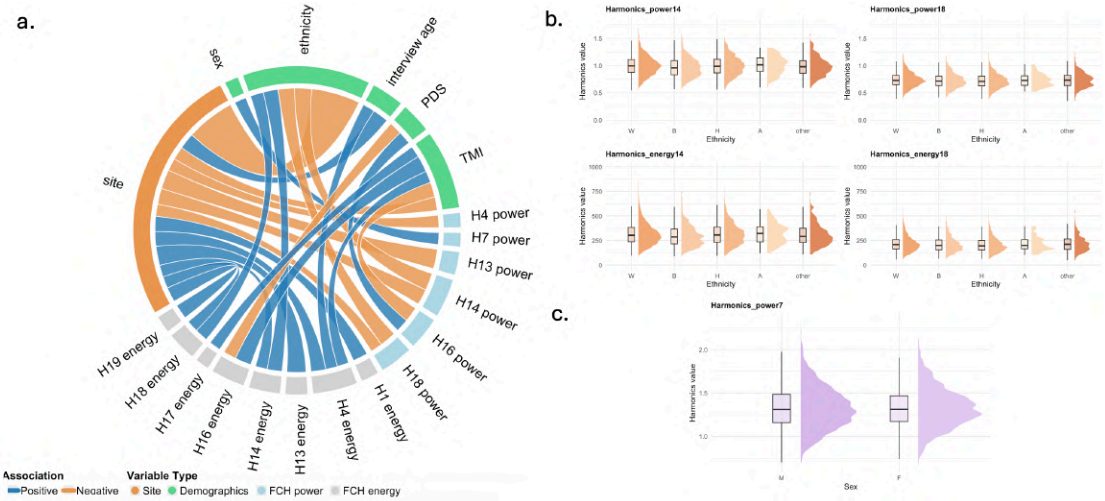

**Figure 23: Correlations between FCH metrics and demographic and developmental variables - original dataset. a,** Circos plots showing significant partial correlations ($|r| > 0.03$) between FCH energy (grey) and power (light blue) with key variables for the original dataset. **b,** Half-violin plot showing the distribution of FCH-derived metrics significantly associated with ethnicity. The x-axis represents ethnicity groups: W = White, B = Black, H = Hispanic, A = Asian, other (i.e., any other different ethnicity from those analysed); the y-axis indicates the metric value. **c,** Half-violin plot showing the distribution of FCH-derived metrics significantly associated with sex. The x-axis represents sex groups: M = males, F = females; the y-axis indicates the metric value (N = 4,453). PDS = Pubertal Development Scale; TMI = Triponderal Mass Index.

Results are now reported for both the original and the harmonized FCH datasets (Figure 24). Importantly, after harmonization, only correlations involving site were affected, while all other relationships remained stable. Before harmonization, site showed significant associations with all other variables, including both demographics and connectome-derived measures.

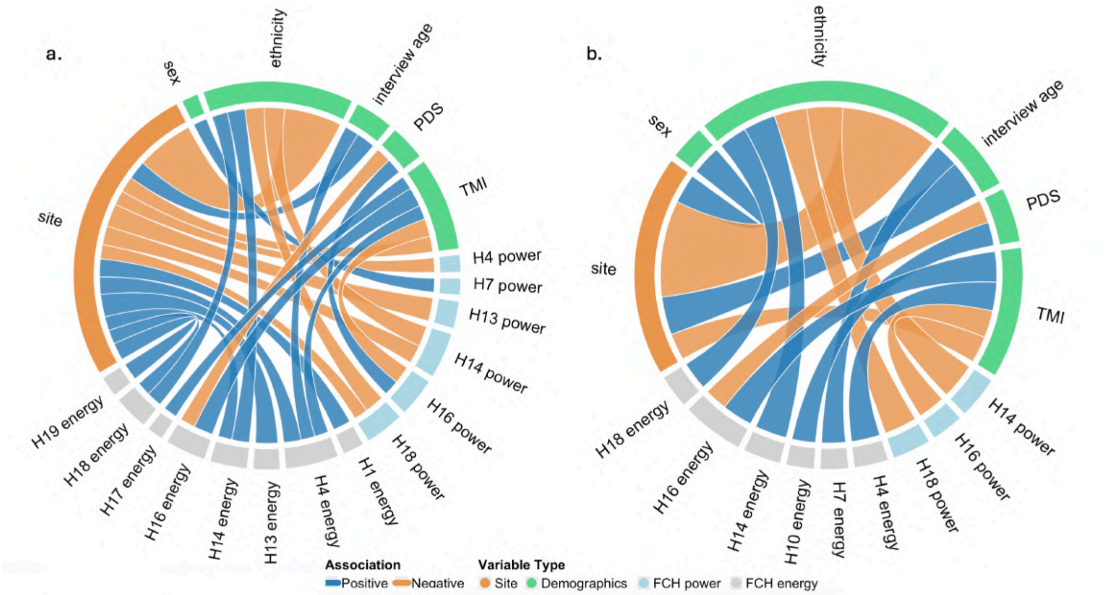

**Figure 24: Correlations between FCH metrics and covariates for both datasets. a,** Circos plots showing significant partial correlations (|r| > 0.03) between FCH energy/power and demographics, for the original dataset. **b,** Circos plots showing significant partial correlations (|r| > 0.03) between FCH energy/power and demographics, for the harmonized dataset (N = 4,453). PDS = Pubertal Development Scale; TMI = Triponderal Mass Index.

Effect sizes were generally small ($r > 0.03$) but consistent with a small-to-medium effect ($0.03 < r < 0.05$) in the context of the ABCD Study. These results indicate that demographic variables show selective associations with specific FCH harmonics.

##### 2.5.2 Partial correlation analysis on LEiDA metrics

We performed the same analysis on all LEiDA-derived metrics (FO, DT, and Markov Chain transition probabilities) for all states that were successfully assigned to one of the reference resting-state networks, across $K = 2 - 20$. Results are reported for the original LEiDA dataset (Figure 25).

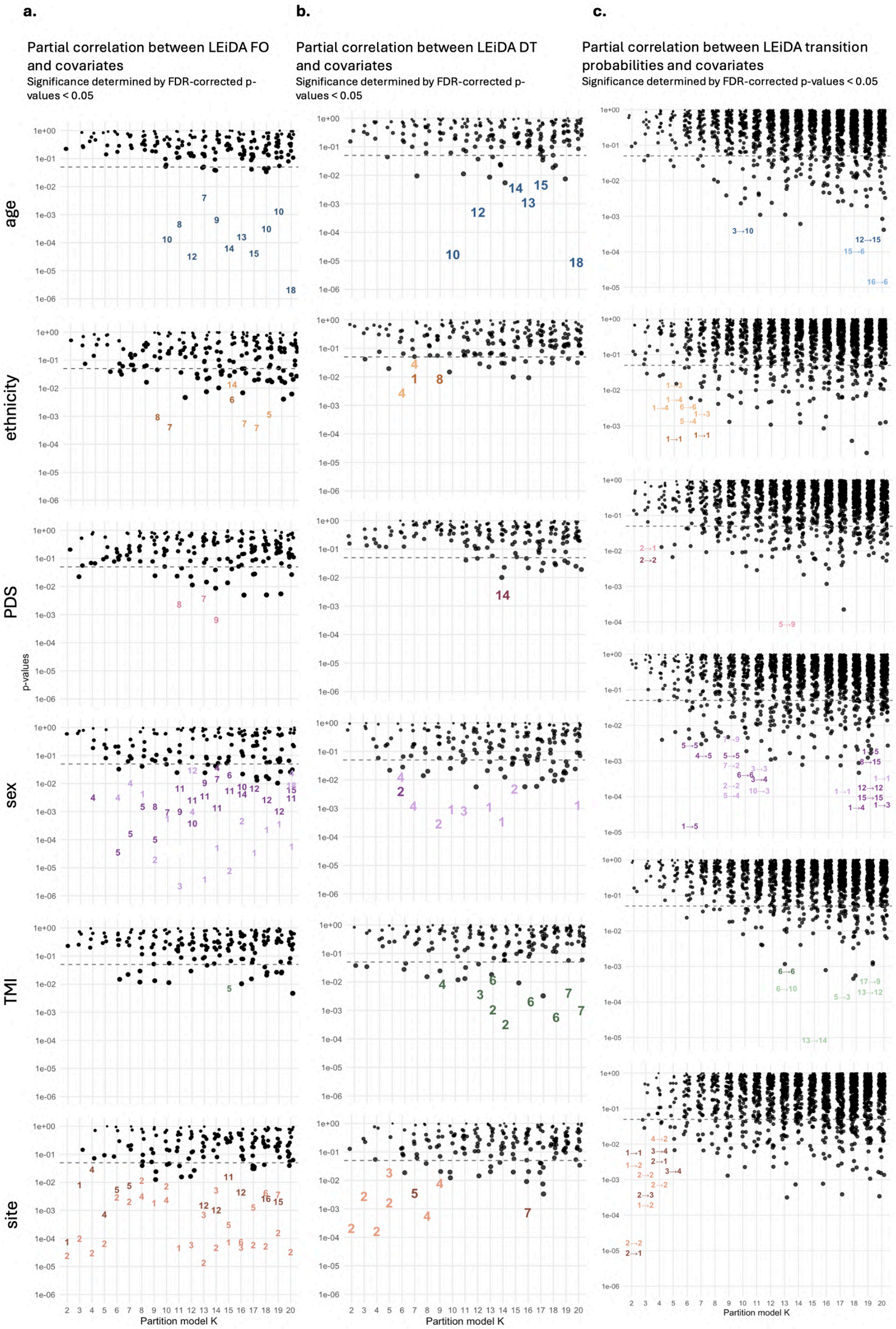

**Figure 25: Partial correlations between LEiDA metrics and demographics - original dataset.** Correlation coefficients and significance (color scale) are reported for LEiDA states a) probabilities, b) lifetimes and c) transitions significantly associated with demographic variables in the harmonized dataset (N = 4453). Rows represent LEiDA states, columns the K partitions. Positive associations are represented with lighter colors, while negative associations are represented with darker colors. PDS = Pubertal Development Scale; TMI = Triponderal Mass Index.

##### 2.5.3 Additional results from partial correlations on LEiDA metrics

For completeness, tables preceding the plots report all significant associations between LEiDA metrics and covariates across the 19 clustering solutions ($K = 2 - 20$). Separate tables are provided for Fractional Occupancy (FO), Dwell Time (DT), and transition probabilities, in both the original and harmonized datasets. Each table reports the state, clustering solution, covariate, and direction of effect. These detailed tabular summaries complement the subsequent graphical representations.

To visualize the distribution of data corresponding to significant associations, half-violin plots were used for categorical covariates (ethnicity, sex, and site), while scatter plots were used for continuous covariates (TMI, interview age, and PDS).

Half-violin plots illustrate the distribution of LEiDA metrics across categories, whereas scatter plots show the relationship with continuous covariates, displaying the distribution of a continuous variable within each group of a categorical factor. The width of the violin at any given value represents the density of data points, so wider sections indicate more frequent values. In these plots, the median is overlaid, allowing a clear comparison of central tendency and variability between groups, while preserving a detailed view of the overall distribution shape.

Additionally, associations between LEiDA metrics and continuous variables are summarized in heatmaps, where each cell represents a significant association. Color intensity indicates the strength of the association (darker colors correspond to smaller $p$-values), and the correlation value is displayed inside the cell. This provides an intuitive understanding of both the magnitude and direction (positive or negative) of each significant association.

###### 2.5.3.1 Original dataset

###### 2.5.3.1.1 Fractional occupancy (FO)

**Table 3: Detailed associations between Fractional Occupancy, states, and demographic variables - original dataset.** Table providing a comprehensive overview of all significant associations between FO and demographic variables across all states and K values. Specific states, the corresponding K values, and the direction/range of correlations are included.

| Variable | Network(s) affected | States (K) | Effect (Pearson's r) |
|---|---|---|---|
| **Age** | Visual Network | 10 (K=10,18,19); 8 (K=11); 12 (K=12); 7 (K=13); 9 (K=14); 14 (K=15); 13 (K=16); 15 (K=17); 18 (K=20) | Negative (-0.05 to -0.07) |
| **Ethnicity** | DMN | 8 (K=9); 7 (K=10); 6 (K=15); 7 (K=16,17); 5 (K=18) | - |
| | Visual Network | 14 (K=15) | - |
| **PDS** | Visual Network | 8 (K=11); 7 (K=13); 9 (K=14) | Positive (0.04 to 0.05) |
| **Sex** | DMN | 1 (K=8,10,13,14,17-20); 2 (K=9,15,16); 3 (K=11); 4 (K=12,14,20); 5 (K=6–9); 6 (K=10,15); 7 (K=10,14); 8 (K=9); 9 (K=11,13); 10 (K=16); 11 (K=12,20); 12 (K=17–19) | - |
| | Visual | 4 (K=6,7); 12 (K=12); 18 (K=20) | - |
| | Ventral Attention | 4 (K=4); 10 (K=12); 11 (K=11,15) | - |
| | Frontoparietal | 11 (K=13,14); 14 (K=16); 15 (K=20) | - |
| **Site** | DMN | 1 (K=2,3); 5 (K=6,7) | - |
| | Visual | 1 (K=9,11,15); 2 (K=2–8,10,13,14,17–20); 3 (K=12–14,16); 4 (K=8,10); 5 (K=15,17); 6 (K=18); 7 (K=19) | - |
| | Ventral Attention | 4 (K=4,5); 11 (K=15); 12 (K=13,14,16); 16 (K=18) | - |
| | Dorsal Attention | 6 (K=16) | - |
| | Somatomotor | 15 (K=19) | - |
| **TMI** | Visual Network | 5 (K=15) | Negative (-0.04) |

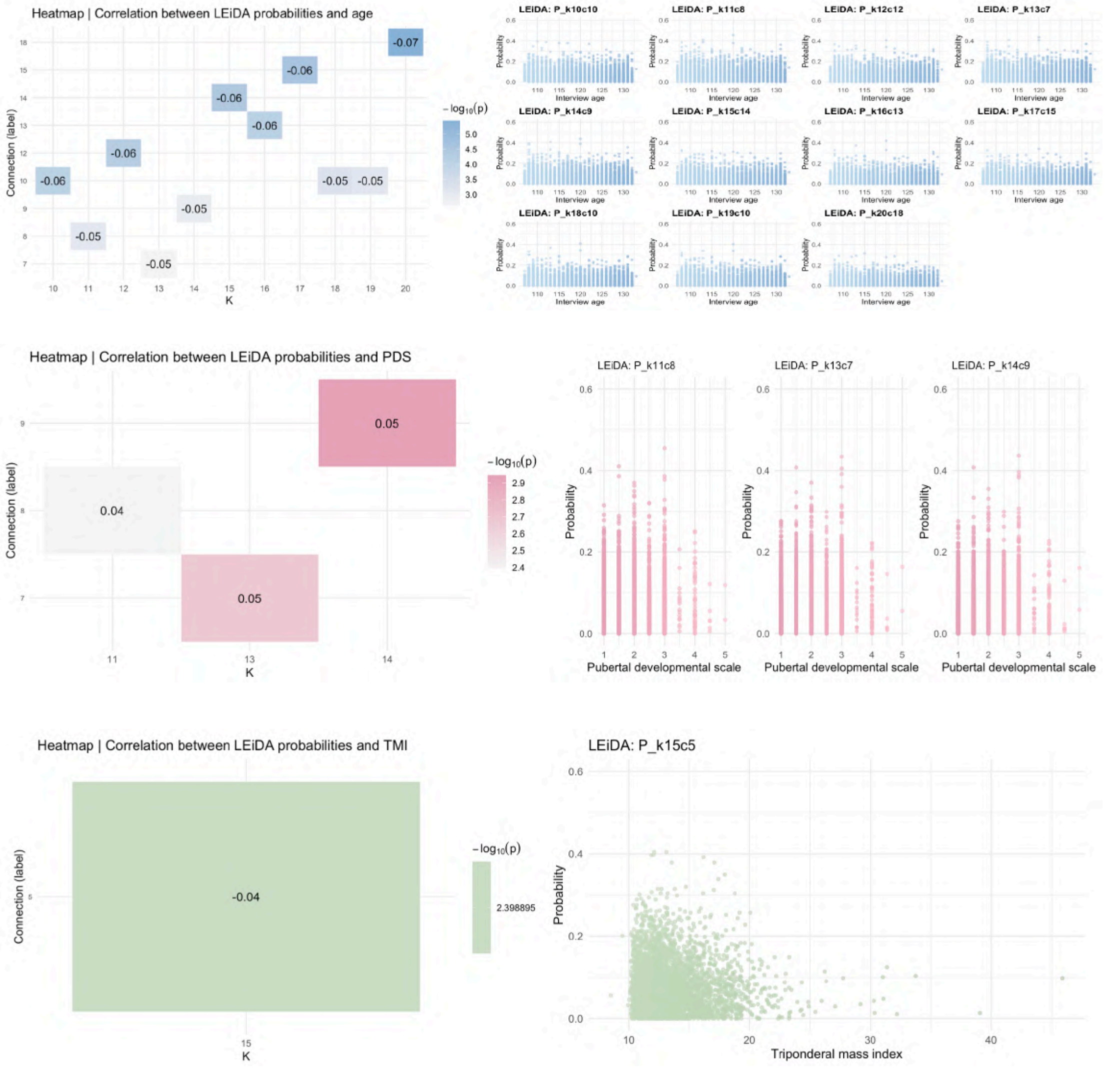

**Figure 26: Heatmaps and scatter plots between LEiDA FO and continuous variables - original dataset.** Heatmap showing correlation values and significance (color scale) for LEiDA state probabilities significantly associated with continuous variables (PDS, interview age and TMI) in the original dataset. Rows represent states, columns the K partitions. Under, scatter plots showing the distribution of LEiDA-derived state probabilities significantly associated with continuous variables (FDR-corrected $p < 0.05$). The x-axis represents covariates values. The y-axis indicates the probability of occurrence for each LEiDA state (N = 4,453). PDS = Pubertal Development Scale; TMI = Triponderal Mass Index.

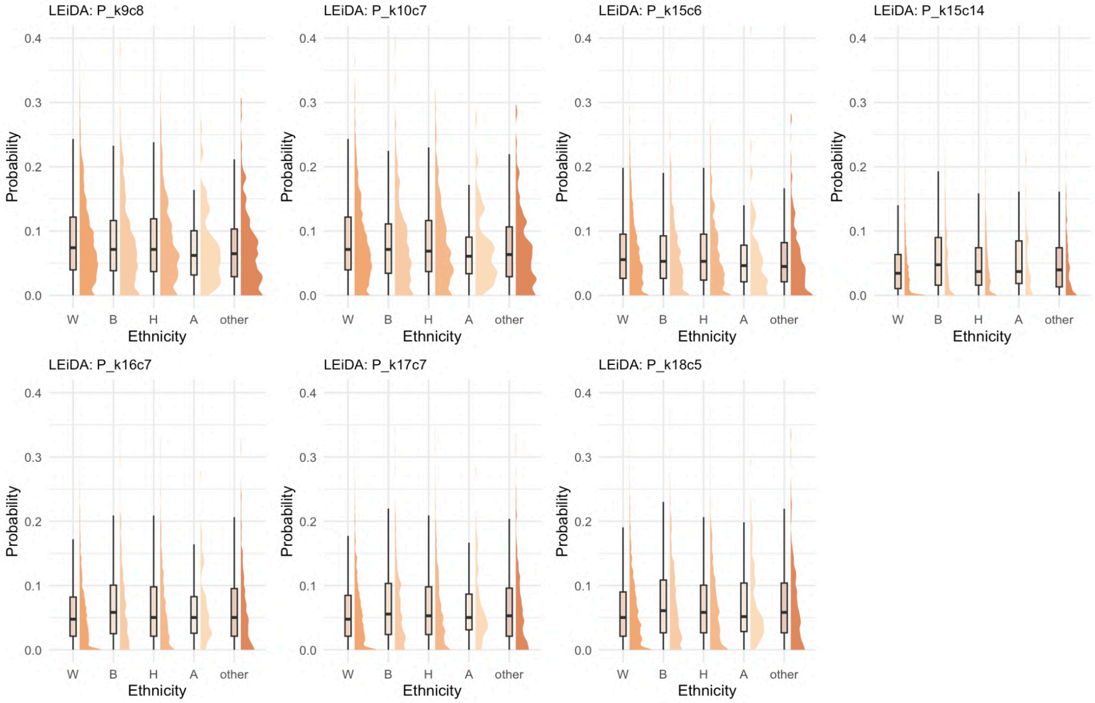

**Figure 27: Distribution of LEiDA state FO across ethnicity groups - original dataset.** Half-violin plots showing the distribution of LEiDA-derived state FO significantly associated with ethnicity (FDR-corrected $p < 0.05$). The x-axis represents ethnicity groups: W = White, B = Black, H = Hispanic, A = Asian, other. The y-axis indicates the probability of occurrence for each LEiDA state. Only states with significant associations are displayed (N = 4,453).

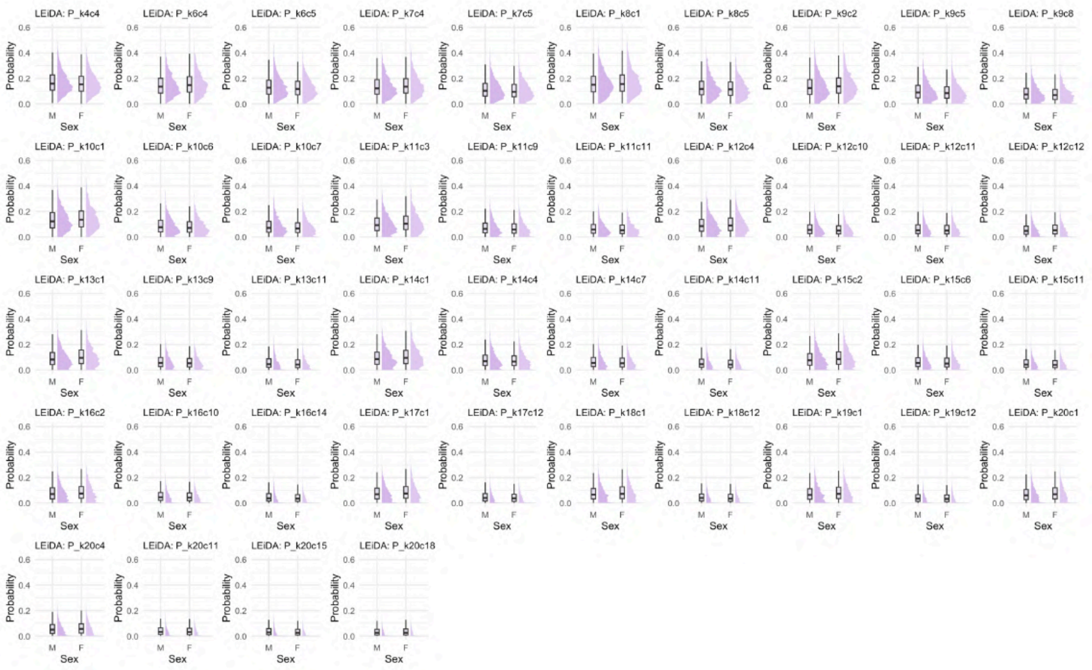

**Figure 28: Distribution of LEiDA state FO across sex groups - original dataset.** Half-violin plots showing the distribution of LEiDA-derived state FO significantly associated with sex (FDR-corrected p < 0.05). The x-axis represents sex groups: M = males, F = females; the y-axis the probability of occurrence for each LEiDA state. Only states with significant associations are displayed (N = 4,453).

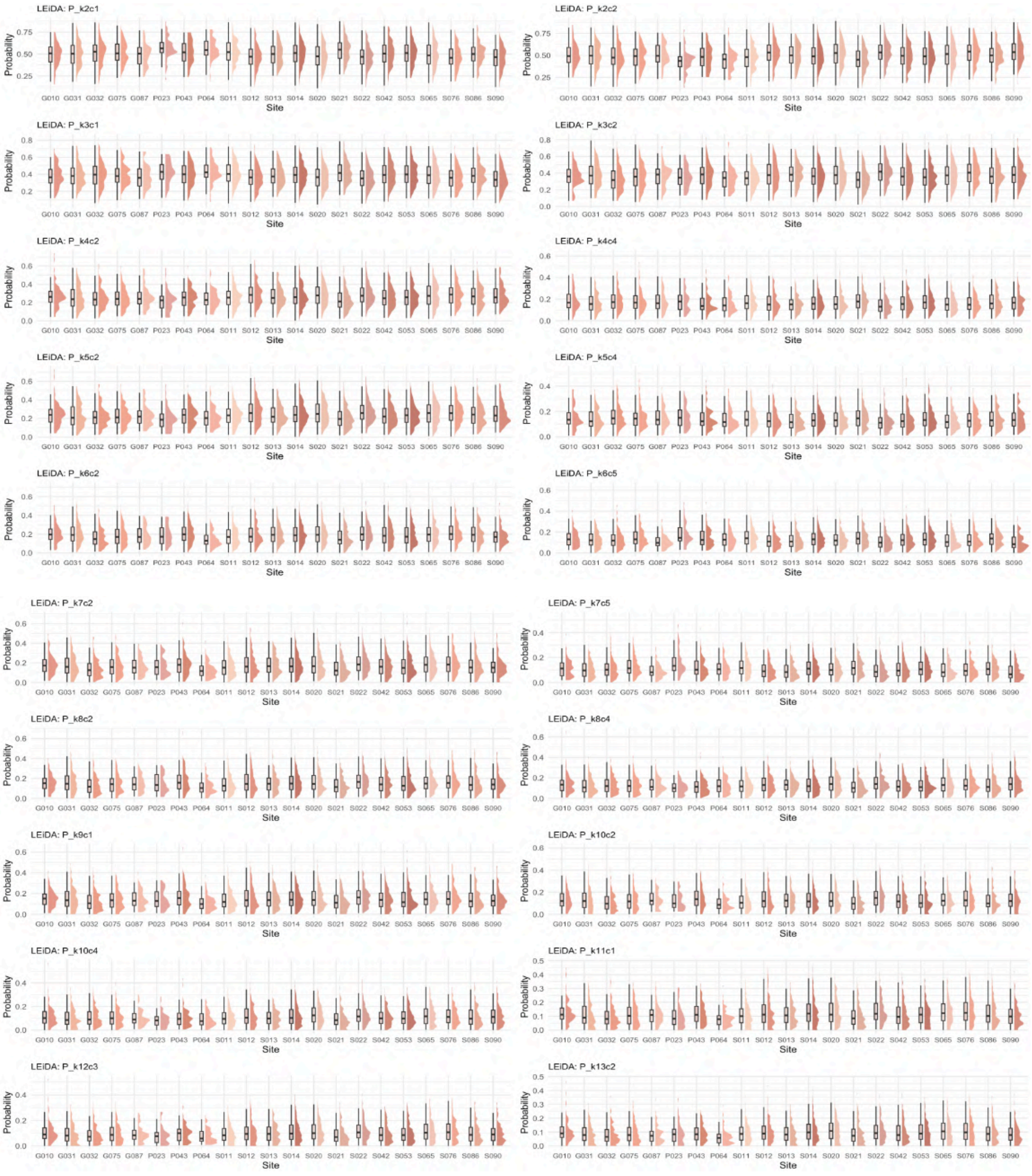
LEiDA: P_k2c1
LEiDA: P_k2c2
LEiDA: P_k3c1
LEiDA: P_k3c2
LEiDA: P_k4c2
LEiDA: P_k4c4
LEiDA: P_k5c2
LEiDA: P_k5c4
LEiDA: P_k6c2
LEiDA: P_k6c5
LEiDA: P_k7c2
LEiDA: P_k7c5
LEiDA: P_k8c2
LEiDA: P_k8c4
LEiDA: P_k9c1
LEiDA: P_k10c2
LEiDA: P_k10c4
LEiDA: P_k11c1
LEiDA: P_k12c3
LEiDA: P_k13c2
Probability
Site
G010 G031 G032 G075 G087 P023 P043 P064 S011 S012 S013 S014 S020 S021 S022 S042 S053 S065 S076 S086 S090

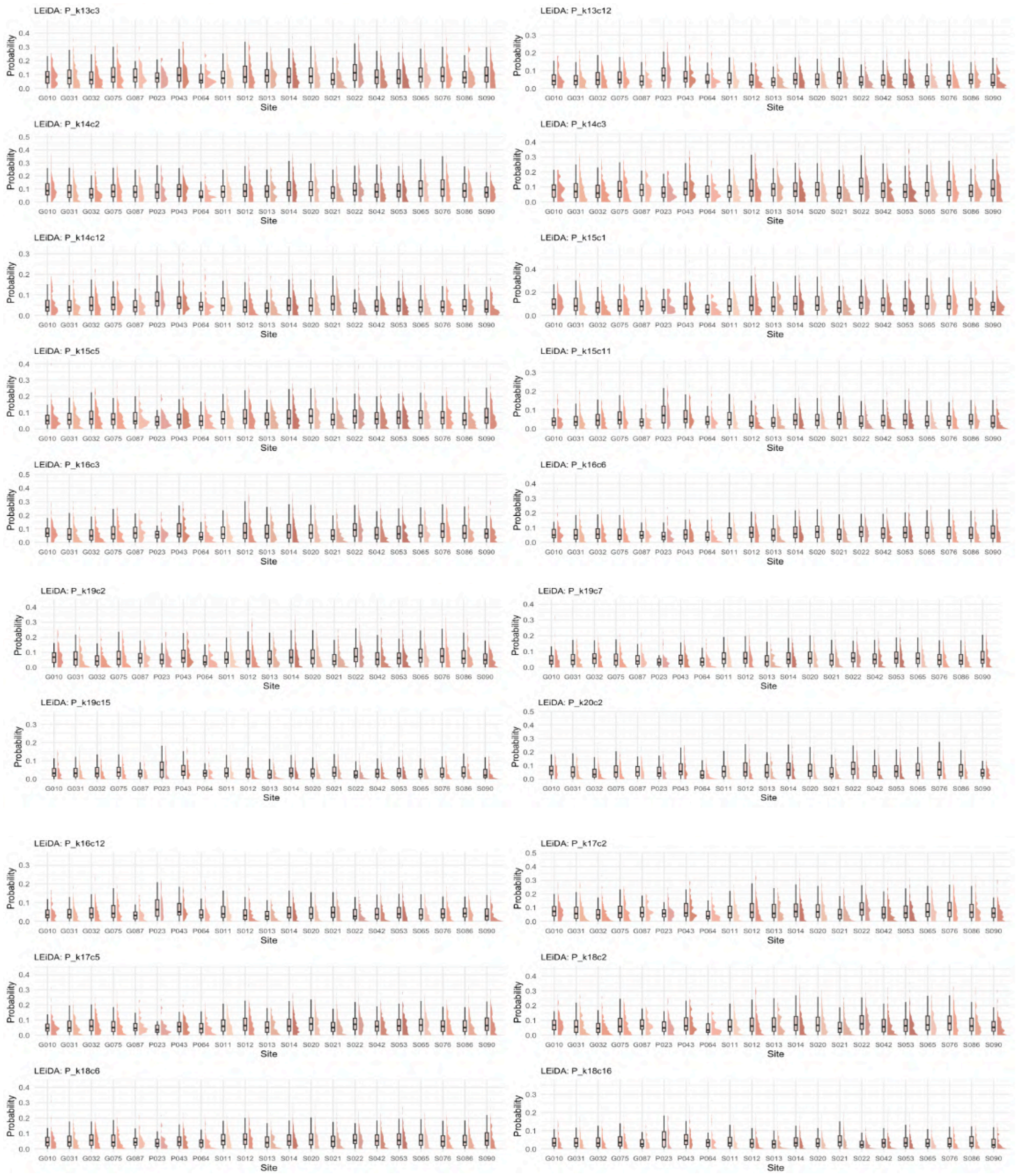

**Figure 29: Distribution of LEiDA state FO across sites - original dataset.** Half-violin plots showing the distribution of LEiDA-derived state FO significantly associated with site (FDR-corrected p $< 0.05$). The x-axis represents different sites. The y-axis indicates the FO for each LEiDA state. Only states with significant associations are displayed (N = 4,453).

##### 2.5.3.1.2 Dwell Time (DT)

**Table 4: Detailed associations between Dwell Time, states, and demographic variables - original dataset.** Table providing a comprehensive overview of all significant associations between DT and demographic variables across all states and K values. Specific states, the corresponding K values, and the direction/range of correlations are included.

| Variable | Network(s) affected | States (K) | Effect (Pearson's r) |
|---|---|---|---|
| **Age** | Visual Network | 10 (K=10); 12 (K=12); 13 (K=16); 14 (K=15); 15 (K=17); 18 (K=20) | Negative (-0.05 to -0.06) |
| **Ethnicity** | DMN | 1 (K=7); 8 (K=9) | - |
| | Visual Network | 4 (K=6,7) | - |
| **PDS** | Visual Network | 14 (K=14) | Negative (-0.05) |
| **Sex** | DMN | 1 (K=10,13,14,20); 2 (K=9,15); 3 (K=11) | - |
| | Visual Network | 2 (K=6); 4 (K=6,7) | - |
| **Site** | DMN | 5 (K=7); 7 (K=16) | - |
| | Visual Network | 2 (K=2–5); 3 (K=5); 4 (K=8,9) | - |
| **TMI** | Visual | 2 (K=13,14); 3 (K=12); 4 (K=9); 6 (K=13,18); 7 (K=19,20) | Negative (-0.04 to -0.05) |
| **TMI** | Dorsal Attention | 6 (K=16) | Negative (-0.05) |

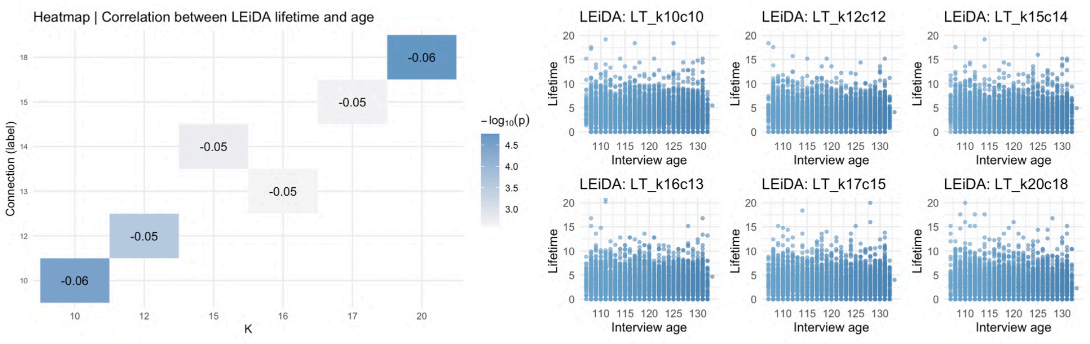

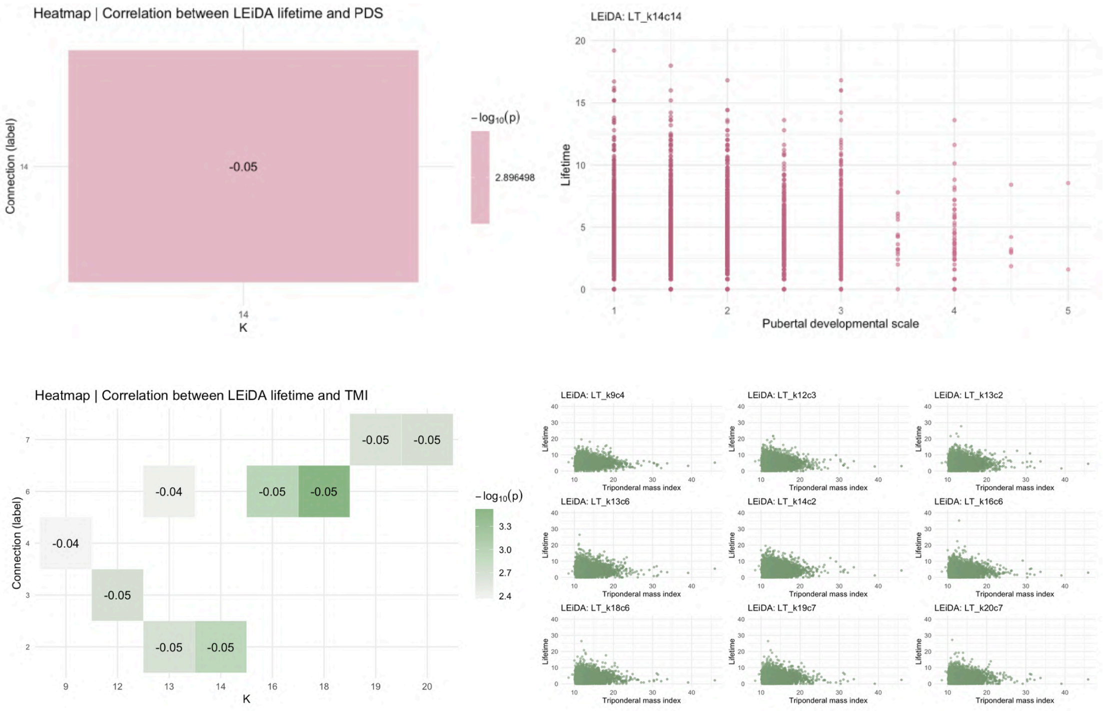

**Figure 30: Heatmaps and scatter plots between LEiDA DT and continuous variables - original dataset.** Heatmap showing correlation values and significance (color scale) for LEiDA state lifetimes significantly associated with continuous variables (PDS, interview age and TMI) in the original dataset. Rows represent states, columns the K partitions. Under, scatter plots showing the distribution of LEiDA-derived state probabilities significantly associated with continuous variables (FDR-corrected $p < 0.05$). The x-axis represents covariates values. The y-axis indicates the probability of occurrence for each LEiDA state (N = 4,453). PDS = Pubertal Development Scale; TMI = Triponderal Mass Index.

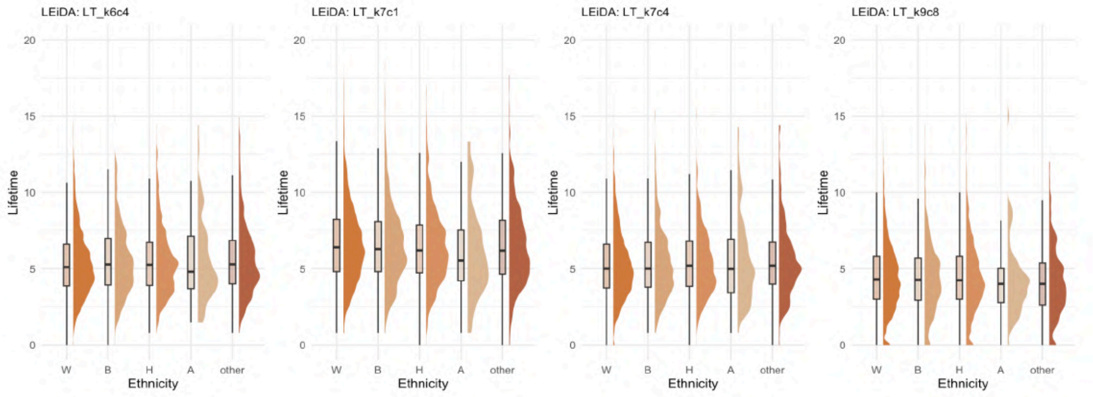

**Figure 31: Distribution of LEiDA state DT across ethnicity groups - original dataset.** Half-violin plots showing the distribution of LEiDA-derived state lifetimes significantly associated with ethnicity (FDR-corrected $p < 0.05$). The x-axis represents ethnicity groups: W = White, B = Black, H = Hispanic, A = Asian, other. The y-axis indicates the DT for each LEiDA state. Only states with significant associations are displayed (N = 4,453).

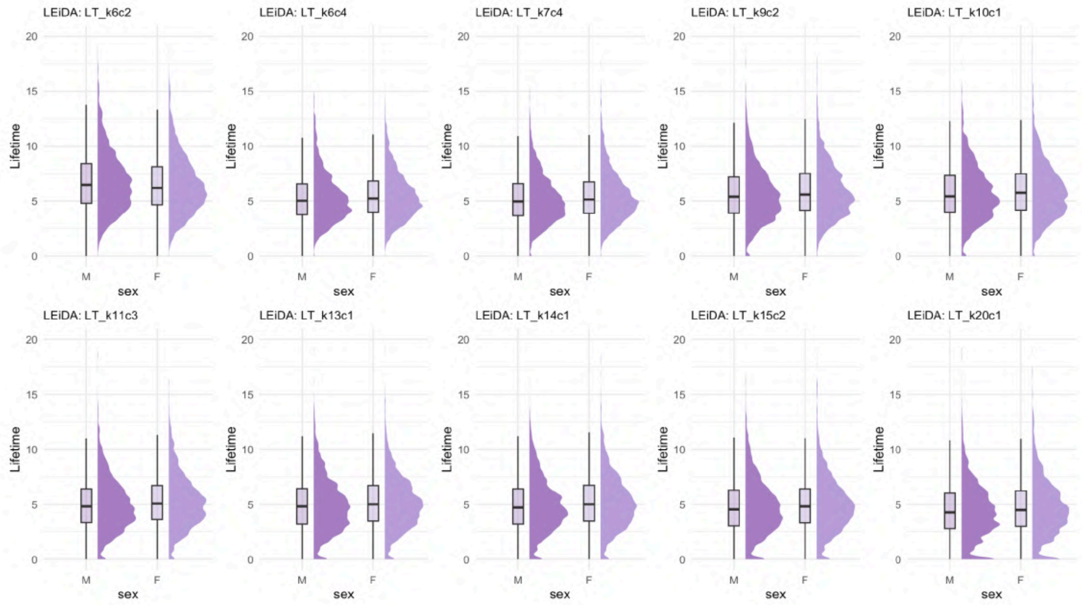

**Figure 32: Distribution of LEiDA state DT across sex groups - original dataset.** Half-violin plots showing the distribution of LEiDA-derived state lifetimes significantly associated with sex (FDR-corrected $p < 0.05$). The x-axis represents sex groups: M = males, F = females. The y-axis indicates the DT for each LEiDA state. Only states with significant associations are displayed (N = 4,453).

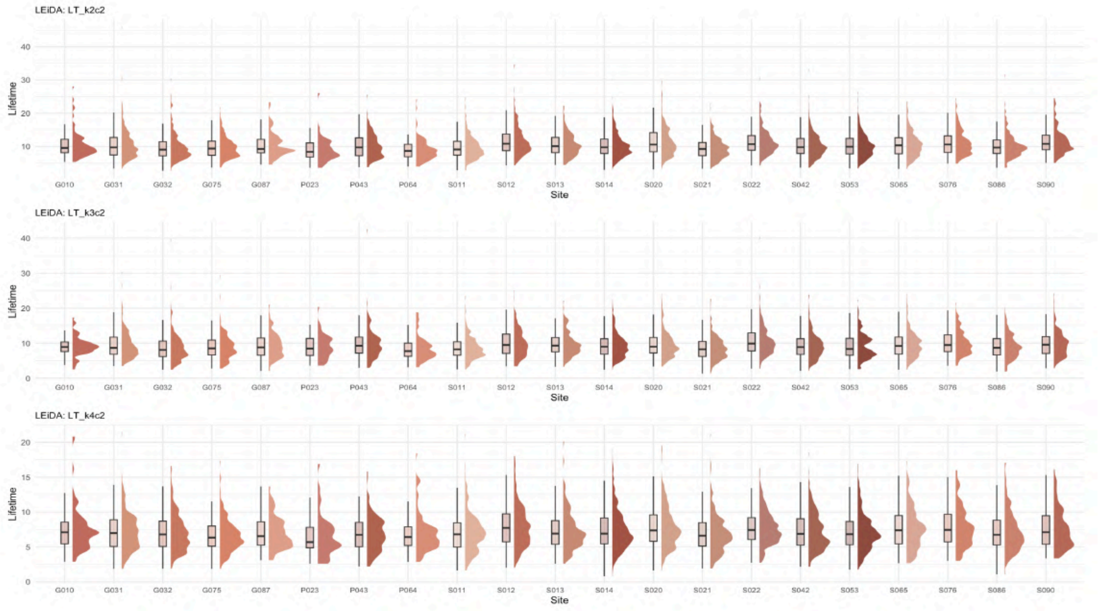

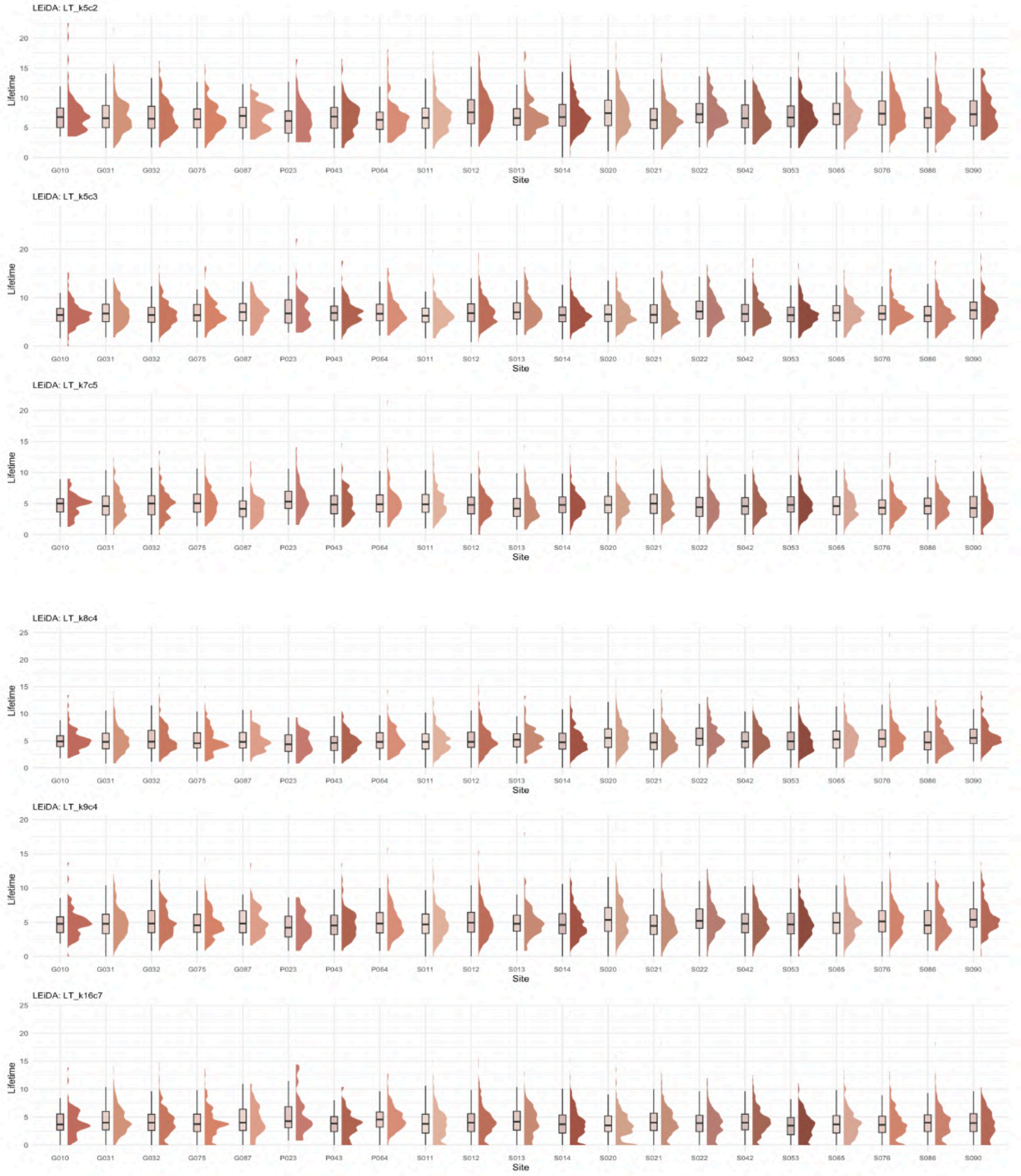

**Figure 33: Distribution of LEiDA state DT across sites - original dataset.** Half-violin plots showing the distribution of LEiDA-derived state lifetimes significantly associated with site (FDR-corrected $p < 0.05$). The x-axis represents different sites. The y-axis indicates DT for each LEiDA state. Only states with significant associations are displayed (N = 4,453).

###### 2.5.3.1.3 Markov Chain transition probabilities

**Table 5: Detailed associations between transitions, states, and demographic variables - original dataset.** Table providing a comprehensive overview of all significant associations between trajectories and demographic variables across all states and K values. Specific states, the corresponding K values, and the direction/range of correlations are included.

| Variable | Trajectories affected | States (K) | Effect (Pearson's r) |
|---|---|---|---|
| **Age** | DMN to SMN | 12 to 15 (K=19) | Negative (-0.06) |
| | VIS to VIS | 3 to 10 (K=10) | Negative (-0.05) |
| | DAN to VIS | 15 to 6 (K=18); 16 to 6 (K=20) | Positive (0.06 to 0.07) |
| **Ethnicity** | DMN to DMN | 1 to 1 (K=5,7) | - |
| | DMN to VIS | 1 to 3 (K=5,7); 5 to 4 (K=6) | - |
| | DMN to VAN | 1 to 4 (K=4,5) | - |
| | DAN to DAN | 6 to 6 (K=6) | - |
| **PDS** | VIS to DMN | 2 to 1 (K=3); 5 to 9 (K=13) | Positive (0.04 to 0.06) |
| | VIS to VIS | 2 to 2 (K=3) | Negative (-0.04) |
| **Sex** | DMN to DMN | 1 to 5 (K=6,19); 5 to 5 (K=6,9); 2 to 2 (K=9); 6 to 6 (K=10); 3 to 3 (K=11); 3 to 4 (K=11); 1 to 1 (K=17,20); 1 to 4 (K=18); 12 to 12 (K=19); 1 to 3 (K=20) | - |
| | DMN to VIS | 5 to 4 (K=9) | - |
| | VIS to DMN | 4 to 5 (K=7) | - |
| | DAN to DMN | 7 to 2 (K=9); 10 to 3 (K=11) | - |
| | DAN to VIS | 7 to 9 (K=9) | - |
| | FPN to SMN | 8 to 15 (K=19) | - |
| | SMN to SMN | 15 to 15 (K=19) | - |
| **Site** | DMN to DMN | 1 to 1 (K=2) | - |
| | DMN to VIS | 1 to 2 (K=2,3) | - |
| | VIS to DMN | 2 to 1 (K=2,4) | - |
| | VIS to VIS | 2 to 2 (K=2-4) | - |
| | VIS to DAN | 2 to 3 (K=3) | - |
| | VIS to VAN | 3 to 4 (K=4,5) | - |
| | VAN to VIS | 4 to 2 (K=4) | - |
| **TMI** | VIS to DMN | 13 to 12 (K=19) | Positive (0.06) |
| | VIS to VIS | 6 to 6 (K=13, r=-0.05); 6 to 10 (K=13, r=0.05); 5 to 3 (K=17, r=0.06) | Overall positive (0.05 to 0.07) |
| | VIS to DAN | 17 to 9 (K=19) | Positive (0.06) |
| | FPN to VIS | 13 to 14 (K=15) | Positive (0.07) |

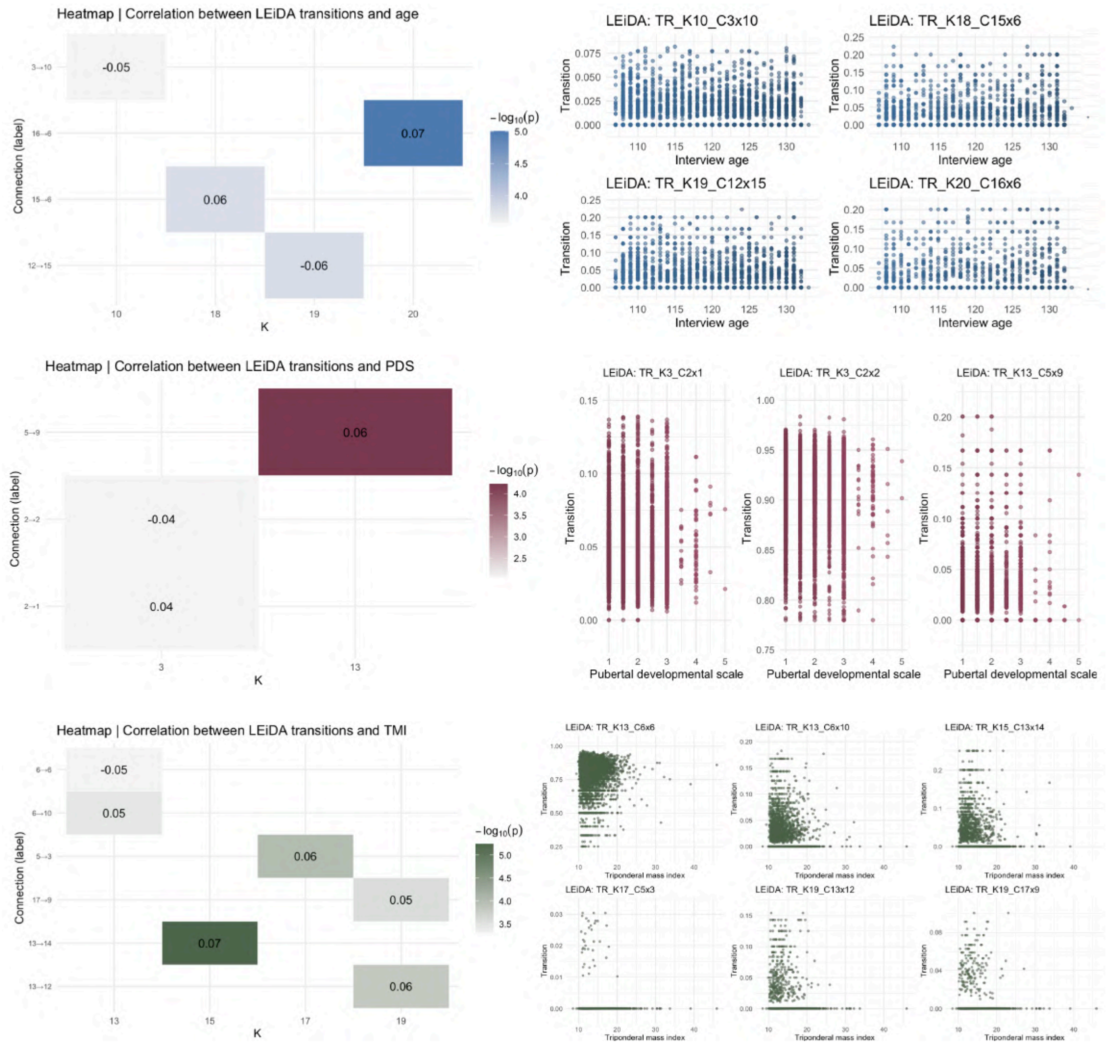

**Figure 34: Heatmaps and scatter plots between LEiDA transitions and continuous variables - original dataset.** Heatmap showing correlation values and significance (color scale) for LEiDA state transitions significantly associated with continuous variables (PDS, interview age and TMI) in the original dataset. Rows represent states, columns the K partitions. Under, scatter plots showing the distribution of LEiDA-derived state probabilities significantly associated with continuous variables (FDR-corrected $p < 0.05$). The x-axis represents covariates values. The y-axis indicates the probability of occurrence for each LEiDA state (N = 4,453). PDS = Pubertal Development Scale; TMI = Triponderal Mass Index.

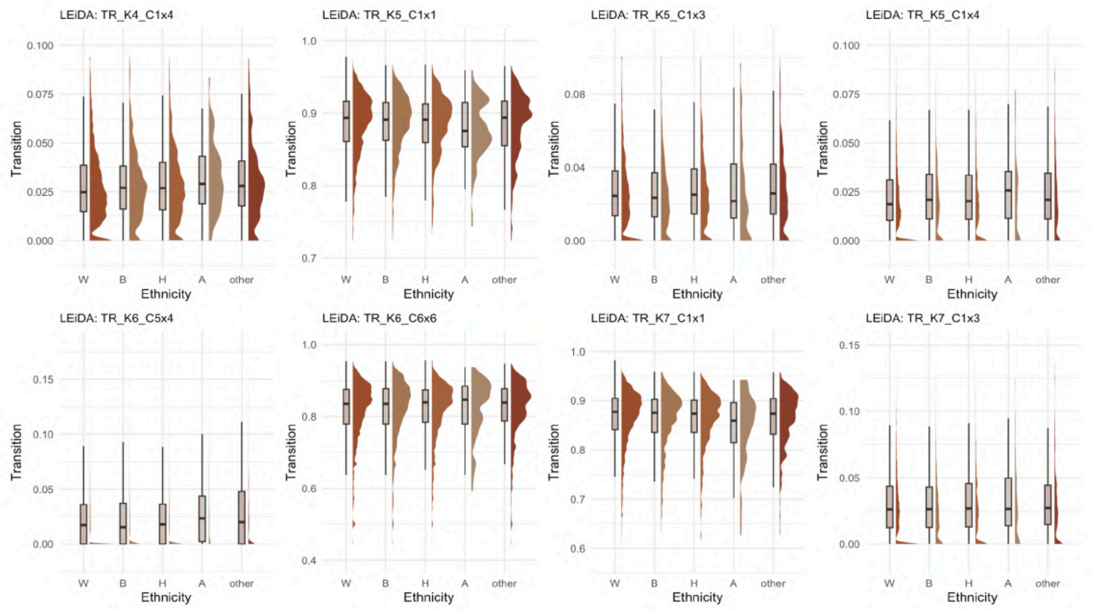

**Figure 35: Distribution of LEiDA state transitions across ethnicity groups - original dataset.** Half-violin plots showing the distribution of LEiDA-derived transitions significantly associated with ethnicity (FDR-corrected $p < 0.05$). The x-axis represents ethnicity groups: W = White, B = Black, H = Hispanic, A = Asian, other. The y-axis indicates the probability for each transition. Only states with significant associations are displayed (N = 4,453).

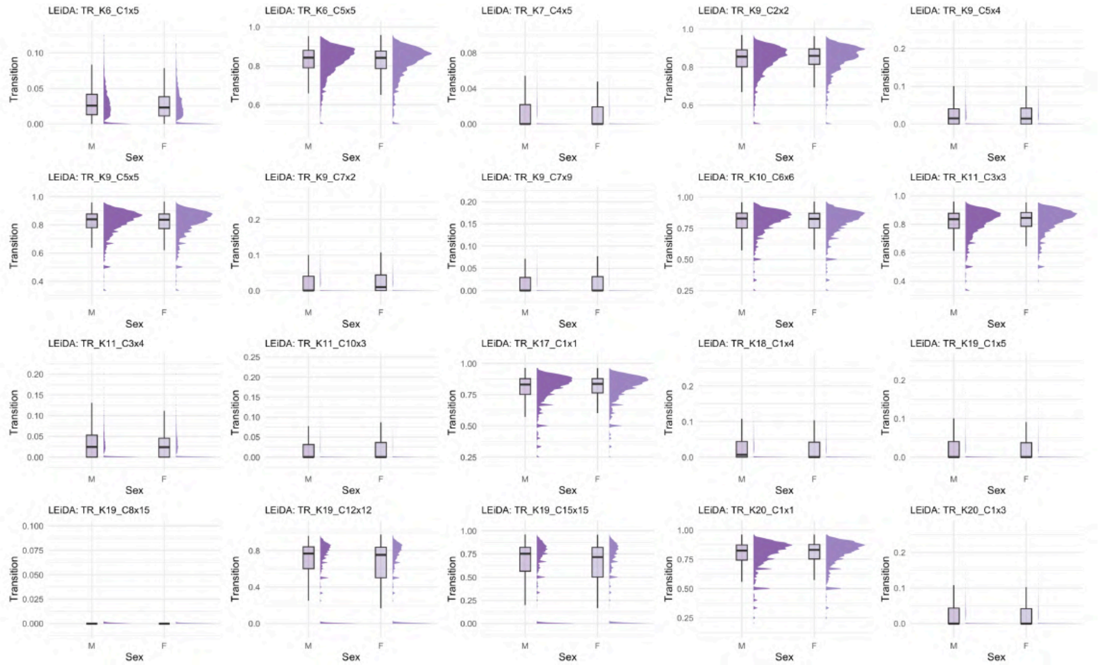

**Figure 36: Distribution of LEiDA state transitions across sex groups - original dataset.** Half-violin plots showing the distribution of LEiDA-derived state transitions significantly associated with sex (FDR-corrected $p < 0.05$). The x-axis represents sex groups: M = males, F = females. The y-axis indicates the probability for each LEiDA transition. Only states with significant associations are displayed (N = 4,453).

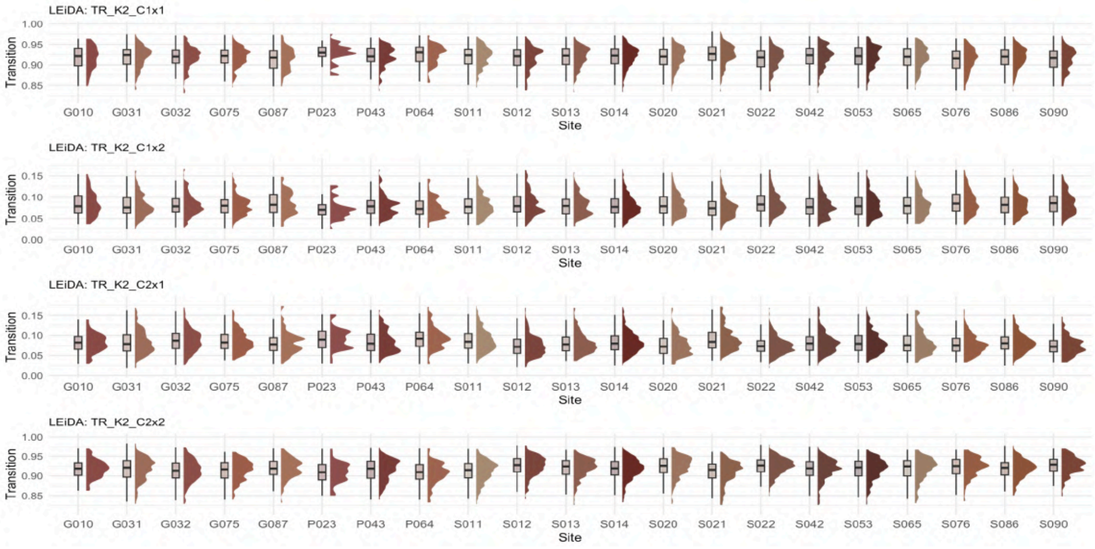

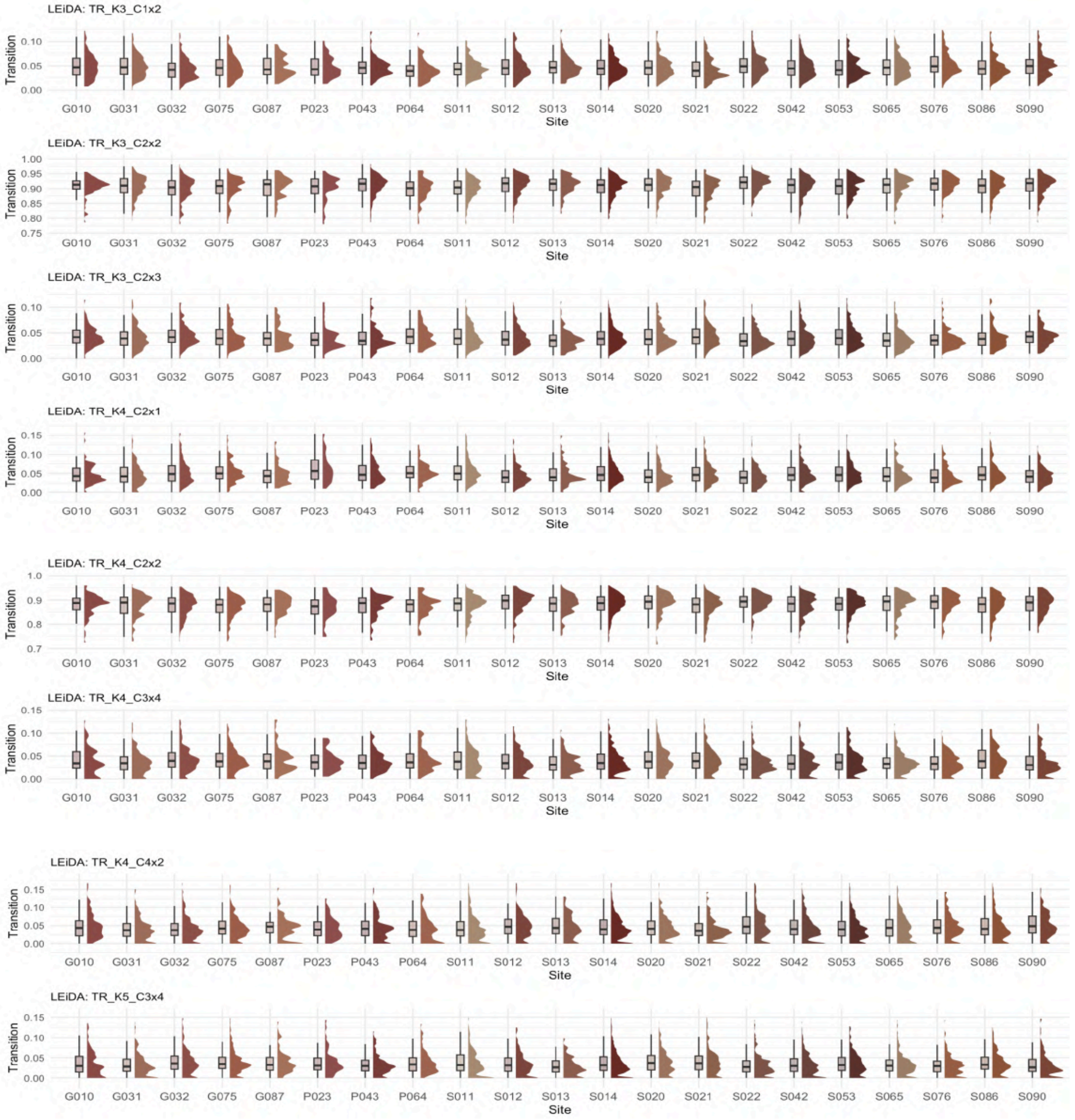

**Figure 37: Distribution of LEiDA state transitions across sites - original dataset.** Half-violin plots showing the distribution of LEiDA-derived transitions significantly associated with site (FDR-corrected $p < 0.05$). The x-axis represents different sites. The y-axis indicates the probability for each LEiDA transition. Only transitions with significant associations are displayed (N = 4,453).

The large number of LEiDA states and RSNs affected by site-related differences underscores that acquisition-specific variability can substantially influence LEiDA-derived metrics, highlighting the necessity of applying a harmonization procedure.

##### 2.5.3.2 Harmonized dataset

In this case, the variable site was excluded from these analyses, as harmonization rendered LEiDA-derived metrics entirely independent of acquisition site.

###### 2.5.3.2.1 Fractional occupancy (FO)

**Table 6: Detailed associations between Fractional Occupancy, states, and demographic variables - harmonized dataset.** Table providing a comprehensive overview of all significant associations between FO and demographic variables across all states and K values. Specific states, the corresponding K values, and the direction/range of correlations are included.

| Variable | Network(s) affected | States (K) | Effect (Pearson's r) |
|---|---|---|---|
| **Age** | Visual Network | 8 (K=11); 10 (K=10); 12 (K=12); 13 (K=16); 14 (K=15); 15 (K=17); 18 (K=20) | Negative (-0.04 to -0.06) |
| **Ethnicity** | DMN | 1 (K=5,7); 3 (K=19); 5 (K=18); 6 (K=15); 7 (K=10,14,16,17); 8 (K=9); 9 (K=11,13) | - |
| **PDS** | Visual Network | 7 (K=13); 8 (K=11); 9 (K=14); 13 (K=16) | Positive (0.04 to 0.05) |
| **Sex** | DMN | 1 (K=8,10,13,14,17–20); 2 (K=9,15,16); 3 (K=11); 4 (K=12,14); 5 (K=6–9,19); 6 (K=10); 7 (K=10,14); 8 (K=9); 9 (K=11,13); 10 (K=16); 11 (K=12,20); 12 (K=17–19) | - |
| | Visual | 4 (K=6,7); 12 (K=12) | - |
| | Ventral Attention | 4 (K=4); 10 (K=12); 11 (K=11,15,17); 12 (K=13,16) | - |
| | Frontoparietal | 11 (K=13,14); 14 (K=16); 15 (K=20) | - |
| **TMI** | - | - | - |

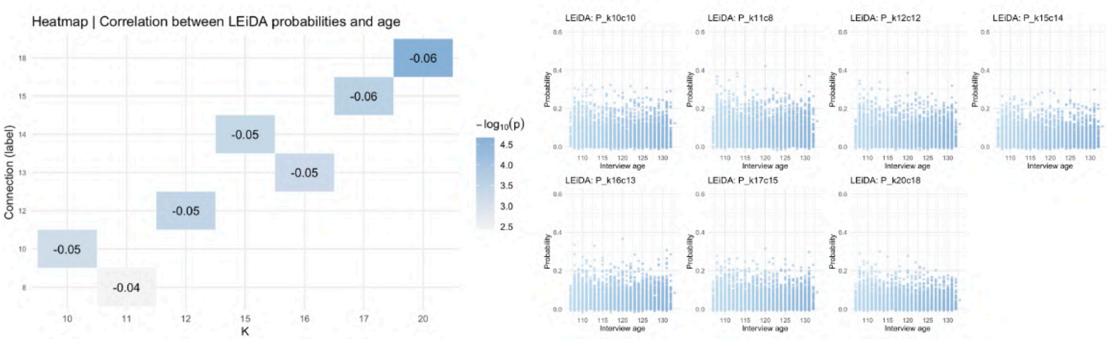

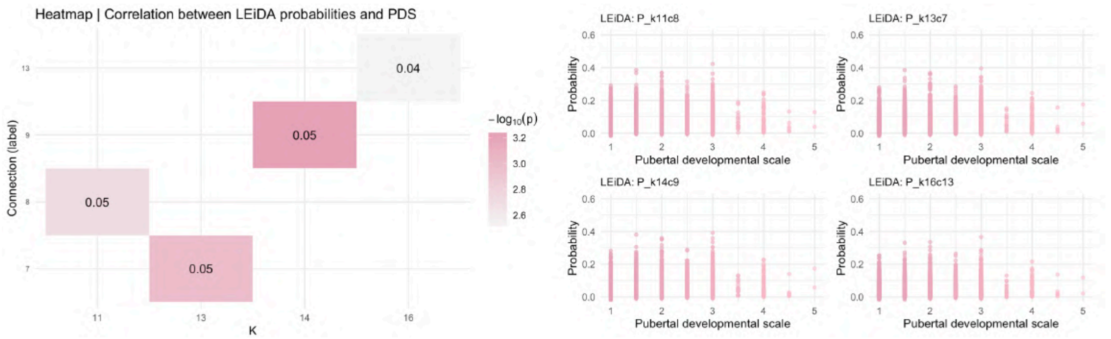

**Figure 38: Heatmaps and scatter plots between LEiDA FO and continuous variables - harmonized dataset.** Heatmap showing correlation values and significance (color scale) for LEiDA state FO significantly associated with continuous variables (PDS, interview age and TMI) in the harmonized dataset. Rows represent states, columns the K partitions. Under, scatter plots showing the distribution of LEiDA-derived state probabilities significantly associated with continuous variables (FDR-corrected $p < 0.05$). The x-axis represents covariates values. The y-axis indicates the probability of occurrence for each LEiDA state (N = 4,453). PDS = Pubertal Development Scale; TMI = Triponderal Mass Index.

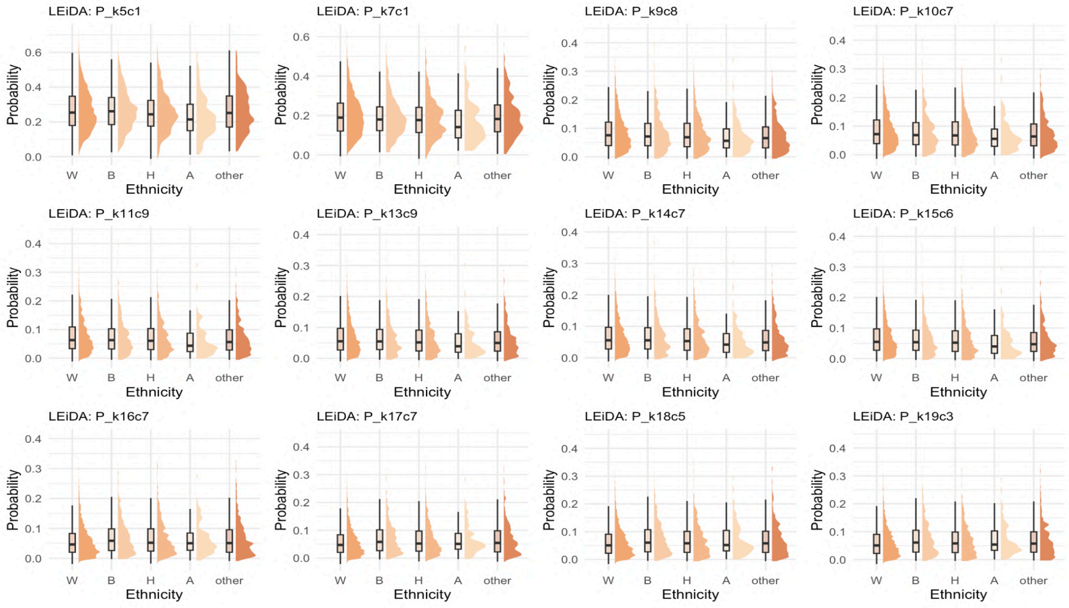

**Figure 39: Distribution of LEiDA state FO across ethnicity groups - harmonized dataset.** Half-violin plots showing the distribution of LEiDA-derived state FO significantly associated with ethnicity (FDR-corrected $p < 0.05$). The x-axis represents ethnicity groups: W = White, B = Black, H = Hispanic, A = Asian, other. The y-axis indicates the probability of occurrence for each state. Only states with significant associations are displayed (N = 4,453).

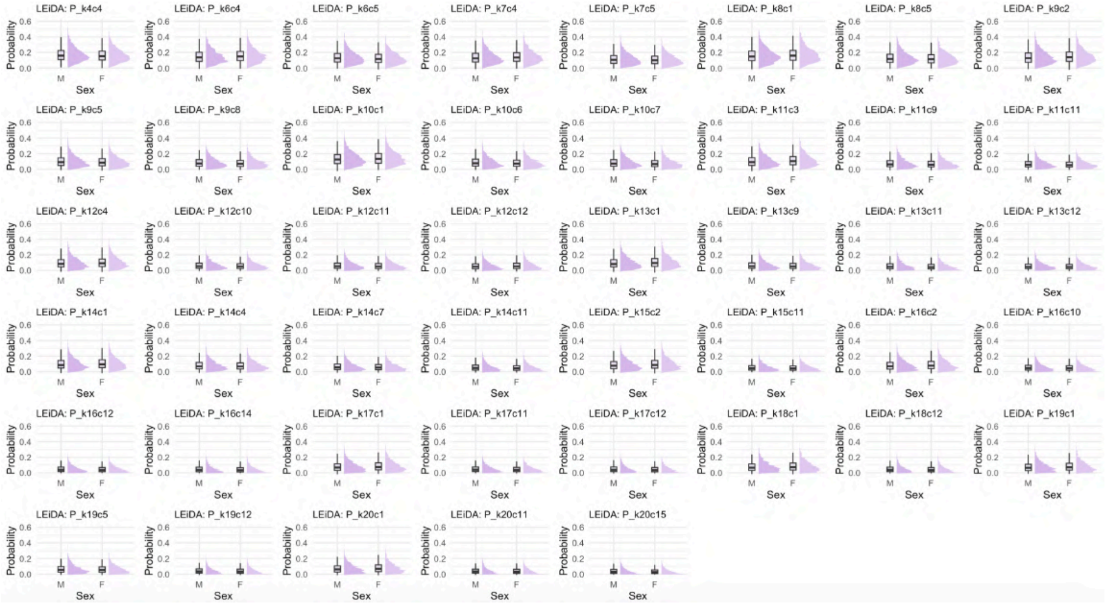

**Figure 40: Distribution of LEiDA state probabilities across sex groups - harmonized dataset.** Half-violin plots showing the distribution of LEiDA-derived state probabilities significantly associated with sex (FDR-corrected $p < 0.05$). The x-axis represents sex groups: M = males, F = females. The y-axis indicates the probability of occurrence for each state. Only states with significant associations are displayed (N = 4,453).

###### 2.5.3.2.2 Dwell Time (DT)

**Table 7: Detailed associations between Dwell Time, states, and demographic variables - harmonized dataset.** Table providing a comprehensive overview of all significant associations between DT and demographic variables across all states and K values. Specific states, the corresponding K values, and the direction/range of correlations are included.

| Variable | Network(s) affected | States (K) | Effect (Pearson's r) |
|---|---|---|---|
| **Age** | Visual Network | 10 (K=10); 12 (K=12); 13 (K=16); 14 (K=15); 18 (K=20) | Negative (-0.04 to -0.06) |
| **Ethnicity** | DMN | 1 (K=7); 6 (K=15); 8 (K=9) | - |
| **PDS** | Visual Network | 14 (K=14) | Negative (-0.05) |
| **Sex** | DMN | 1 (K=10,13,14,20); 2 (K=9,15); 3 (K=11); 5 (K=6,8) | - |
| | Visual Network | 2 (K=6); 4 (K=6,7) | - |
| | Dorsal Attention Network | 7 (K=8) | - |
| **TMI** | Visual | 2 (K=13); 6 (K=13,18); 7 (K=20) | Negative (-0.04 to -0.05) |
| **TMI** | Dorsal Attention | 6 (K=16) | Negative (-0.04) |
| **TMI** | Frontoparietal | 11 (K=13) | Negative (-0.04) |

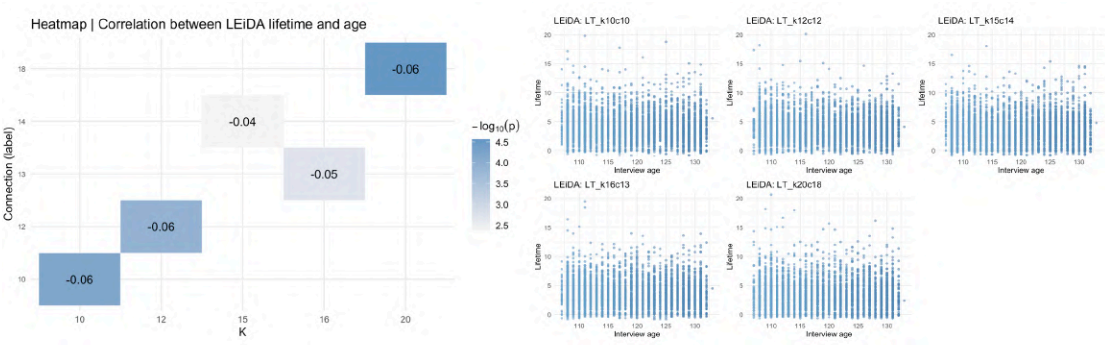

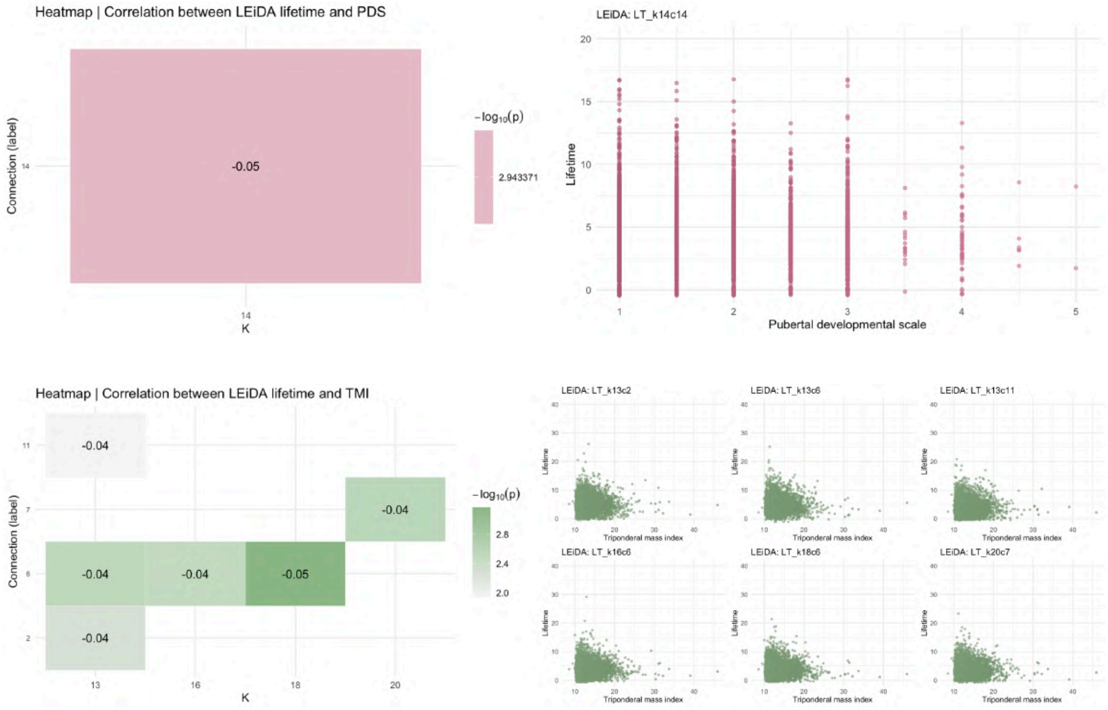

**Figure 41: Heatmaps and scatter plots between LEiDA DT and continuous variables - harmonized dataset.** Heatmap showing correlation values and significance (color scale) for LEiDA state DT significantly associated with continuous variables (PDS, interview age and TMI) in the harmonized dataset. Rows represent states, columns the K partitions. Under, scatter plots showing the distribution of LEiDA-derived state probabilities significantly associated with continuous variables (FDR-corrected $p < 0.05$). The x-axis represents covariates values. The y-axis indicates the probability of occurrence for each LEiDA state (N = 4,453). PDS = Pubertal Development Scale; TMI = Triponderal Mass Index.

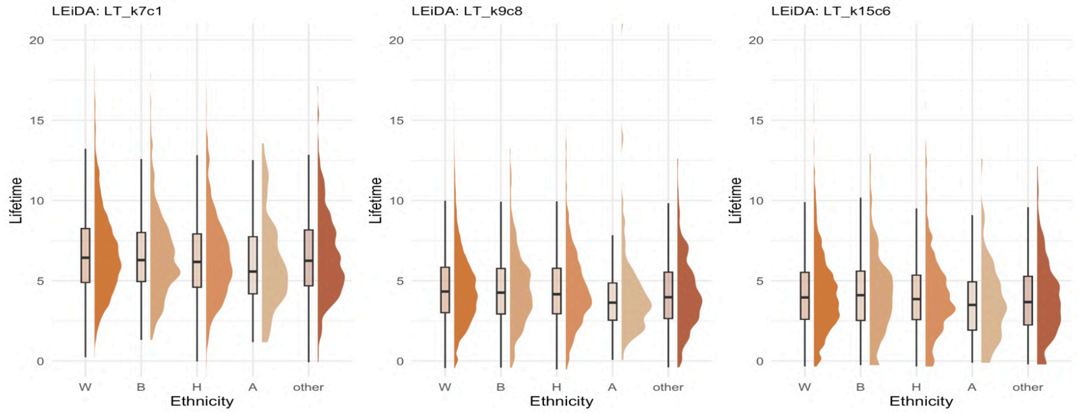

**Figure 42: Distribution of LEiDA state DT across ethnicity groups - harmonized dataset.** Half-violin plots showing the distribution of LEiDA-derived state DT significantly associated with ethnicity (FDR-corrected $p < 0.05$). The x-axis represents ethnicity groups: W = White, B = Black, H = Hispanic, A = Asian, other. The y-axis indicates the DT for each LEiDA state. Only states with significant associations are displayed (N = 4,453).

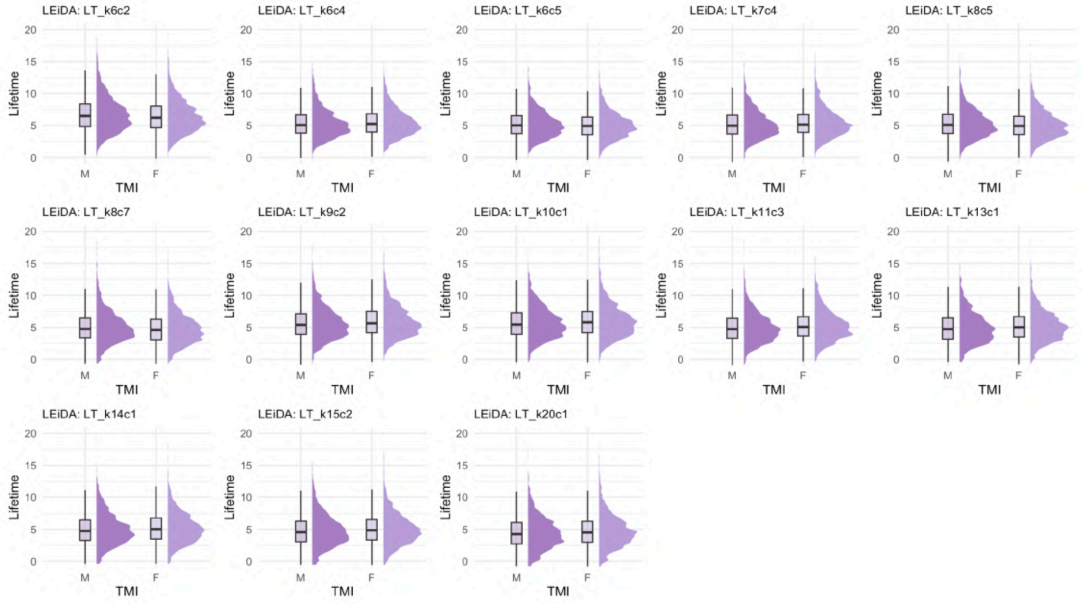

**Figure 43: Distribution of LEiDA state DT across sex groups - harmonized dataset.** Half-violin plots showing the distribution of LEiDA-derived state DT significantly associated with sex (FDR-corrected $p < 0.05$). The x-axis represents sex groups: M = males, F = females. The y-axis indicates the DT for each LEiDA state. Only states with significant associations are displayed (N = 4,453).

###### 2.5.3.2.3 Markov Chain transition probabilities

**Table 8: Detailed associations between transitions, states, and demographic variables - harmonized dataset.** Table providing a comprehensive overview of all significant associations between trajectories and demographic variables across all states and K values. Specific states, the corresponding K values, and the direction/range of correlations are included.

| Variable | Trajectories affected | States (K) | Effect (Pearson's r) |
|---|---|---|---|
| **Age** | DMN to VIS | 2 to 3 (K=16, r=0.07); 5 to 2 (K=18, r=-0.06); 1 to 13 (K=19, r=-0.06); 1 to 6 (K=20, r=-0.05); 4 to 2 (K=20, r=-0.07); 1 to 2 (K=20, r=0.09) | Negative (-0.05 to -0.09) and positive (0.07 to 0.10) |
| | DMN to VAN | 2 to 12 (K=16) | Negative (-0.06) |
| | DMN to FPN | 1 to 16 (K=19) | Negative (-0.05) |
| | VIS to DMN | 10 to 3 (K=19); 19 to 1 (K=20); 2 to 4 (K=20) | Negative (-0.05 to -0.09) |
| | VIS to VIS | 3 to 5 (K=17) | Positive (0.07 to 0.10) |
| | VIS to FPN | 5 to 8 (K=17) | Positive (0.10) |
| | FPN to VIS | 8 to 3 (K=18, r=0.06); 15 to 6 (K=20, r=-0.06) | Negative (-0.06) and positive (0.06) |
| **Ethnicity** | DMN to DMN | 1 to 1 (K=5) | - |
| | DMN to VIS | 1 to 3 (K=5); 5 to 4 (K=6); 2 to 3 (K=16); 1 to 6 (K=19); 1 to 2 (K=20) | - |
| | DMN to DAN | 4 to 16 (K=17) | - |
| | DMN to VAN | 1 to 4 (K=4) | - |
| | DMN to FPN | 10 to 14 (K=16); 1 to 16 (K=19) | - |
| | DMN to SMN | 1 to 15 (K=19) | - |
| | VIS to DMN | 2 to 1 (K=17); 3 to 4 (K=17); 2 to 3 (K=19) | - |
| | VIS to VIS | 4 to 5 (K=15); 3 to 5 (K=17); 7 to 4 (K=19); 9 to 18 (K=20) | - |
| | VIS to DAN | 2 to 10 (K=20) | - |
| | VIS to FPN | 14 to 8 (K=19); 7 to 8 (K=20) | - |
| | VAN to VIS | 16 to 6 (K=18) | - |
| | FPN to DMN | 15 to 1 (K=20) | - |
| | FPN to VIS | 8 to 3 (K=18); 8 to 4 (K=19) | - |
| | SMN to DMN | 15 to 1 (K=19) | - |
| | SMN to VIS | 15 to 7 (K=19) | - |
| **PDS** | DMN to VIS | 2 to 3 (K=16, r=0.09); 4 to 2 (K=20, r=0.11) | Positive (0.09 to 0.11) |

**Table 8: Detailed associations between transitions, states, and demographic variables - harmonized dataset.** Table providing a comprehensive overview of all significant associations between trajectories and demographic variables across all states and K values. Specific states, the corresponding K values, and the direction/range of correlations are included. *(continued)*

| Variable | Trajectories affected | States (K) | Effect (Pearson's r) |
|---|---|---|---|
| | DMN to FPN | 2 to 14 (K=16, r=0.05); 1 to 15 (K=20, r=0.06) | Positive (0.05 to 0.06) |
| | VIS to DMN | 3 to 7 (K=16, r=0.06); 19 to 1 (K=20, r=-0.05) | Positive (0.06) and negative (-0.05) |
| | VIS to VIS | 6 to 14 (K=19, r=-0.11); 6 to 9 (K=20, r=0.06) | Positive (0.06) and negative (-0.11) |
| | VIS to DAN | 2 to 10 (K=20) | Negative (-0.07) |
| | VIS to FPN | 5 to 8 (K=17) | Negative (-0.09) |
| | DAN to VIS | 6 to 4 (K=16, r=-0.05); 10 to 2 (K=20, r=0.07) | Positive (0.07) and negative (-0.05) |
| | DAN to VAN | 6 to 12 (k=16) | Negative (-0.05) |
| | FPN to VIS | 8 to 7 (K=20, r=0.05); 15 to 6 (K=20, r=-0.08) | Positive (0.05) and negative (-0.08) |
| **Sex** | DMN to DMN | 1 to 5 (K=6,7); 5 to 5 (K=6,7,9,19); 6 to 6 (K=10); 3 to 3 (K=11); 3 to 4 (K=11); 10 to 10 (K=16); 1 to 1 (K=17,20); 12 to 12 (K=19); 1 to 3 (K=20) | - |
| | DMN to VIS | 5 to 4 (K=9); 2 to 3 (K=16); 7 to 2 (K=17); 1 to 13 (K=19); 1 to 19 (K=20); 1 to 13 (K=20); 4 to 2 (K=20) | - |
| | DMN to VAN | 2 to 12 (K=16) | - |
| | DMN to FPN | 2 to 14 (K=16) | - |
| | VIS to DMN | 4 to 5 (K=7); 3 to 7 (K=16); 2 to 3 (K=19); 5 to 1 (K=20); 6 to 3 (K=20) | - |
| | VIS to VIS | 4 to 4 (K=7); 7 to 4 (K=19); 6 to 14 (K=19); 4 to 7 (K=19); 6 to 9 (K=20) | - |
| | VIS to DAN | 4 to 6 (K=16); 2 to 10 (K=20) | - |
| | VIS to FPN | 7 to 8 (K=19) | - |
| | DAN to DMN | 7 to 2 (K=9); 10 to 3 (K=11); 18 to 1 (K=19) | - |
| | VAN to VIS | 11 to 5 (K=17) | - |
| | FPN to VIS | 8 to 2 (K=20) | - |
| | SMN to SMN | 15 to 15 (K=19) | - |

**Table 8: Detailed associations between transitions, states, and demographic variables - harmonized dataset.** Table providing a comprehensive overview of all significant associations between trajectories and demographic variables across all states and K values. Specific states, the corresponding K values, and the direction/range of correlations are included. *(continued)*

| **Variable** | **Trajectories affected** | **States (K)** | **Effect (Pearson's r)** |
|---|---|---|---|
| **TMI** | DMN to VIS | 2 to 1 (K=15, r=-0.08); 2 to 3 (K=16, r=-0.10); 5 to 2 (K=18, r=0.09); 4 to 2 (K=20, r=0.07); 1 to 6 (K=20, r=0.08) | Positive (0.07 to 0.09) and negative (-0.08 to -0.10) |
| | VIS to DMN | 10 to 3 (K=19, r=-0.11); 18 to 4 (K=20, r=-0.06); 19 to 1 (K=20, r=0.11) | Positive (0.11) and negative (-0.06 to -0.11) |
| | VIS to VIS | 2 to 5 (K=14, r=-0.06); 5 to 3 (K=17, r=-0.08); 3 to 5 (K=17, r=-0.13); 6 to 3 (K=18, r=-0.08); 4 to 7 (K=19, r=0.05); 7 to 4 (K=19, r=-0.07); 5 to 7 (K=20, r=0.06); 2 to 10 (K=20, r=0.06) | Positive (0.05 to 0.06) and negative (-0.06 to -0.13) |
| | VIS to FPN | 5 to 8 (K=17, r=0.06); 6 to 8 (K=18, r=-0.07); 6 to 16 (K=19, r=-0.06); 7 to 8 (K=19, r=-0.08); 7 to 8 (K=20, r=0.05); 2 to 8 (K=20, r=-0.06) | Positive (0.05 to 0.06) and negative (-0.06 to -0.09) |
| | DAN to VIS | 6 to 4 (K=16) | Negative (-0.05) |
| | FPN to DMN | 15 to 1 (K=20) | Negative (-0.07) |
| | FPN to VIS | 15 to 2 (K=20, r=0.05); 8 to 2 (K=20, r=-0.05) | Positive (0.05) and negative (-0.05) |
| | SMN to DMN | 15 to 1 (K=19) | Negative (-0.05) |

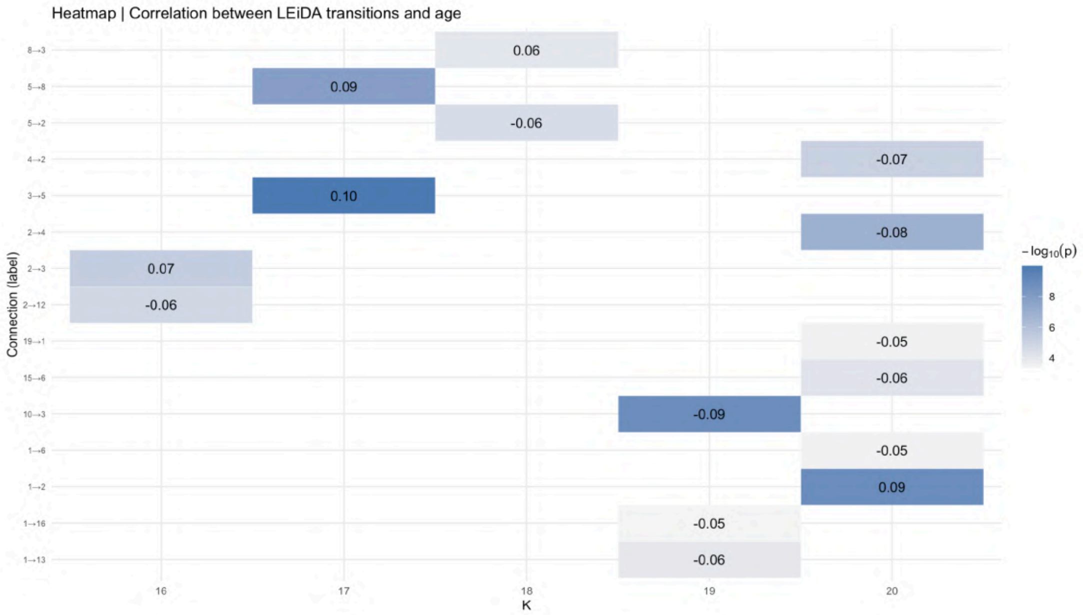
Heatmap | Correlation between LEiDA transitions and age
Connection (label)
K
16
17
18
19
20
0.06
0.09
-0.06
-0.07
0.10
-0.08
0.07
-0.06
-0.05
-0.06
-0.09
-0.05
0.09
-0.05
-0.06
$-\log_{10}(p)$
8
6
4

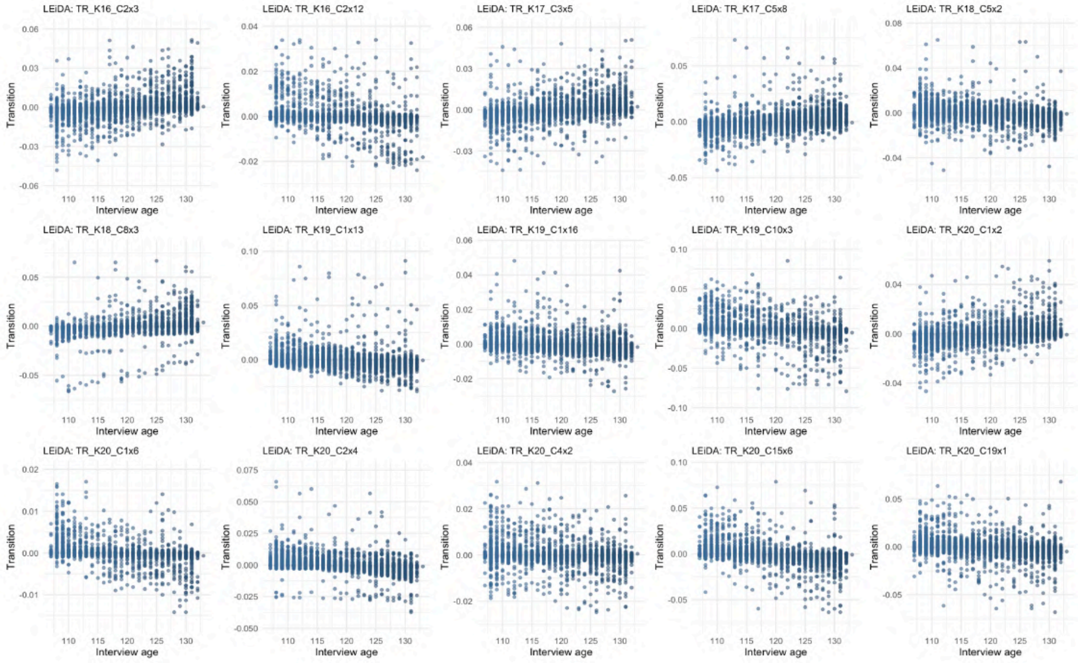
LEiDA: TR_K16_C2x3
LEiDA: TR_K16_C2x12
LEiDA: TR_K17_C3x5
LEiDA: TR_K17_C5x8
LEiDA: TR_K18_C5x2
LEiDA: TR_K18_C8x3
LEiDA: TR_K19_C1x13
LEiDA: TR_K19_C1x16
LEiDA: TR_K19_C10x3
LEiDA: TR_K20_C1x2
LEiDA: TR_K20_C1x6
LEiDA: TR_K20_C2x4
LEiDA: TR_K20_C4x2
LEiDA: TR_K20_C15x6
LEiDA: TR_K20_C19x1
Transition
Interview age

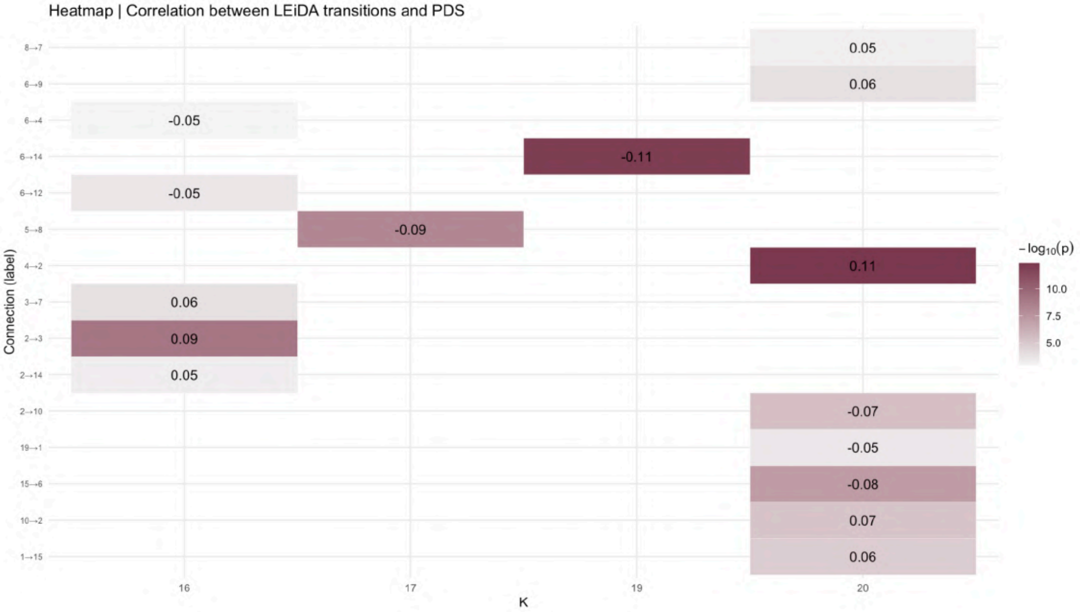
Heatmap | Correlation between LEiDA transitions and PDS
8→7
6→9
6→4
6→14
6→12
5→8
4→2
3→7
2→3
2→14
2→10
19→1
15→6
10→2
1→15
Connection (label)
0.05
0.06
-0.05
-0.11
-0.05
-0.09
0.11
0.06
0.09
0.05
-0.07
-0.05
-0.08
0.07
0.06
16
17
19
20
K
$-\log_{10}(p)$
10.0
7.5
5.0

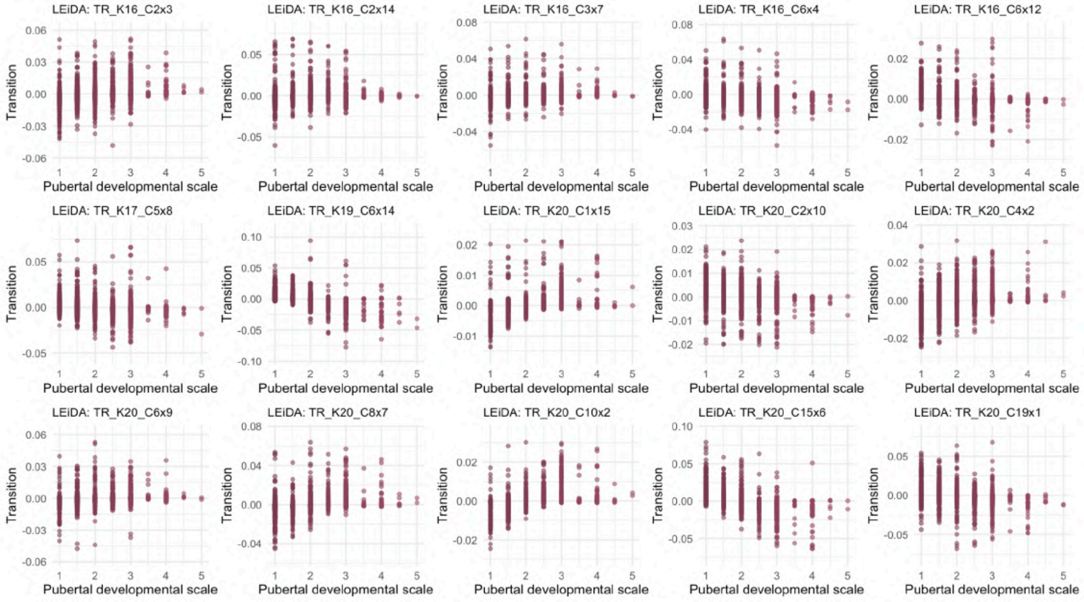
LEiDA: TR_K16_C2x3
LEiDA: TR_K16_C2x14
LEiDA: TR_K16_C3x7
LEiDA: TR_K16_C6x4
LEiDA: TR_K16_C6x12
LEiDA: TR_K17_C5x8
LEiDA: TR_K19_C6x14
LEiDA: TR_K20_C1x15
LEiDA: TR_K20_C2x10
LEiDA: TR_K20_C4x2
LEiDA: TR_K20_C6x9
LEiDA: TR_K20_C8x7
LEiDA: TR_K20_C10x2
LEiDA: TR_K20_C15x6
LEiDA: TR_K20_C19x1
Transition
Pubertal developmental scale

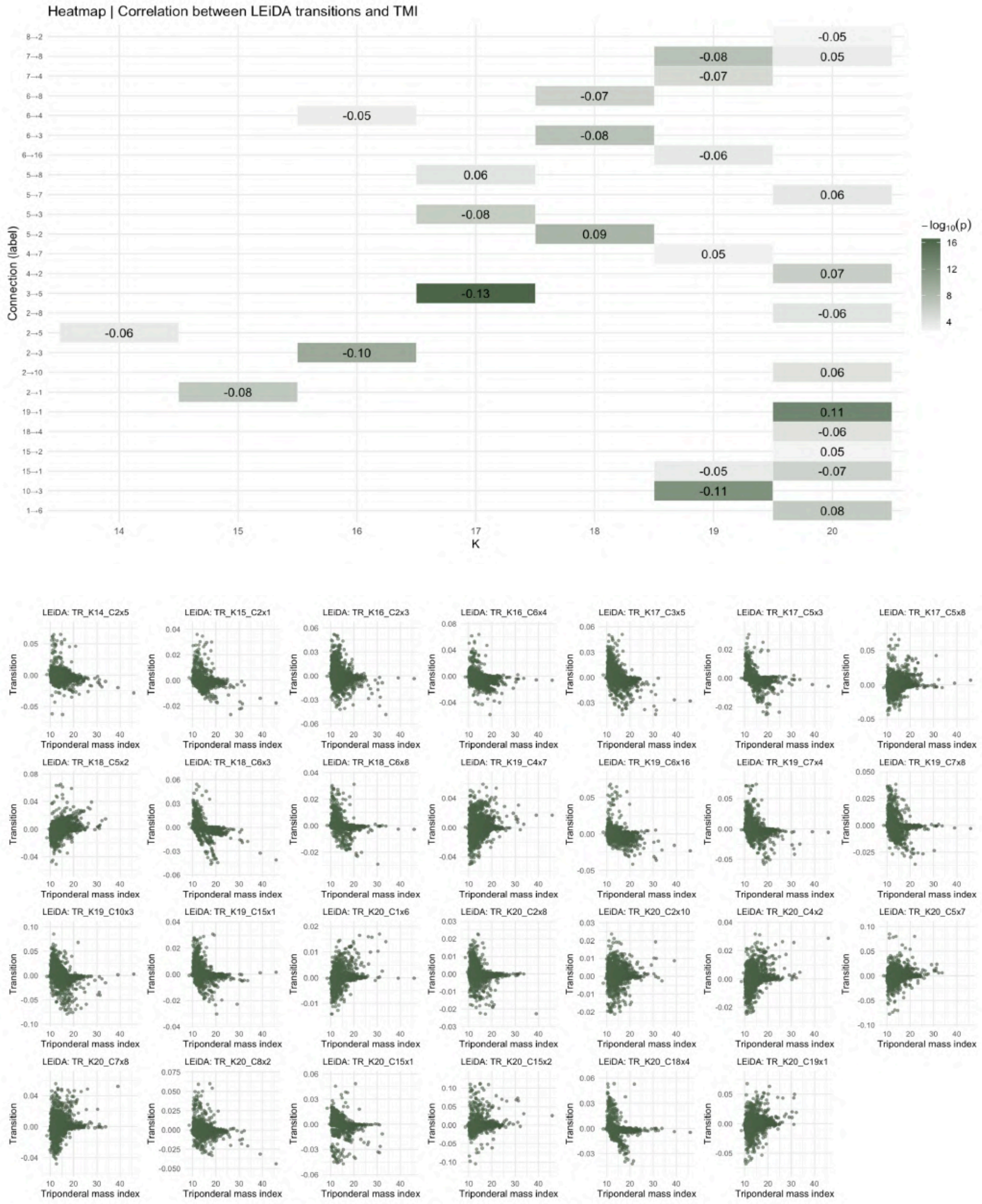

**Figure 44: Heatmaps and scatter plots between LEiDA transitions and continuous variables - harmonized dataset.** Heatmap showing correlation values and significance (color scale) for LEiDA state transitions significantly associated with continuous variables (PDS, interview age and TMI) in the harmonized dataset. Rows represent states, columns the K partitions. Under, scatter plots showing the distribution of LEiDA-derived state probabilities significantly associated with continuous variables (FDR-corrected $p < 0.05$). The x-axis represents covariates values. The y-axis indicates the probability of occurrence for each LEiDA state (N = 4,453). PDS = Pubertal Development Scale; TMI = Triponderal Mass Index.

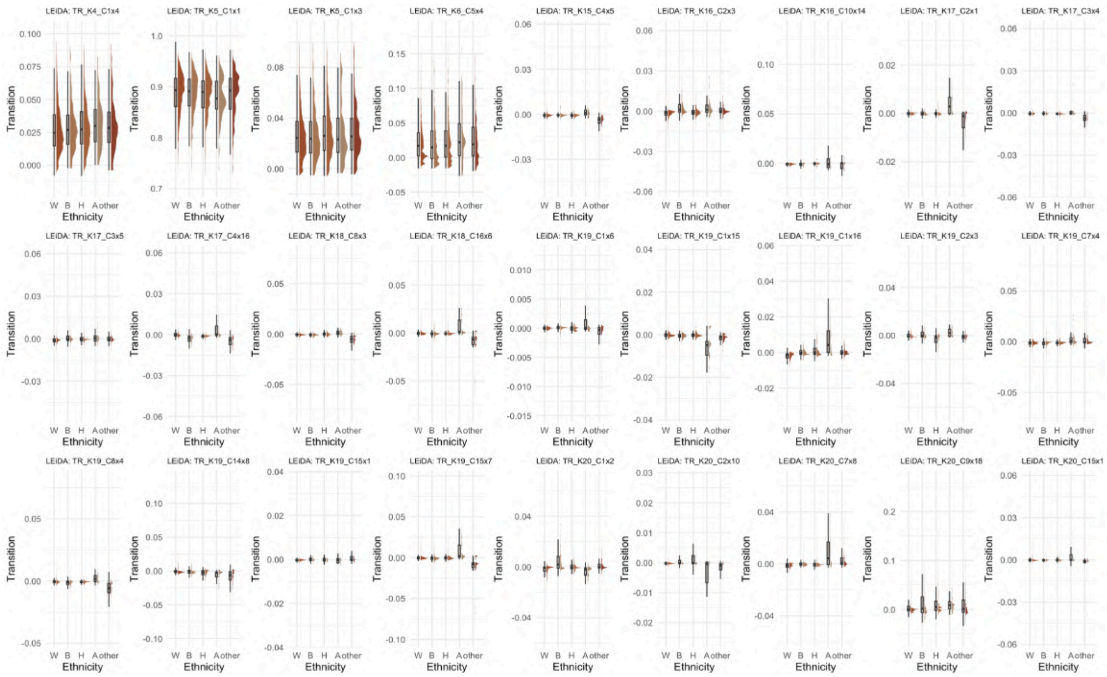

**Figure 45: Distribution of LEiDA state transitions across ethnicity groups - harmonized dataset.** Half-violin plots showing the distribution of LEiDA-derived state transitions significantly associated with ethnicity (FDR-corrected $p < 0.05$). The x-axis represents ethnicity groups: W = White, B = Black, H = Hispanic, A = Asian, other. The y-axis indicates the probability for each LEiDA transition. Only states with significant associations are displayed (N = 4,453).

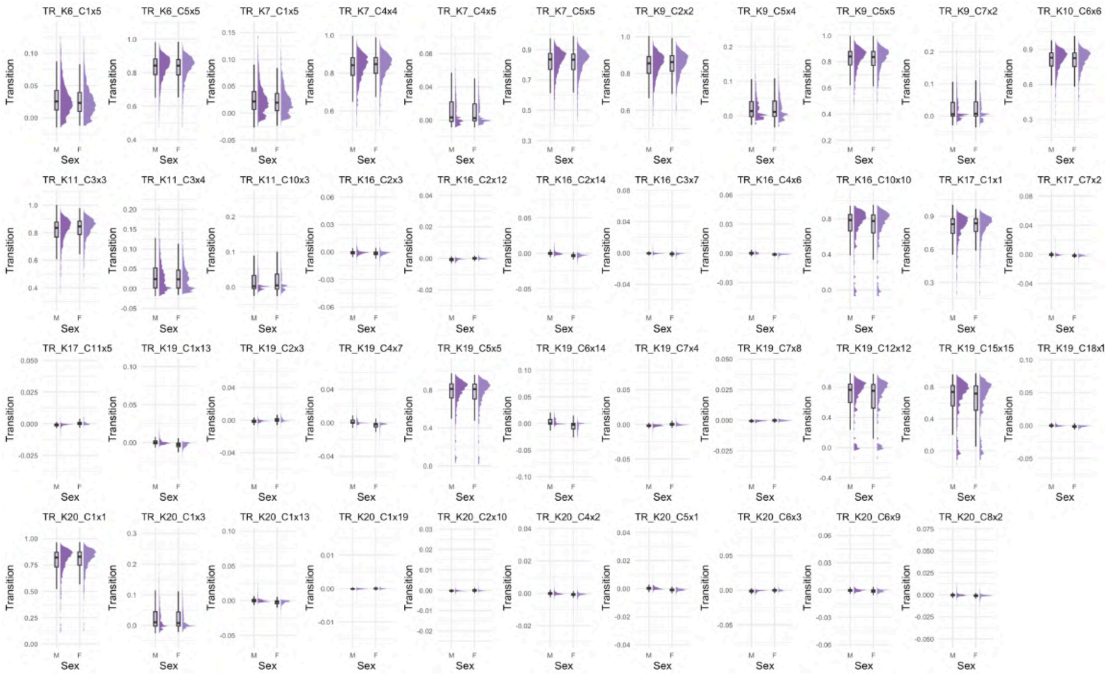

**Figure 46: Distribution of LEiDA state transitions across sex groups - harmonized dataset.** Half-violin plots showing the distribution of LEiDA-derived state transitions significantly associated with sex (FDR-corrected $p < 0.05$). The x-axis represents sex groups: M = males, F = females. The y-axis indicates the probability for each LEiDA transition. Only states with significant associations are displayed (N = 4,453).

For both datasets, it is important to note the distribution characteristics of LEiDA-derived transition probabilities. In many cases, these values are heavily concentrated near zero, reflecting the low likelihood of transitions between certain network pairs. As a result, some distribution plots appear tightly clustered around zero, making visual differences between groups difficult to discern. Additionally, several variables exhibit a high proportion of outliers or skewed distributions, which increases the range displayed on the y-axis. These patterns arise naturally from inter-individual variability, particularly in transitions involving non-dominant networks. Nevertheless, we present results for all transitions that showed significant associations with the demographic variables under study.

In general, transitions within the same RSN (e.g., DMN to DMN or Dorsal Attention to Dorsal Attention) exhibit higher probabilities, with median values around 0.9. In contrast, transitions between different networks typically show much lower probabilities: for example, DMN to Visual Network transitions have a median around 0.03, and DMN to Ventral Attention transitions around 0.02.

Interestingly, also transitions within the same RSN but represented by different LEiDA centroids show lower probabilities than those within the same RSN and the same centroid. This highlights subtle distinctions captured by LEiDA during clustering, reflecting how different centroids can represent slightly divergent dynamic patterns within the same network.

#### 2.6 Elastic Net analysis

To maintain consistency with previous studies investigating the relationship between various MRI features (sMRI, DTI, rs-fMRI) and TMI, we applied an ElasticNet regression model to predict TMI based on covariates with added: rs-fMRI features, LEiDA-derived metrics, FCH-derived measures, and the combination of all imaging features respectively.

This analysis was conducted on both original and harmonized datasets, whereas the following results are referred to the original dataset.

**Table 9: Model performance for predicting triponderal mass index from demographic, developmental and brain-based features - original dataset.** Performance of elastic net regression models trained to predict triponderal mass index (TMI) using different feature sets in the original dataset (N = 4,453). Covariates; demographic covariates only, Rs-fMRI; whole-brain resting-state fMRI connectivity, LEiDA; LEiDA dynamic-state metrics, FCH; FCH harmonic coefficients, All; all features combined. Metrics reported include the coefficient of determination (R2) and negative mean absolute error (negMAE), averaged across 5 cross-validation folds, with standard deviation (STD) shown for both training and held-out test sets.

| Model | Metric | Training – mean | Training – STD | Test – mean | Test – STD |
|---|---|---|---|---|---|
| Covariates | *R2* | 0.1045 | 0.0025 | 0.0865 | 0.0084 |
| Covariates | *negMAE* | -1.7577 | 0.0119 | -1.7726 | 0.0308 |
| rs-fMRI | *R2* | 0.1659 | 0.0175 | 0.1081 | 0.0117 |
| rs-fMRI | *negMAE* | -1.6970 | 0.0243 | -1.7497 | 0.0216 |
| LEiDA | *R2* | 0.0969 | 0.0048 | 0.0707 | 0.0054 |
| LEiDA | *negMAE* | -1.7782 | 0.0122 | -1.7939 | 0.0264 |
| FCH | *R2* | 0.1103 | 0.0045 | 0.0841 | 0.0095 |
| FCH | *negMAE* | -1.7526 | 0.0126 | -1.7752 | 0.0288 |
| All | *R2* | 0.1437 | 0.0156 | 0.0898 | 0.0063 |
| All | *negMAE* | -1.7297 | 0.0199 | -1.7729 | 0.0273 |
